# Enzyme-free synthesis of natural phospholipids in water

**DOI:** 10.1101/746156

**Authors:** Luping Liu, Yike Zou, Ahanjit Bhattacharya, Dongyang Zhang, Susan Q. Lang, K. N. Houk, Neal K. Devaraj

## Abstract

All living organisms synthesize phospholipids as the primary constituent of their cell membranes. While phospholipids can spontaneously self-assemble in water to form membrane-bound vesicles, their aqueous synthesis requires pre-existing membrane-embedded enzymes. This limitation has led to models in which the first cells used simpler types of membrane building blocks and has hampered integration of phospholipid synthesis into artificial cells. Here we demonstrate that a combination of ion pairing and self-assembly of reactants allows high-yielding synthesis of cellular phospholipids in water. Acylation of 2-lysophospholipids using cationic thioesters occurs in mildly alkaline solutions resulting in the formation of cell-like membranes. A variety of membrane-forming natural phospholipids can be synthesized. Membrane formation takes place in water from natural alkaline sources, such as soda lakes and hydrothermal oceanic vents. When formed vesicles are transferred to more acidic solutions, electrochemical proton gradients are spontaneously established and maintained.

## Main Text

Cellular membranes composed of glycerophospholipids are found in all living organisms (*1*). Bacterial and eukaryotic membranes consist of diacylphospholipids, in which two ester linkages connect a polar head group to two hydrophobic tails. Cells synthesize diacylphospholipids through enzymatic acylation of lysophospholipids (*2*). As several enzymes involved in phospholipid biosynthesis must themselves be membrane-bound for enzyme activity, it is unclear if phospholipid membranes could have formed in the absence of advanced enzymatic machinery (*3*–*6*). Enzyme-free synthesis of glycerophospholipids can occur using wet-dry cycling (*7*, *8*) and acylation of glycerophosphates can take place in the presence of a large excess of activated long-chain acyl imidazole derivatives, but the reactions require organic co-solvent due to the hydrophobicity of the acylating precursors and result in unnatural phospholipids (*9*, *10*). A high-yielding synthesis of natural phospholipids in water would shed light on the origin and evolution of cellular membranes and open up new routes for lipid synthesis in artificial cells (*11*, *12*).

In cells, lysophospholipid esterification requires acyl thioesters and the action of an acyltransferase. Inspired by acyltransferase reactions in lipid biosynthesis (*2*), and past hypotheses on the role of thioesters in the origins of life (*13*), we asked if synthetic acyl thioesters **2** in lieu of acyl coenzyme A, could acylate 1-acyl-2-hydroxy-*sn*-glycero-3-phosphocholines **1** in water to give the desired natural phosphatidylcholines (PC) **3** (Fig. 1A). An ionized polar head group on **2** would facilitate transacylation by increasing the long-chain acyl thioester’s solubility in water and driving the self-assembly of micelles, possibly in combination with the lysophospholipid reactant. We initially investigated the synthesis of a naturally occurring phospholipid, 1,2-dioleoyl-*sn*-glycero-3-phosphocholine **3a** (DOPC) (*14*), in aqueous solution via acylation of lysophospholipid 1-oleoyl-2-hydroxy-*sn*-glycero-3-phosphocholine **1a** (Fig. 1A). A water-soluble anionic thioester, sodium 2-(oleoylthio)ethane-1-sulfonate **2a**, was chosen as an oleoyl donor (*15*). Only trace conversion was observed (<1%) and addition of acylation catalysts to accelerate the transacylation reaction did not have a substantial effect (Table S1). However, to enable spontaneous membrane formation in water, acylation would have to be high-yielding and occur readily at near stoichiometric ratios of starting materials, to prevent amphiphilic precursors from disrupting membrane formation.

**Fig. 1.**
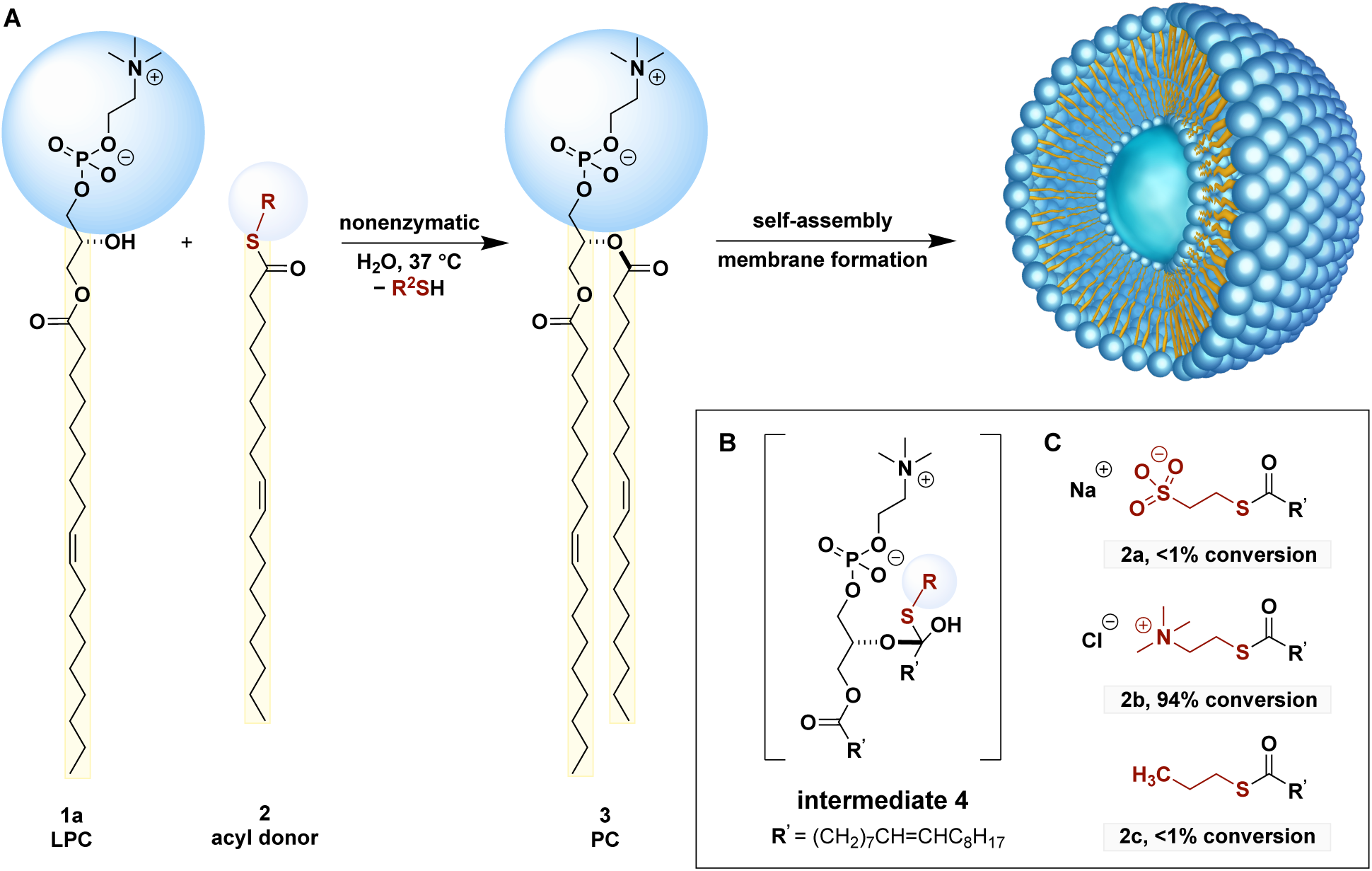
Nonenzymatic synthesis of natural phospholipids. (**A**) *De novo* synthesis of membrane-forming phospholipids in water. LPC: lysophosphatidylcholine. PC, phosphatidylcholine. (**B**) Proposed reaction intermediate. (**C**) Reactive fatty acyl derivatives: acyl donors **2**, with which reactions were carried out with 1-oleoyl-2-hydroxy-*sn*-glycero-3-phosphocholine **1a** (0.5 mM) and acylation reagents (0.75 mM) in the presence of Na_2_CO_3_/NaHCO_3_ buffer (pH = 10.6) at 37 °C for 5 hours.

To improve the yield of the desired phospholipid, we sought a way to increase the rate of transacylation while minimizing the hydrolysis of the acyl donor. Transacylation reactions between acyl donors and alcohols proceed via a tetrahedral intermediate (Fig. 1B) (*16*, *17*). We reasoned that the observed poor reactivity in the transacylation reaction is likely due to the inability of acylation reagent **2a** to lower the *Δ*G for the tetrahedral intermediate. We therefore sought to synthesize an acyl donor that would stabilize the reaction intermediate. In principle, a positively charged leaving group on the acyl donor **2** would lead to a favorable Coulombic interaction (*18*) with the negatively charged phosphate group of lysophospholipid **1** (Fig. 1B), thus stabilizing intermediate **4**, and accelerating the anticipated transacylation reaction through ion pairing. Based on this hypothesis, we prepared 2-(oleoylthio)-*N*,*N*,*N*-trimethylethan-1-aminium chloride **2b**, containing a positively charged quaternary amine head group (Fig. 1C). Precursor **2b** is accessible in one synthetic step from oleic acid and thiocholine. **2b** could also be obtained using prebiotically relevant condensing agents such as dicyandiamide (Fig. S1). (*19*) We evaluated the esterification reaction of lysophospholipid **1a** (0.5 mM) with **2b** (0.75 mM) in alkaline bicarbonate buffer (pH = 10.6) in the absence of additional catalyst. 94% conversion of lysophospholipid **1a** to DOPC **3a** was obtained after 5 hours at 37 °C (Fig. 1C). In contrast, under the same reaction conditions, we observed only trace acylation of **1a** with either anionic **2a** or the charge neutral acylation reagent **2c** (<1% conversion).

Our initial results suggested that ion pairing is involved in the aqueous transacylation of 2-lysophospholipids. To better understand the effect of ion pairing on lysophospholipid acylation, density functional theory (DFT) calculations were performed (*20*). The *n*-butyroyl group was employed in place of the oleoyl group as a model system (Fig. 2). Calculations were performed at D3-B3LYP/6-311++G(2d,p)//B3LYP/6-31G (d,p) level with the Cramer-Truhlar polarizable continuum model (Solvation Model Density, SMD) for water (*21*). *Δ*G for the formation of the tetrahedral intermediate **INT1** from model acylation reagent 2-(butyroylthio)ethane-1-sulfonate was calculated as 24.1 kcal/mol, while for **INT2** from 2-(butyroylthio)-*N*,*N*,*N*-trimethylethan-1-aminium it was 16.5 kcal/mol (Fig. 2A). These computational results predict a higher reactivity of the quaternary ammonium-containing reagent analogous to **2b** in the transacylation reaction, in line with our experimental observations. Further, the optimized structures of the reaction intermediates indicate that intramolecular Coulombic interactions are important for the stability of the tetrahedral intermediate (Fig. 2B). In **INT1**, the sulfonate side chain is distal to the phosphate group, which results from the electrostatic repulsion of these two negatively charged groups. In contrast, the optimized structure of **INT2** shows electrostatic attraction between the negatively charged phosphonate group and the two positively charged quaternary ammonium groups. Our experimental and theoretical findings are consistent with the hypothesis that a positively charged head group on **2** stabilizes the reaction intermediate, resulting in efficient transacylation.

**Fig. 2.**
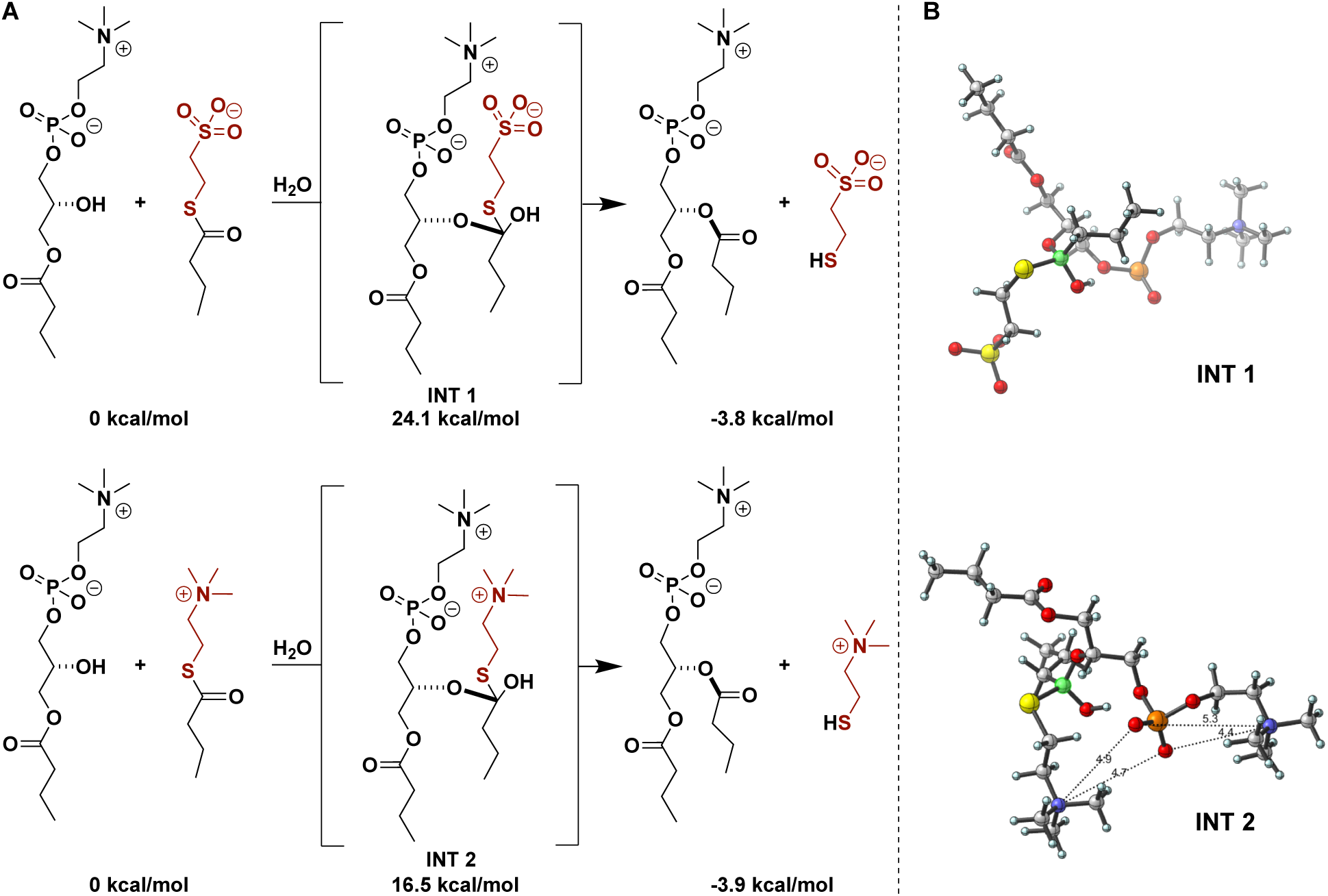
Predicted effects of thioester charge on phospholipid synthesis. (**A**) DFT-calculated transacylation reactions. (**B**) Optimized structures of reaction intermediates.

We next explored the range of reaction conditions under which phospholipid could be synthesized. The transacylation reaction between lysophospholipid **1a** and acyl donor **2b** proceeded rapidly and in high yield in alkaline bicarbonate buffers (pH 9.5–10.6) (Fig. 3A) (*22*) In addition, over a longer time period, good reaction yields were obtained in Na_2_CO_3_/NaHCO_3_ buffer at pH = 8.8 (63% yield, 24 hours, entry 2). In contrast no desired product was obtained using phosphate buffered saline (PBS) at pH = 7.4 (<1% yield, 24 hours, entry 1). Based on these results, we tested if the reaction could proceed in more complex naturally derived alkaline water samples. Numerous models have proposed that life may have started near alkaline hydrothermal vents or soda lakes. (*23*, *24*) To test if synthesis of phospholipids could take place in such environments, we obtained water samples from the Lost City Hydrothermal Field (LCHF, pH = 9.1) and Mono Lake (pH = 10), a Californian soda lake. Acylation of **1a** with **2b** to form phospholipid **3a** took place in water from the LCHF (42% yield, 48 hours, entry 3) and Mono Lake water (76% yield, 5 hours, entry 5). These results suggest that natural alkaline water sources would have been privileged environments for thioester mediated acylation of lysophospholipids.

**Fig. 3.**
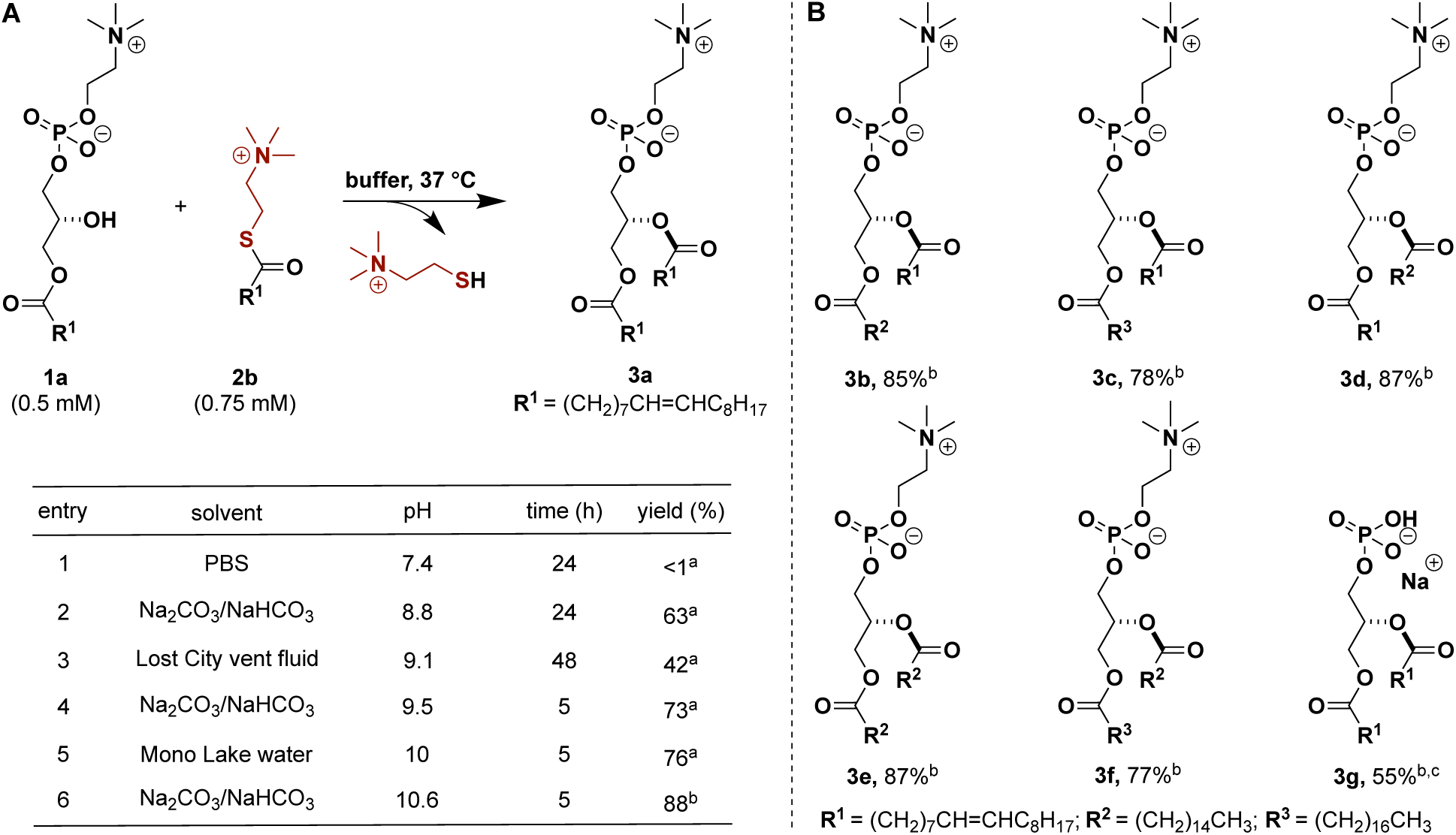
Ion pairing-enabled natural phospholipid synthesis. (**A**) Synthesis of phospholipid **3a** using different water sources. (**B**) The scope of synthesizing natural phospholipids. Reactions were carried out with lysophospholipids (0.5 mM) and acylation reagents (0.75 mM) in buffer at 37 °C. PBS, phosphate buffered saline. ^a^HPLC yield. ^b^Isolated yield after reactions were performed in Na_2_CO_3_/NaHCO_3_ buffer (pH = 10.6) for 5 hours. ^c^1.5 mM of reagent **2b** was used.

Having optimized reaction conditions (entry 6, Fig. 3A), we investigated the generality of the technique by attempting to synthesize a variety of naturally occurring phospholipids. We explored the acylation of multiple lysophosphatidylcholines with acylation agents of different chain length and composition. Good acylation yields were achieved when **2b** was reacted with lysophospholipids possessing either unsaturated (oleoyl) or saturated (palmitoyl and stearoyl) acyl chains (**3b** and **3c**) (Fig. 3B). Changing the acyl donor to a saturated palmitoyl group through the use of 2-(palmitoylthio)-*N*,*N*,*N*-trimethylethan-1-aminium chloride **2d**, led to similar reactivities (**3d**–**3f**), and enabled the synthesis of dipalmitoylphosphatidylcholine, a key component of pulmonary surfactant (*25*).We also tested if we could synthesize phosphatidic acids, as they are universal intermediates in glycerophospholipid biosynthesis. (*26*) Acylation of the sodium salt of 1-oleoyl-2-hydroxy-sn-glycero-3-phosphate with **2b**, led to a 55% yield of phosphatidic acid **3g**, demonstrating that head group substitution is tolerated.

Both the lysophospholipids and acylation agents self-assemble to form micelles under the optimized reaction conditions (Fig. S2). The reaction may occur specifically within mixed micelles, possibly with preorganization of the amphiphilic reagents improving reaction rate and selectivity. The transacylation reaction is chemoselective, being unperturbed by the presence of non-amphiphilic biomolecules possessing nucleophilic functional groups. Under the optimized reaction conditions, no oleoylation products were observed when combining acylation reagent **2b** with either serine, threonine or glycerophosphocholine as substrates (Fig. S3–S5). We also observed <1% of product when the non-amphiphilic acyl donor acetylthiocholine chloride was used for the acylation of **1a** (Fig. S6). Shorter single-chain lipid precursors are more prebiotically plausible reactants, but compared to longer chain precursors, require higher concentrations to form micelles. Indeed, when we reacted the ten-carbon chain substrate lysophospholipid 1-decanoyl-2-hydroxy-*sn*-glycero-3-phosphocholine **1e** (0.5 mM) with decanoyl donor **2e**, 2-(decanoylthio)-*N*,*N*,*N*-trimethylethan-1-aminium chloride (0.75 mM) at 37 °C for 5 hours, no desired 1,2-didecanoyl-*sn*-glycero-3-phosphocholine (DDPC) was observed (Fig. S9). However, when the concentration of the reactants was raised about 10 times, such that micelles were formed (Fig. S2C and S2F), we were able to obtain DDPC in 83% yield (Fig. S9), providing further evidence that mixed micelle formation is necessary for transacylation to occur.

Enabling the synthesis of phospholipids through esterification reactions in water should result in the spontaneous *de novo* formation of cell-like lipid membranes. This is due to the expected micelle to lamellar transition as single-chain lysolipids are converted to phospholipids. We used time-lapse fluorescence microscopy to observe the *de novo* formation of vesicles during the optimized reaction (Fig. 4A). Neither lysophospholipid **1a** nor the oleoyl thioester **2b** alone formed vesicles in water, and no visible structures (Fig. 4A) were observed after initial mixing of **1a** (0.5 mM) and **2b** (0.75 mM) in the presence of Na_2_CO_3_/NaHCO_3_ buffer of pH = 8.8 at 37 °C. However, after 10 minutes tubular structures appeared, and after 30 minutes spherical vesicles were observed. Likewise, large vesicles were formed when **1a** (0.5 mM) and **2b** (0.75 mM) were mixed in water collected from the LCHF (Figure 4B). Membrane-bound vesicles were also detected upon mixing the reactants in Mono Lake water, and transmission electron microscopy (TEM) was able to confirm the generation of vesicles (Fig. 4C).

**Fig. 4.**
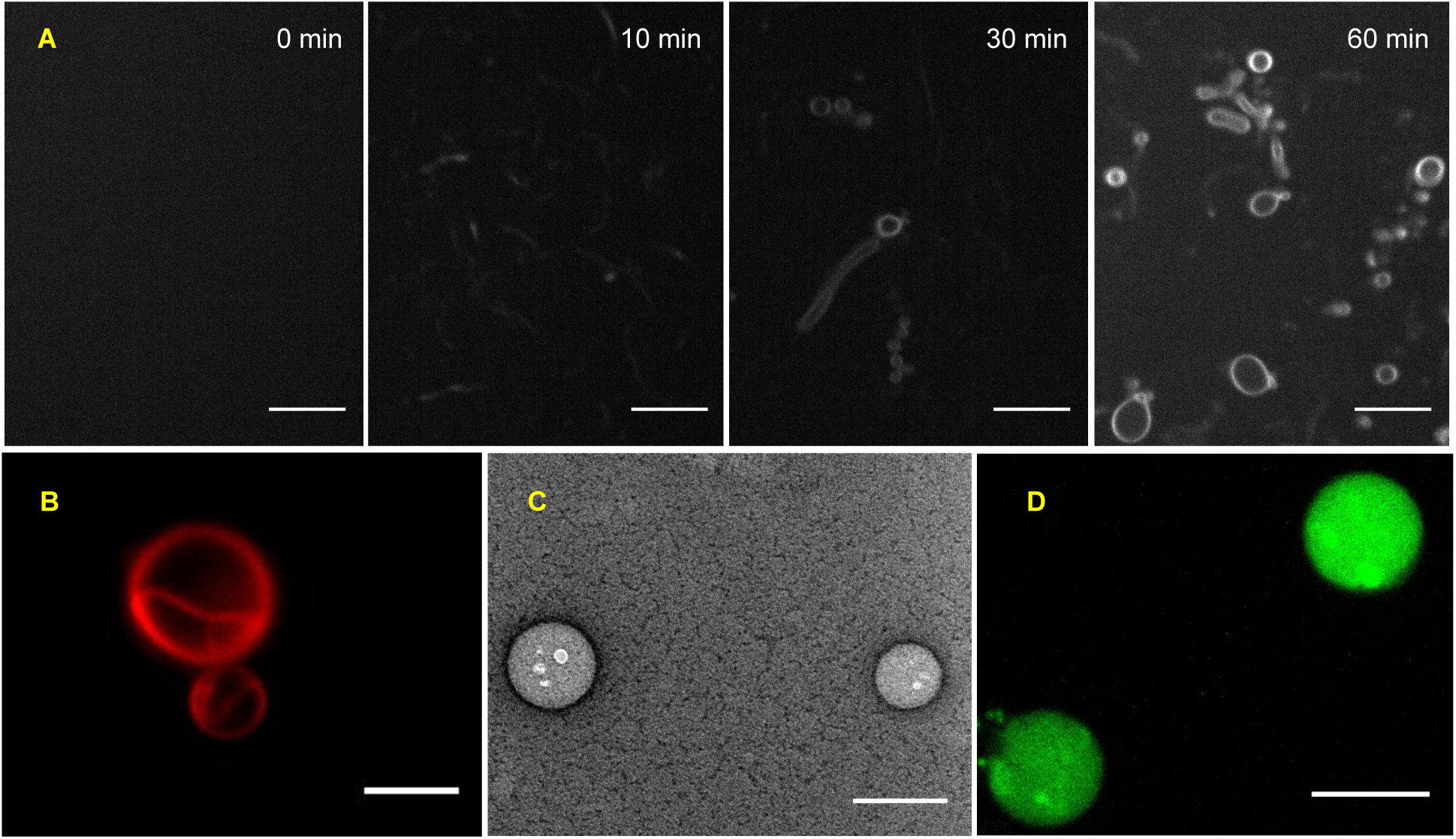
Nonenzymatic formation of phospholipid membranes. The reaction was carried out by mixing 1-oleoyl-2-hydroxy-*sn*-glycero-3-phosphocholine **1a** (0.5 mM), oleoylation reagent **2b** (0.75 mM) in different solvents at 37 °C. (**A**) Na_2_CO_3_/NaHCO_3_ buffer (pH = 8.8) was used as a solvent. Samples were taken at different time points and were stained by using 0.1 mol % Nile red dye. Fluorescence micrographs are presented. Scale bar: 10 μm. (**B**) Lost City vent fluid was used as a solvent. The sample was taken after 48 hours and was stained by using 0.1 mol % Nile red dye. Fluorescence micrograph is presented. Scale bar: 5 μm. (**C**) Mono Lake water was used as a solvent. The sample was taken after 5 hours. Negative staining transmission electron micrograph is presented. Scale bar: 200 nm. (**D**) Fluorescence micrograph of vesicles formed in Mono Lake water (pH=10) containing the pH indicator dye 8-hydroxypyrene-1,3,6-trisulfonic acid (HPTS), two hours after the external media was exchanged to citrate buffer (pH = 4.6) using spin-filtration. Scale bar: 10 μm.

All living organisms harness proton gradients across phospholipid membranes to generate energy. Vesicles formed from simpler amphiphiles, such as fatty acids, are unable to maintain proton gradients, suggesting that phospholipid membranes appeared early in the origin of life.(*27*) *De novo* phospholipid vesicle formation in alkaline solution results in an alkaline interior of formed vesicles. If the exterior solution is then exchanged for a more acidic media, a proton gradient might spontaneously form. To test whether this was possible, a water-soluble pH indicator dye, 8-hydroxypyrene-1,3,6-trisulfonic acid (HPTS), was added prior to the reaction of **1a** with **2b** in the alkaline Mono Lake water (pH = 10). HPTS is highly fluorescent at alkaline pH, but >99% fluorescence is quenched at pH = 4.6 (Fig. S13). After 5 hours of reaction, microscopy indicated that HPTS was successfully encapsulated in the in situ formed membrane vesicles (Fig. S14A). To test if a proton gradient could be maintained, the vesicle media was exchanged to citrate buffer (pH = 4.6). Microscopy showed the fluorescence intensity was initially maintained, indicating the spontaneous formation of a proton gradient. Fluorescence slowly diminished by 80% over 2 hours (Fig. 4D). This finding demonstrates that phospholipid membranes generated in alkaline water sources are capable of forming and maintaining a proton gradient over hours. It is tempting to speculate that similar phenomena may have occurred in the early origin of membranes, perhaps as alkaline hydrothermal vent water was diluted in the more acidic water of the Hadean ocean. (*28*, *29*) Such a scenario may have driven selection pressures for developing primitive catalysts for maintaining such proton gradients, which could have eventually been utilized to form chemical energy.

## Supporting information

Experimental materials and methods

## Acknowledgments

N. K. D. acknowledges financial support for this work provided by the National Science Foundation (CHE-1254611) and the Human Frontier Science Program (LIY000237/2016). K. N. H. acknowledges National Science Foundation (CHE-1764328), the National Institutes of Health, National Institute of General Medical Sciences (R01 GM109078), for financial support of this research. S. Q. L. acknowledges financial support for this work provided by National Science Foundation (OCE-1536702). Computation time was provided by the UCLA Institute for Digital Research and Education (IDRE).

## Supplementary Materials

Materials and methods

Supplementary Text

Fig. S1–S17

Table S1

Copies of HPLC and NMR Spectra Reference (*20*)

## References and Notes

1. S. Jackowski, J. E. Cronan Jr., C. O. Rock, Biochemistry of Lipids, Lipoproteins and Membranes, Elsevier. 20, 80–81 (1991).

2. W. E. M. Lands, J. Biol. Chem. 231, 883–888 (1958).

3. P. K. Schmidli, P. Schurtenberger, P. L. Luisi, J. Am. Chem. Soc. 113, 8127–8130 (1991).

4. D. W. Deamer, D. E. Boatman, J. Cell Biol. 84, 461–467 (1980).

5. S. L. Morris-Natschke, F. Gumus, C. J. Marasco Jr., K. L. Meyer, M. Marx, C. Layne, M. D. Piantadosi, E. J. Modest, J. Med. Chem. 36, 2018–2025 (1993).

6. T. Harayama, M. Eto, H. Shindou, Y. Kita, E. Otsubo, D. Hishikawa, S. Ishii, K. Sakimura, M. Mishina, T. Shimizu, Cell Metabolism 20, 295–305 (2014).

7. W. R. Hargreaves, S. J. Mulvihill, D. W. Deamer, Nature 266, 78–80 (1977).

8. M. Rao, J. Eichberg, J. Oró, J. Mol. Evol. 18, 196–202 (1982).

9. C. Fernandez-Garcia, M. W. Powner, Synlett 28, 78–83 (2017).

10. C. Bonfio, C. Caumes, C. D. Duffy, B. H. Patel, C. Percivalle, M. Tsanakopoulou, J. D. Sutherland, J. Am. Chem. Soc. 141, 3934–3939 (2019).

11. J. W. Szostak, D. P. Bartel, P. L. Luisi, Nature 409, 387–390 (2001).

12. I. Budin, J. W. Szostak, Proc. Natl. Acad. Sci. U. S. A. 108, 5249–5254 (2011).

13. C. de Ouve, American Scientist 83, 428–437 (1995).

14. C. S. Ejsinga, J. L. Sampaioa, V. Surendranatha, E. Duchoslavb, K. Ekroosc, R. W. Klemma, K. Simonsa, A. Shevchenkoa, Proc. Natl. Acad. Sci. U. S. A. 106, 2136–2141 (2009).

15. R. J. Brea, C. M. Cole, N. K. Devaraj, Angew. Chem. Int. Ed. 53, 14102–14105 (2014).

16. M. L. Bender, Chem. Rev. 60, 53–113 (1960).

17. R. A. McClelland, L. J. Santry, Acc. Chem. Res. 16, 394–399 (1983).

18. M. Raynal, P. Ballester, A. Vidal-Ferran, P. W. N. M. van Leeuwen, Chem. Soc. Rev. 43, 1660–1733 (2014).

19. G. Steinman, M. N. Cole, Proc. Natl. Acad. Sci. U. S. A. 58, 735–742 (1967).

20. All calculations were carried out using Gaussian 09, Revision A.02 (full citation please see Supporting Information).

21. A.V. Marenich, C. J. Cramer, D. G. Truhlar, J. Chem. Phys. B 113, 6378–6396 (2009).

22. D. E. Delory, E. J. King, Biochem. J. 39, 245 (1945).

23. S. Kempe, E. T. Degens, Chem. Geol. 53, 95–108 (1985).

24. W. Martin, M. J. Russell, Phil. Trans. R. Soc. B Biol. Sci. 362, 1887–1925 (2007).

25. W. R. Perkins, R. B. Dause, R. A. Parente, S. R. Minchey, K. C. Neuman, S. M. Gruner, T. F. Taraschi, A. S. Janoff, Science 273, 330–332 (1996).

26. K. Athenstaedt, G. Daum, Eur. J. Biochem. 266, 1–16 (1999).

27. I. A. Chen, J. W. Szostak, Proc. Natl. Acad. Sci. U. S. A. 101, 7965–7970 (2004).

28. J. W. Morese, F. T. Mackenzie, Aquat. Geochem. 4, 301–319 (1998).

29. G. Macleod, C. McKeown, A. J. Hall, M. J. Russell, Orig. Life Evol. Biospheres 24, 19–41 (1994).

