## Supplementary material for "Enzyme-free synthesis of natural phospholipids in water": Experimental materials and methods

**This PDF file includes:**

Materials and methods
Supplementary Text
Fig. S1−S17
Table S1
Copies of HPLC and NMR Spectra
Reference (*20*)

**Table of Contents**


| | | |
|---|---|---|
| | General Information | S3 |
| 1. | Catalyst Screening for The Transacylation Reaction | S4 |
| 2. | Synthesis of Acyl Thioester **2b**–**2e** | S5 |
| 3. | Critical Micelle Concentration of Reactants | S9 |
| 4. | General Procedure for Transacylation Reaction | S10 |
| 5. | Substrate Exploration | S12 |
| 6. | In Situ Formation of Liposomes | S16 |
| 7. | In Situ Encapsulation of HPTS and Proton Gradient Exploration | S18 |
| 8. | HPLC/ELSD Spectra | S19 |
| 9. | $^1$H and $^{13}$C NMR Spectra | S21 |
| 10. | Theoretical Study | S33 |
| 11. | Full Reference List | S46 |

**General information**

Chemicals were purchased as reagent grade and used without further purification except as indicated below. Lysophospholipids were purchased from Avanti Polar Lipids, Inc. Solvents ($CH_2Cl_2$, $CHCl_3$, MeOH, DMF, THF, $H_2O$) were purchased from commercial suppliers. Solvents were removed under reduced pressure using a rotary evaporator and drying under high vacuum if appropriate ($10^{-3}$ mbar). Thin-layer chromatography (TLC) was performed using silica gel pre-coated plastic sheets (Polygram SIL G/$UV_{254}$, 0.2 mm, with fluorescent indicator; Macherey-Nagel) plastic sheets, which were visualized with a UV lamp (254), or Potassium Permanganate stain (1.5g of $KMnO_4$, 10g $K_2CO_3$, and 1.25mL 10% NaOH in 200 mL $H_2O$). Column chromatography (CC) was carried out using Merck silica gel (60 Å, 230−400 mesh, particle size 0.040−0.063 mm) using technical grade solvents. Elution was accelerated using compressed air. All reported yields, unless otherwise specified, refer to spectroscopically and chromatographically pure compounds. Nomenclature follows the suggestions proposed by the computer program ChemBioDraw of CBD/CambridgeSoft. $^{1}H$, $^{13}C$ nuclear magnetic resonance (NMR) spectra were recorded on a Bruker ECA-500 or VX-500 spectrometer in a suitable deuterated solvent. The solvent employed and respective measuring frequency are indicated for each experiment. Chemical shifts are reported with tetramethylsilane (TMS) serving as a universal reference for all nuclides. The resonance multiplicity is described as s (singlet), d (doublet), t (triplet), q (quadruplet), m (multiplet), and br (broad). All spectra were recorded at 298 K unless otherwise noted, processed with MestReNova 12.0.0 suite, and coupling constants are reported as observed. The residual deuterated solvent signal relative to tetramethylsilane was used as the internal reference in $^{1}H$ NMR spectra (e.g. $CDCl_3$ = 7.26 ppm), and is reported as follows: chemical shift in ppm (multiplicity, coupling constant $J$ in Hz, number of protons). $^{13}C$, $^{19}F$, $^{31}P$ NMR spectra were referenced according to Ξ-values (IUPAC recommendations 2008) relative to the internal references set in $^{1}H$ NMR spectra (e.g. $^{13}C$: $Me_4Si$ 0.00 ppm). An Agilent 6230 time-of-flight mass spectrometer (TOF-MS) with Jet Stream electrospray ionization source (ESI) was used for high resolution mass spectrometry (HR-MS) analysis. The Jet Stream ESI source was operated under positive ion mode with the following parameters: VCap: 3500 V; fragmentor voltage: 160 V; nozzle voltage: 500 V; drying gas temperature: 325 °C, sheath gas temperature: 325 °C, drying gas flow rate: 7.0 L/min; sheath gas flow rate: 10 L/min; nebulizer pressure: 40 psi. High performance liquid chromatography (HPLC) was performed on ZORBAX SB-C18 liquid chromatograph. Liquid chromatography-mass spectrometry (LC-MS) was performed on Eclipse Plus C8 analytical chromatograph. All solvents used were HPLC-grade solvents purchased from Fisher Scientific. Transmission electron microscopy (TEM) images were recorded on a FEI Tecnai™ Spirit G2 BioTWIN 120 kV microscope equipped with a 4k (16 megapixel) Eagle camera. Spinning-disk confocal microscopy images were acquired on a Yokagawa spinning-disk system (Yokagawa, Japan) built around an Axio Observer Z1motorized inverted microscope (Carl Zeiss S-3 Microscopy GmbH, Germany) with a 63x, 1.40 NA oil immersion objective to an Evolve 512×512 EMCCD camera (Photometrics, Canada) using ZEN imaging software (Carl Zeiss Microscopy GmbH, Germany). A condenser/objective with a phase stop of Ph2 was used to obtain the phase-contrast images. Fluorescence measurements were performed on a Tecan infinite F200 plate reader instrument.

### 1. Catalyst Screening for The Transacylation Reaction

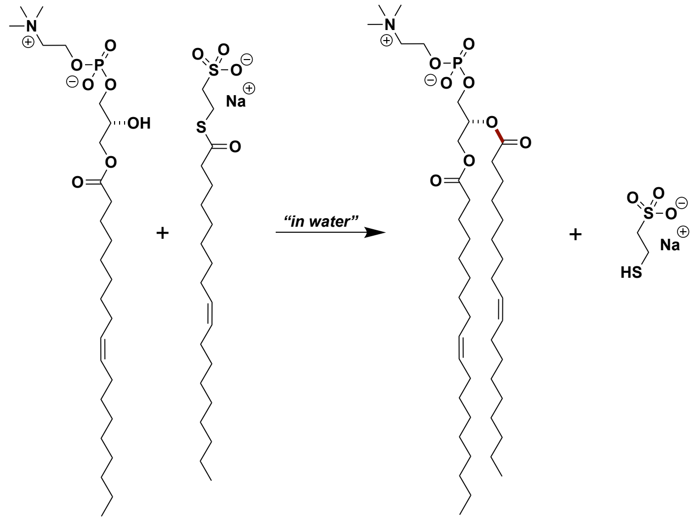


1-Oleoyl-2-hydroxy-*sn*-glycero-3-phosphocholine **1a** and sodium 2-(oleoylthio)ethane-1-sulfonate **2a** were added to the catalyst in water (100 μL). The reaction mixture was stirred at 37 °C for 24 hours.

**Table S1. Catalyst screening for the transacylation reaction.**

| Entry | Catalyst/μmol [a] | **1a**/μmol | **2a**/μmol | Time/hour | Yield[b] |
|---|---|---|---|---|---|
| 1 | DBSA/0.6 | 4 | 6 | 24 | N.D. |
| 2 | $ZnCl_2$/0.6 | 4 | 6 | 24 | N.D. |
| 3 | DMAP/0.6 | 4 | 6 | 24 | N.D. |
| 4 | Imidazole/0.6 | 4 | 6 | 24 | N.D. |
| 5 | DBSA/60 | 4 | 6 | 24 | N.D. |
| 6 | $ZnCl_2$/60 | 4 | 6 | 24 | <5% |
| 7 | DMAP/60 | 4 | 6 | 24 | <10% |
| 8 | Imidazole/60 | 4 | 6 | 24 | 12% |

[a]DMAP, 4-dimethylaminopyridine; DBSA, *p*-dodecylbenzene-sulfonic acid. [b]Yields of product **3a** were determined by LC-MS of reaction mixtures on the basis of the calibration curve of pure **3a**.

### 2. Synthesis of Acyl Thioester 2b–2e

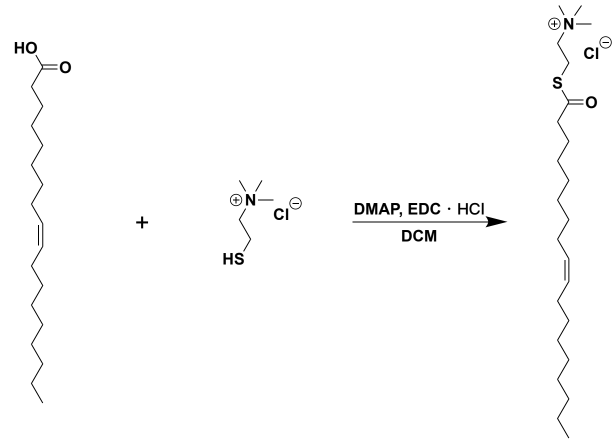


oleic acid cholinethiol chloride 2b

**2-(oleoylthio)-*N*,*N*,*N*-trimethylethan-1-aminium chloride (2b)**

4-Dimethylaminopyridine (15 mg, 0.12 mmol) and *N*-(3-dimethylaminopropyl)-*N′*-ethylcarbodiimide hydrochloride (EDC•HCl) (248 mg, 1.3 mmol) was added to a solution of oleic acid (367 mg, 1.3 mmol) in $CH_2Cl_2$ (10 mL) at 0 °C under argon. The reaction mixture was stirred at 0 °C for 30 minutes. The solution was added dropwise to thiocholine chloride (188 mg, 1.2 mmol) at –78 °C under argon. The reaction was warmed to room temperature and stirred for 4 hours. The reaction solvent was removed under reduced pressure. The residue was purified by column chromatography on silica gel using 10%–50% MeOH/$CH_2Cl_2$ as the eluent yielding the title compound as a colorless solid (490 mg, 97%).

**$^1$H NMR** (500 MHz, $CD_3OD$): δ 5.35 (t, *J* = 5.0 Hz, 2 H), 3.47–3.43 (m, 2 H), 3.29–3.25 (m, 2 H), 3.22 (s, 9 H), 2.65 (t, *J* = 5.0 Hz, 2 H), 2.07–2.01 (m, 4 H), 1.67 (t, *J* = 5.0 Hz, 2 H), 1.35–1.30 (m, 20 H), 0.91 (t, *J* = 5 Hz, 3 H).

**$^{13}$C NMR** (126 MHz, $CD_3OD$): δ 199.29, 130.82, 130.69, 65.99, 53.63, 44.75, 44.64, 33.03, 30.82, 30.76, 30.61, 30.44, 30.34, 30.26, 30.13, 29.94, 28.18, 28.13, 28.08, 26.57, 26.44, 26.30, 23.72, 22.65, 14.62.

**HRMS** *m/z* (ESI): calcd. for $C_{23}H_{46}NOS$ [M-Cl]$^+$: 384.3295; found: 384.3293.

**Prebiotically plausible route to thioester 2b**

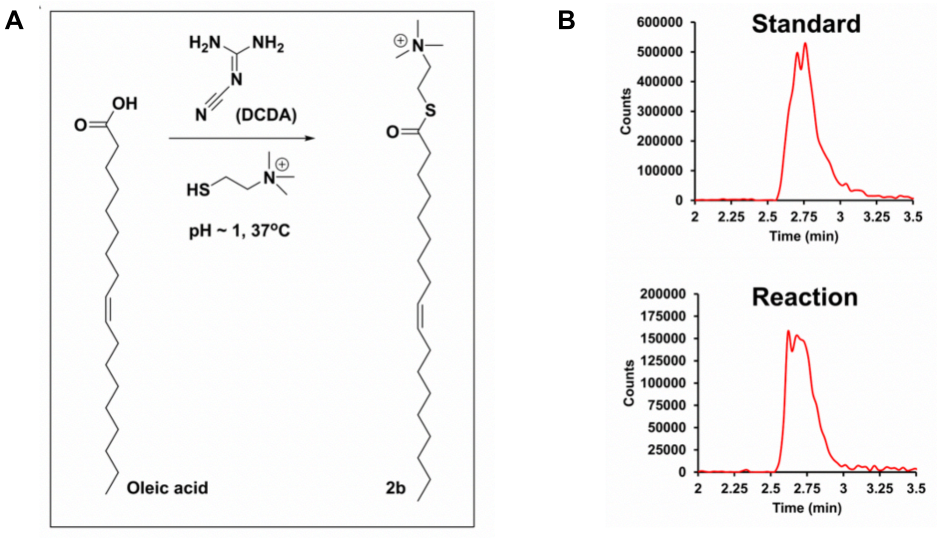


**Figure S1. Prebiotically plausible route to thioester 2b**. **A**. Reaction scheme for dicyandiamide (DCDA) mediated condensation of oleic acid and thiocholine chloride to form thioester **2b**. **B**. Comparison of Extracted Ion Chromatogram (EIC) corresponding to m/z 384 (*positive ion mode*) for an analytical standard sample of **2b** (*top*) and that of **2b** generated in the reaction (*bottom*).

Method: A turbid dispersion of oleic acid (15.5 mg) and dicyandiamide (32.7 mg) was prepared in 0.1 mL aqueous HCl (pH ~1) by stirring at room temperature. Into this, choline thiol chloride (18.8 mg, dissolved in 0.1 mL $H_2O$) is added and the dispersion was tumbled at 37 °C overnight. A small volume of the dispersion is analyzed on HPLC-MS and the compound **2b** could be detected.

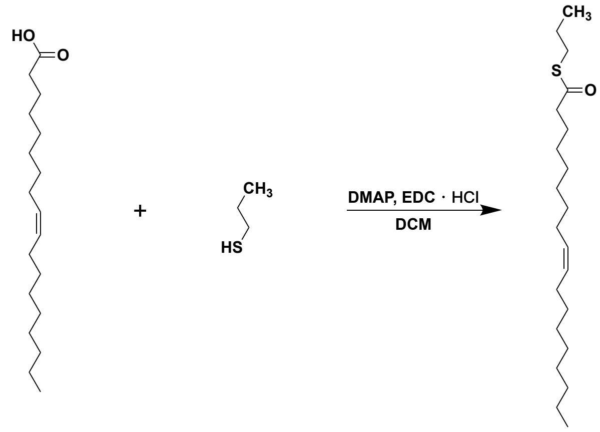


oleic acid propane-1-thiol 2c

**$S$-propyl ($Z$)-octadec-9-enethioate (2c)**

4-Dimethylaminopyridine (15 mg, 0.12 mmol) and *N*-(3-dimethylaminopropyl)-*N'*-ethylcarbodiimide hydrochloride (EDC•HCl) (248 mg, 1.3 mmol) was added to a solution of oleic acid (367 mg, 1.3 mmol) in $CH_2Cl_2$ (10 mL) at 0 °C under argon. The reaction mixture was stirred at 0 °C for 30 minutes. Propane-1-thiol (91 mg, 1.2 mmol) was added dropwise to the reaction mixture at 0 °C under argon. The reaction was warmed to room temperature and stirred for 4 hours. The reaction solvent was removed under reduced pressure. The residue was purified by column chromatography on silica gel using 30%–50% Hexane/$CH_2Cl_2$ as the eluent yielding the title compound as a colorless oil (395 mg, 97%).

**$^{1}$H NMR** (500 MHz, $CDCl_3$): δ 5.43 (dt, $J$ = 10.0 Hz, $J$ = 5.0 Hz, 2 H), 2.85 (t, $J$ = 5.0 Hz, 2 H), 2.53 (t, $J$ = 5.0 Hz, 2 H), 2.00 (q, $J$ = 5.0 Hz, 4 H), 1.65 (t, $J$ = 5.0 Hz, 2 H), 1.62–1.54 (m, 2 H), 1.26–1.33 (m, 20 H), 0.96 (t, $J$ = 5.0 Hz, 3 H), 0.88 (t, $J$ = 5.0 Hz, 3 H).

**$^{13}$C NMR** (126 MHz, $CDCl_3$): δ 199.96, 130.16, 129.88, 44.31, 32.06, 30.83, 29.92, 29.81, 29.67, 29.47, 29.32, 29.30, 29.20, 29.07, 27.37, 27.30, 25.86, 23.15, 22.83, 14.27, 13.48.

**HRMS** *m/z* (ESI): calcd. for $C_{21}H_{41}OS$ $[M+H]^+$: 341.2873; found: 341.2877.

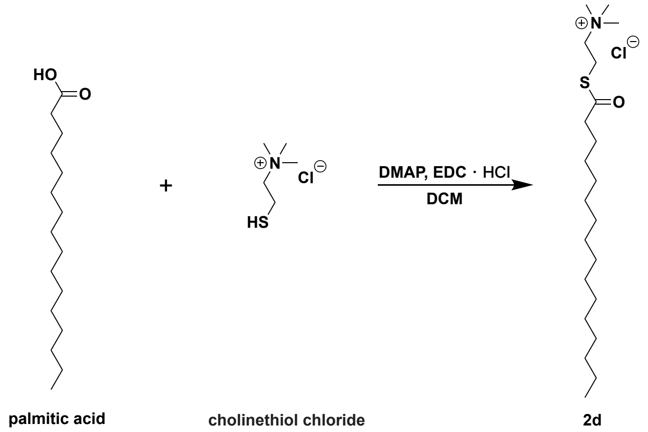


palmitic acid cholinethiol chloride 2d

**2-(palmitoylthio)-*N*,*N*,*N*-trimethylethan-1-aminium chloride (2d)**

4-Dimethylaminopyridine (15 mg, 0.12 mmol) and *N*-(3-dimethylaminopropyl)-*N'*-ethylcarbodiimide hydrochloride (EDC•HCl) (248 mg, 1.3 mmol) was added to a solution of palmitic acid (333 mg, 1.3 mmol) in $CH_2Cl_2$ (10 mL) at 0 °C under argon. The reaction mixture was stirred at 0 °C for 30 minutes. Then thiocholine chloride (188 mg, 1.2 mmol) was added to the solution at 0 °C under argon. The reaction was warmed to room temperature and stirred for 4 hours. Then the reaction solvent was removed under reduced pressure. The residue was purified by column chromatography on silica gel using 10%–50% MeOH/$CH_2Cl_2$ as the eluent yielding the title compound as a colorless solid (150 mg, 32%).

**$^1$H NMR** (500 MHz, $CD_3OD$): δ 3.47–3.44 (m, 2 H), 3.29–3.25 (m, 2 H), 3.22 (s, 9 H), 2.65 (t, $J$ = 5.0 Hz, 2 H), 1.67 (t, $J$ = 5.0 Hz, 2 H), 1.36–1.29 (m, 24 H), 0.90 (t, $J$ = 5 Hz, 3 H).

**$^{13}$C NMR** (126 MHz, $CD_3OD$): δ 199.55, 66.10, 53.67, 53.64, 53.61, 44.64, 33.04, 30.77, 30.74, 30.69, 30.52, 30.45, 30.35, 29.94, 26.47, 23.71, 22.61, 14.50.

**HRMS** *m/z* (ESI): calcd. for $C_{21}H_{44}NOS$ $[M-Cl]^+$: 358.3138; found: 358.3139.

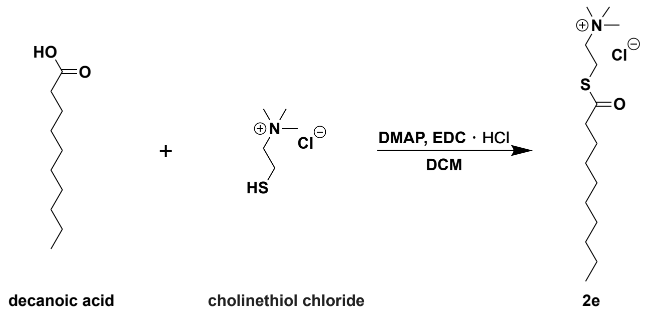


**2-(decanoylthio)-*N*,*N*,*N*-trimethylethan-1-aminium chloride (2e)**

4-Dimethylaminopyridine (15 mg, 0.12 mmol) and *N*-(3-dimethylaminopropyl)-*N*′-ethylcarbodiimide hydrochloride (EDC•HCl) (248 mg, 1.3 mmol) was added to a solution of decanoic acid (224 mg, 1.3 mmol) in $CH_2Cl_2$ (10 mL) at 0 °C under argon. The reaction mixture was stirred at 0 °C for 30 minutes. Then thiocholine chloride (188 mg, 1.2 mmol) was added to the solution at 0 °C under argon. The reaction was warmed to room temperature and stirred for 4 hours. Then the reaction solvent was removed under reduced pressure. The residue was purified by column chromatography on silica gel using 10%–20% MeOH/$CH_2Cl_2$ as the eluent yielding the title compound as a colorless solid (120 mg, 33%).

**$^1$H NMR** (500 MHz, $CD_3OD$): δ 3.40–3.37 (m, 2 H), 3.25–3.21 (m, 2 H), 3.16 (s, 9 H), 2.58 (t, $J$ = 5.0 Hz, 2 H), 1.60 (t, $J$ = 5.0 Hz, 2 H), 1.29–1.23 (m, 12 H), 0.84 (t, $J$ = 5 Hz, 3 H).

**$^{13}$C NMR** (126 MHz, $CD_3OD$): δ 199.60, 66.10, 53.54, 53.51, 53.48, 44.64, 33.05, 30.54, 30.41, 29.95, 26.53, 23.75, 22.53, 14.48.

**HRMS** *m/z* (ESI): calcd. for $C_{15}H_{32}NOS$ $[M-Cl]^+$: 274.2199; found: 274.2197.

### 3. Critical Micelle Concentration of Reactants

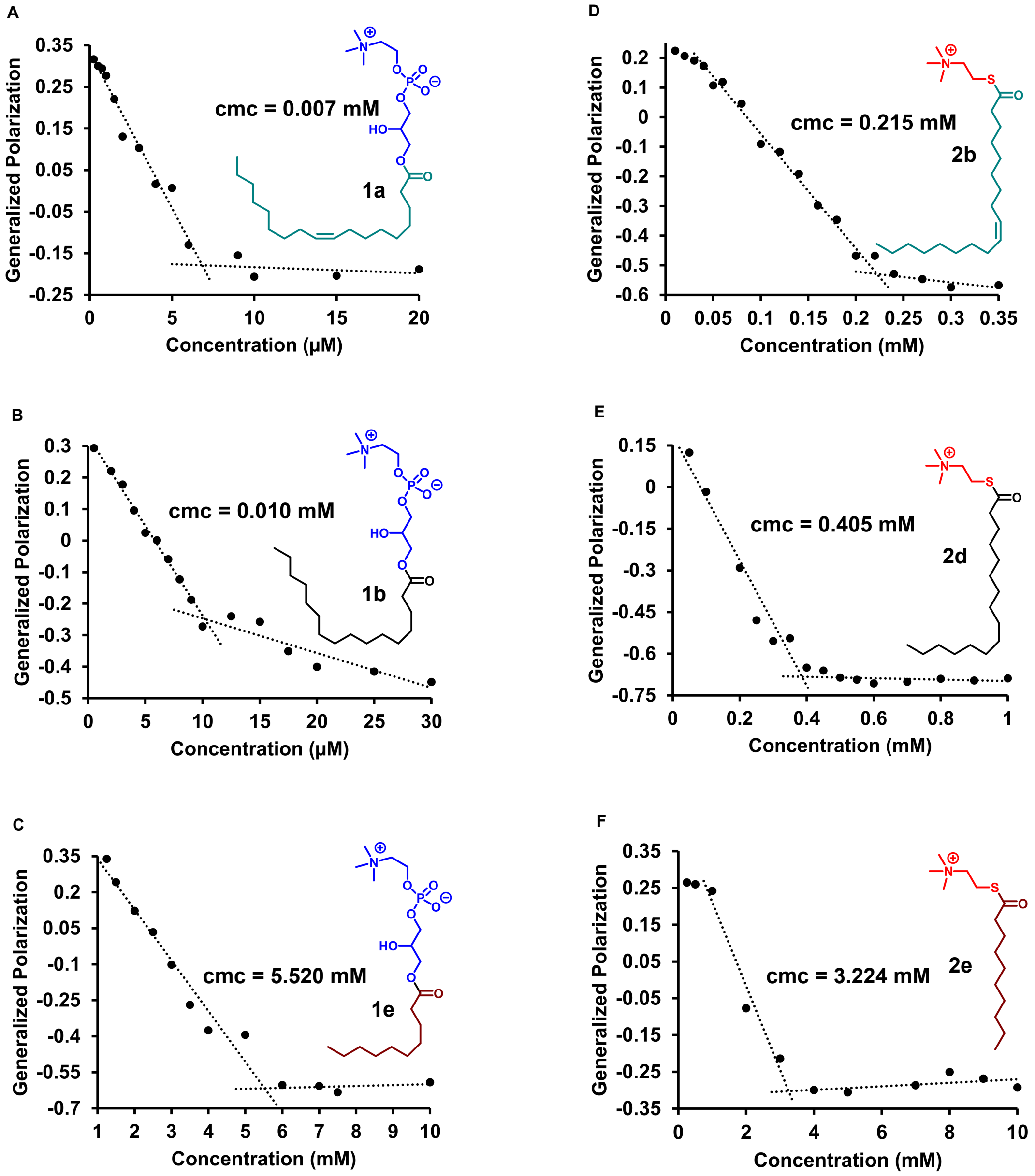


**Figure S2. Critical micelle concentration (cmc) of reactants.** The cmc values of reactants were estimated using a method based on the fluorescence properties of the dye Laurdan in water at 37 °C. Representative plots are shown.

The cmc's of the various amphiphiles (thioesters and lysolipids) described in this study were estimated using a method based on the solvatochromic fluorescent dye Laurdan. Laurdan shows a sharp change in the value of generalized polarization at or near the cmc [or critical aggregation concentration (cac)] of an amphiphile. Initially, a concentrated solution of the amphiphile was prepared in Milli-Q $H_2O$ by hydration of a thin film. Afterwards, various dilutions were prepared by adding requisite volumes of Milli-Q $H_2O$. The volumes of various dilutions prepared ranged between 0.25-1.0 mL for the amphiphiles in this study. The samples were allowed to equilibrate at 37 ºC for 0.5 h, following which 0.4 µL of Laurdan (2.5 mM in EtOH) were added to each and mixed by gentle tapping. Then, 20 µL of each dilution was transferred to a 384 well plate and analyzed on a Tecan Infinite Plate Reader maintained at 37 ºC. The samples were excited at 364 nm and emission spectra acquired over 430-500 nm. Generalized polarization (GP) was calculated as follows: GP = $(I_{440}-I_{490})/(I_{440}+I_{490})$.

#### 4. General Procedure for Transacylation Reaction

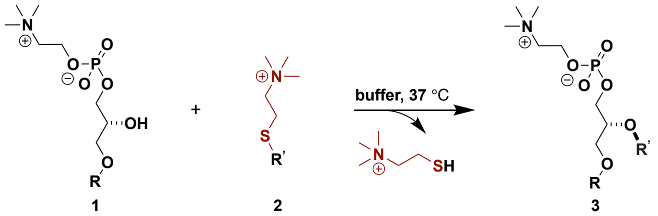


Unless specified otherwise, to a mixture of lysophospholipid **1** (4 µmol) and acylation reagent **2** (6 µmol), freshly prepared $Na_2CO_3$/$NaHCO_3$ buffer of pH = 10.6 (8 mL) was added at 37 °C. The reaction mixture was tumbled at 37 °C for 5 hours. Purification was performed by column chromatography or preparative thin layer chromatography on silica gel.

**1,2-Dioleoyl-*sn*-glycero-3-phosphocholine (3a)**

A white powder was obtained. An isolated yield of 88% was obtained using 60/40/2–80/20/2 MeOH/$CHCl_3$/$NH_3$•$H_2O$ as the eluents.

**$^1$H NMR** (500 MHz, $CDCl_3$): δ 5.36–5.29 (m, 4 H), 5.18 (dq, *J* = 10.0 Hz, *J* = 5.0 Hz, 1 H), 4.38 (dd, *J* = 10.0 Hz, *J* = 5.0 Hz, 1 H), 4.31–4.28 (m, 2 H), 4.11 (q, *J* = 5.0 Hz, 1 H), 3.97–3.87 (m, 2 H), 3.79 (t, *J* = 5.0 Hz, 2 H), 3.68 (d, *J* = 5.0 Hz, 2 H), 3.36 (s, 9 H), 2.27 (q, *J* = 5.0 Hz, 4 H), 2.00 (q, *J* = 5.0 Hz, 8 H), 1.59–1.54 (m, 4 H), 1.33–1.25 (m, 38 H), 0.86 (t, *J* = 5 Hz, 6 H).

**$^{13}$C NMR** (126 MHz, $CDCl_3$): δ 173.66, 173.30, 130.12, 129.79, 70.63, 66.46, 63.45, 63.41, 63.12, 59.42, 59.38, 54.51, 34.43, 34.23, 32.03, 29.89, 29.66, 29.45, 29.40, 29.38, 29.31, 29.27, 29.24, 27.34, 27.32, 25.08, 25.00, 22.82, 14.27.

**HRMS** *m/z* (ESI): calcd. for $C_{44}H_{84}NO_8PNa$ $[M+Na]^+$: 808.5827; found: 808.5826.

**1-Palmitoyl-2-oleoyl-*sn*-glycero-3-phosphocholine (3b)**

A white powder was obtained. An isolated yield of 85% was obtained using 60/40/2–80/20/2 MeOH/$CHCl_3$/$NH_3$•$H_2O$ as the eluents.

**$^1$H NMR** (500 MHz, $CDCl_3$): δ 5.37–5.30 (m, 2 H), 5.20 (dq, *J* = 10.0 Hz, *J* = 5.0 Hz, 1 H), 4.39 (dd, *J* = 10.0 Hz, *J* = 5.0 Hz, 1 H), 4.34–4.30 (m, 2 H), 4.11 (q, *J* = 5.0 Hz, 1 H), 3.98–3.89 (m, 2 H), 3.83 (t, *J* = 5.0 Hz, 2 H), 3.38 (s, 9 H), 3.12–3.02 (m, 2 H), 2.27 (dq, *J* = 10.0 Hz, *J* = 5.0 Hz, 4 H), 2.00 (q, *J* = 5.0 Hz, 4 H), 1.60–1.53 (m, 2 H), 1.29–1.24 (m, 44 H), 0.87 (t, *J* = 5.0 Hz, 6 H).

**$^{13}$C NMR** (126 MHz, $CDCl_3$): δ 173.74, 173.36, 130.17, 130.13, 129.81, 70.73, 70.66, 70.59, 66.54, 63.51, 63.29, 63.21, 63.09, 59.41, 59.37, 54.84, 54.70, 54.54, 34.62, 34.45, 34.43, 34.26, 34.10, 32.06, 29.90, 29.87, 29.82, 29.70, 29.68, 29.52, 29.49, 29.47, 29.43, 29.33, 29.26, 27.36, 27.33, 25.10, 25.03, 22.96, 22.84, 22.69, 14.45, 14.34, 14.29, 14.23, 14.12.

**HRMS** *m/z* (ESI): calcd. for $C_{42}H_{82}NO_8PNa$ $[M+Na]^+$: 782.5670; found: 782.5671.

**1-Stearoyl-2-oleoyl-*sn*-glycero-3-phosphocholine (3c)**

A white powder was obtained. An isolated yield of 78% was obtained using 60/40/2–80/20/2 MeOH/$CHCl_3$/$NH_3$•$H_2O$ as the eluents.

**$^1$H NMR** (500 MHz, $CDCl_3$): δ 5.36–5.29 (m, 2 H), 5.20 (dq, *J* = 10.0 Hz, *J* = 5.0 Hz, 1 H), 4.39 (dd, *J* = 10.0 Hz, *J* = 5.0 Hz, 1 H), 4.32–4.30 (m, 2 H), 4.11 (q, *J* = 5.0 Hz, 1 H), 3.99–3.90 (m, 2 H), 3.81 (t, *J* = 5.0 Hz, 2 H), 3.37 (s, 9 H), 3.06–3.04 (m, 2 H), 2.28 (q, *J* = 10.0 Hz, 4 H), 2.00 (q, *J* = 5.0 Hz, 4 H), 1.60–1.53 (m, 4 H), 1.29–1.24 (m, 46 H), 0.87 (t, *J* = 5.0 Hz, 6 H).

$^{13}$**C NMR** (126 MHz, $CDCl_3$): δ 173.73, 173.34, 130.16, 130.15, 130.13, 129.83, 129.81, 70.66, 66.56, 63.48, 63.43, 63.13, 59.41, 59.37, 54.62, 34.45, 34.27, 32.07, 32.04, 29.90, 29.89, 29.86, 29.82, 29.81, 29.76, 29.69, 29.68, 29.51, 29.48, 29.47, 29.41, 29.38, 29.32, 29.28, 29.24, 27.36, 27.33, 25.09, 25.02, 22.84, 22.83, 14.28.
**HRMS** *m/z* (ESI): calcd. for $C_{44}H_{86}NO_8PNa$ $[M+Na]^+$: 810.5983; found: 810.5972.

**1-Oleoyl-2-palmitoyl-*sn*-glycero-3-phosphocholine (3d)**
A white powder was obtained. An isolated yield of 87% was obtained using 60/40/2–80/20/2 $MeOH/CHCl_3/NH_3$•$H_2O$ as the eluents.
$^1$**H NMR** (500 MHz, $CDCl_3$): δ 5.37–5.30 (m, 2 H), 5.20 (dq, *J* = 10.0 Hz, *J* = 5.0 Hz, 1 H), 4.39 (dd, *J* = 10.0 Hz, *J* = 5.0 Hz, 1 H), 4.35–4.31 (m, 2 H), 4.12 (q, *J* = 5.0 Hz, 1 H), 4.00–3.91 (m, 2 H), 3.82 (t, *J* = 5.0 Hz, 2 H), 3.38 (s, 9 H), 2.68 (br s, 2 H), 2.28 (q, *J* = 10.0 Hz, 4 H), 2.00 (q, *J* = 5.0 Hz, 4 H), 1.59–1.56 (m, 4 H), 1.33–1.23 (m, 42 H), 0.87 (t, *J* = 5.0 Hz, 6 H).
$^{13}$**C NMR** (126 MHz, $CDCl_3$): δ 173.69, 173.38, 130.15, 130.14, 130.13, 129.84, 129.83, 129.82, 70.69, 70.63, 66.58, 63.49, 63.13, 59.40, 59.36, 54.64, 34.47, 34.25, 32.07, 32.04, 29.96, 29.90, 29.88, 29.86, 29.85, 29.83, 29.81, 29.69, 29.67, 29.52, 29.50, 29.47, 29.41, 29.38, 29.31, 29.28, 29.24, 27.35, 27.32, 25.11, 25.01, 22.84, 22.83, 14.28.
**HRMS** *m/z* (ESI): calcd. for $C_{42}H_{82}NO_8PNa$ $[M+Na]^+$: 782.5670; found: 782.5673.

**1,2-Dipalmitoyl-*sn*-glycero-3-phosphocholine (3e)**
A white powder was obtained. An isolated yield of 87% was obtained using 60/40/2–80/20/2 $MeOH/CHCl_3/NH_3$•$H_2O$ as the eluents.
$^1$**H NMR** (500 MHz, $CDCl_3$): δ 5.18 (dq, *J* = 10.0 Hz, *J* = 5.0 Hz, 1 H), 4.39 (dd, *J* = 10.0 Hz, *J* = 5.0 Hz, 1 H), 4.32–4.28 (m, 2 H), 4.12 (q, *J* = 5.0 Hz, 1 H), 4.96–3.90 (m, 2 H), 3.80 (t, *J* = 5.0 Hz, 2 H), 3.66–3.57 (m, 2 H), 3.37 (s, 9 H), 2.26 (q, *J* = 10.0 Hz, 4 H), 1.59–1.54 (m, 4 H), 1.28–1.24 (m, 46 H), 0.86 (t, *J* = 5.0 Hz, 6 H).
$^{13}$**C NMR** (126 MHz, $CDCl_3$): δ 173.70, 173.34, 70.65, 66.45, 63.46, 63.41, 63.12, 59.42, 59.38, 54.52, 34.46, 34.26, 32.06, 29.86, 29.84, 29.83, 29.81, 29.75, 29.69, 29.68, 29.51, 29.47, 29.31, 29.28, 25.10, 25.02, 22.83, 14.27.
**HRMS** *m/z* (ESI): calcd. for $C_{40}H_{80}NO_8PNa$ $[M+Na]^+$: 756.5514; found: 756.5515.

**1-Stearoyl-2-palmitoyl-*sn*-glycero-3-phosphocholine (3f)**
A white powder was obtained. An isolated yield of 77% was obtained using 60/40/2–80/20/2 $MeOH/CHCl_3/NH_3$•$H_2O$ as the eluents.
$^1$**H NMR** (500 MHz, $CDCl_3$): δ 5.20 (dq, *J* = 10.0 Hz, *J* = 5.0 Hz, 1 H), 4.39 (dd, *J* = 10.0 Hz, *J* = 5.0 Hz, 1 H), 4.34–4.31 (m, 2 H), 4.12 (q, *J* = 5.0 Hz, 1 H), 3.99–3.92 (m, 2 H), 3.81 (t, *J* = 5.0 Hz, 2 H), 3.38 (s, 9 H), 2.83 (br s, 2 H), 2.28 (q, *J* = 10.0 Hz, 4 H), 1.58–1.55 (m, 4 H), 1.29–1.25 (m, 50 H), 0.87 (t, *J* = 5.0 Hz, 6 H).
$^{13}$**C NMR** (126 MHz, $CDCl_3$): δ 173.74, 173.38, 70.70, 70.64, 66.58, 63.49, 63.45, 63.14, 59.40, 59.36, 54.65, 34.48, 34.28, 32.07, 29.87, 29.86, 29.83, 29.82, 29.78, 29.70, 29.52, 29.51, 29.48, 29.32, 29.29, 25.11, 25.03, 22.84, 14.28.
**HRMS** *m/z* (ESI): calcd. for $C_{42}H_{84}NO_8PNa$ $[M+Na]^+$: 784.5827; found: 784.5816.

**1,2-Dioleoyl-*sn*-glycero-3-phosphate sodium salt (3g)**
A white powder was obtained. An isolated yield of 55% was obtained using 20/80–50/50 $MeOH/CHCl_3$ as the eluents.

**$^{1}$H NMR** (500 MHz, CDCl$_3$): δ 5.36–5.29 (m, 4 H), 5.27–5.20 (br s, 1 H), 4.40 (d, $J$ = 10.0 Hz, 1 H), 4.20 (br s, 1 H), 3.88–3.82 (m, 2 H), 3.35–3.36 (m, 3 H), 2.29 (dt, $J$ = 10.0 Hz, $J$ = 5.0 Hz, 4 H), 2.00 (q, $J$ = 5.0 Hz, 8 H), 1.58–1.55 (m, 4 H), 1.28–1.26 (m, 38 H), 0.88 (t, $J$ = 5 Hz, 6 H).

**$^{13}$C NMR** (126 MHz, CDCl$_3$): δ 173.98, 130.07, 129.76, 71.11, 63.29, 62.93, 34.49, 34.30, 32.08, 30.06, 30.04, 29.95, 29.74, 29.58, 29.52, 29.50, 27.44, 27.40, 25.13, 25.05, 22.85, 14.27.

**HRMS** *m/z* (ESI): calcd. for $C_{39}H_{72}O_8P$ [M-Na]$^-$: 699.4970; found: 699.4667.

### 5. Substrate Tolerance

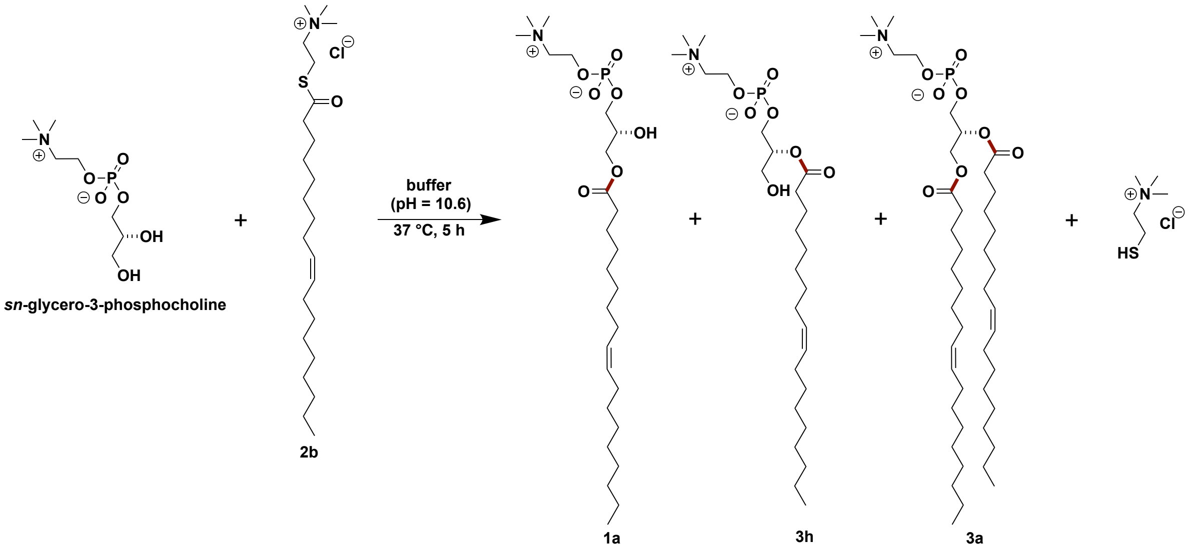


**Figure S3. Oleoylation reaction of *sn*-glycero-3-phosphocholine.** To a mixture of *sn*-glycero-3-phosphocholine (4 μmol) and oleoylation reagent **2b** (6 μmol), $Na_2CO_3/NaHCO_3$ buffer of pH = 10.6 (8 mL) was added. The reaction was tumbled at 37 °C for 5 hours. Acylated products **1a**, **3a**, **3h** were not detected.

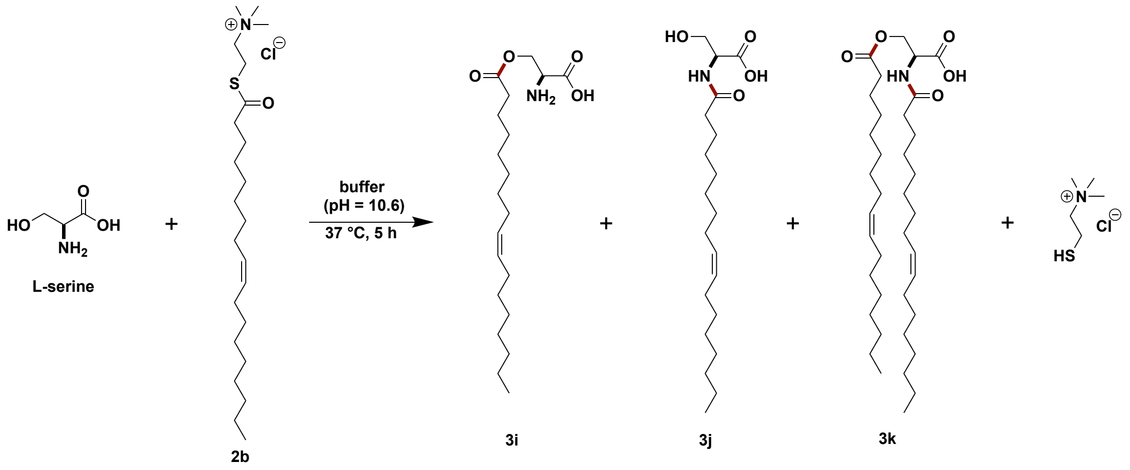


**Figure S4. Oleoylation reaction of L-serine.** To a mixture of L-serine (4 μmol) and oleoylation reagent **2b** (6 μmol), $Na_2CO_3/NaHCO_3$ buffer of pH = 10.6 (8 mL) was added. The reaction was tumbled at 37 °C for 5 hours. Acylated products **3i**, **3j**, **3k** were not detected.

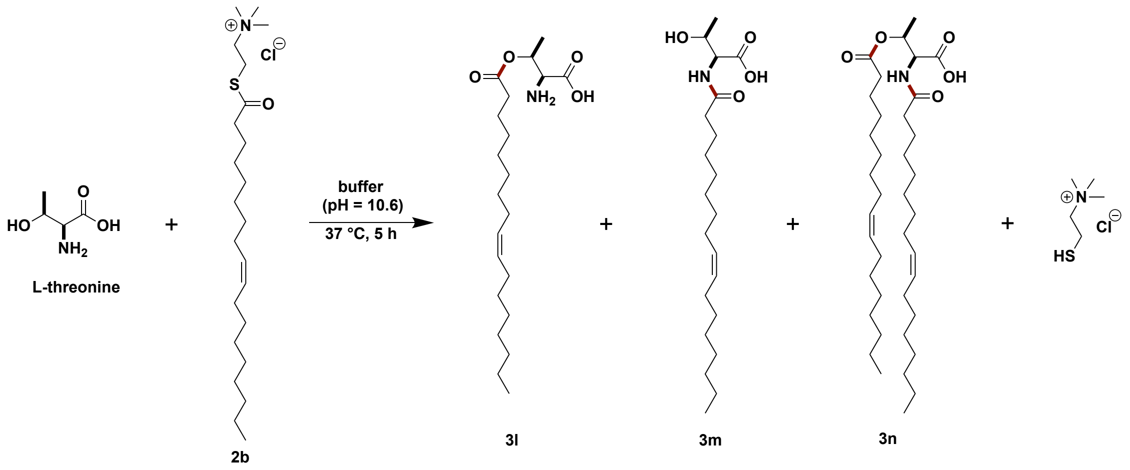


**Figure S5. Oleoylation reaction of L-threonine.** To a mixture of L-threonine (4 µmol) and oleoylation reagent **2b** (6 µmol), $Na_2CO_3/NaHCO_3$ buffer of pH = 10.6 (8 mL) was added. The reaction was tumbled at 37 °C for 5 hours. Acylated products **3l**, **3m**, **3n** were not detected.

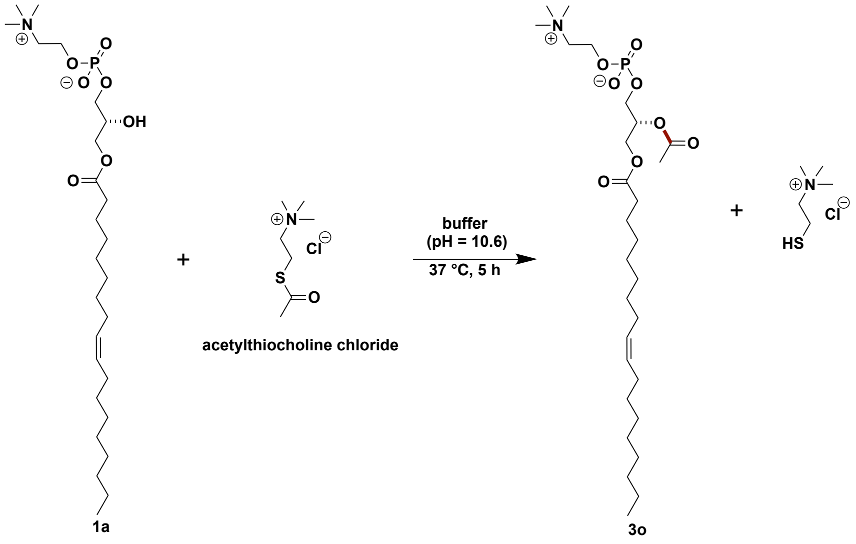


**Figure S6. Acetylation reaction of 1-oleoyl-2-hydroxy-*sn*-glycero-3-phosphocholine 1a.** To a mixture of lysophospholipid **1a** (4 µmol) and acetylthiocholine chloride (6 µmol), $Na_2CO_3/NaHCO_3$ buffer of pH = 10.6 (8 mL) was added. The reaction was tumbled at 37 °C for 5 hours. Less than 1% yield of acylated product **3o** was detected (HPLC yield).

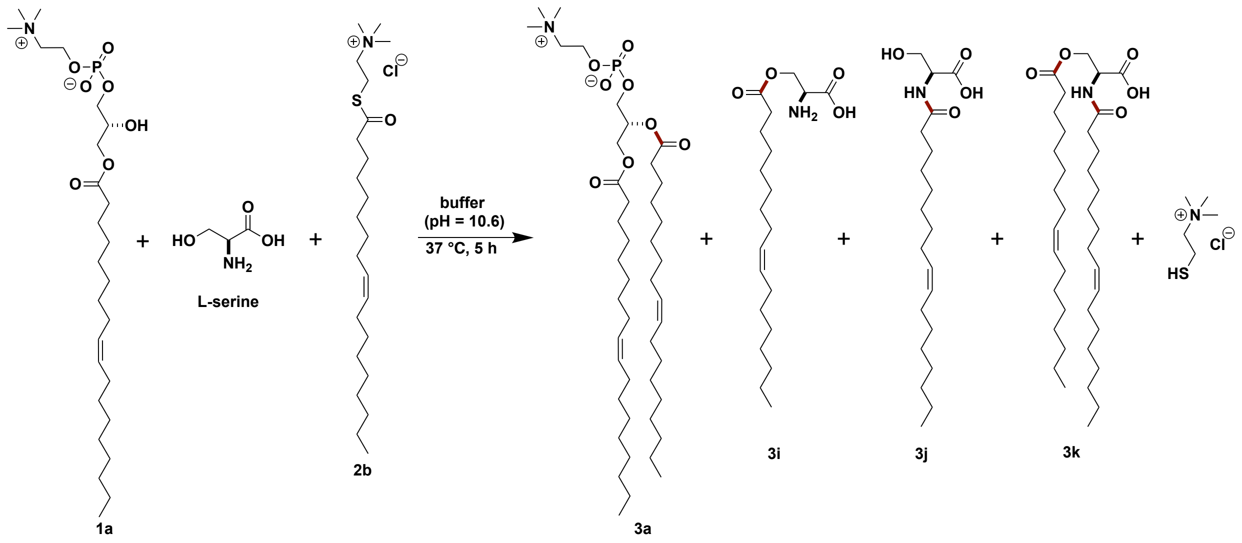


**Figure S7. Oleoylation reaction of 1-oleoyl-2-hydroxy-*sn*-glycero-3-phosphocholine 1a in the presence of L-serine.** To a mixture of lysophospholipid **1a** (4 µmol), L-serine (4 µmol) and oleoylation reagent **2b** (6 µmol), $Na_2CO_3/NaHCO_3$ buffer of pH = 10.6 (8 mL) was added. The reaction was tumbled at 37 °C for 5 hours. Acylated products **3i**, **3j**, **3k** were not detected, but an 88% yield of **3a** was obtained (HPLC yield).

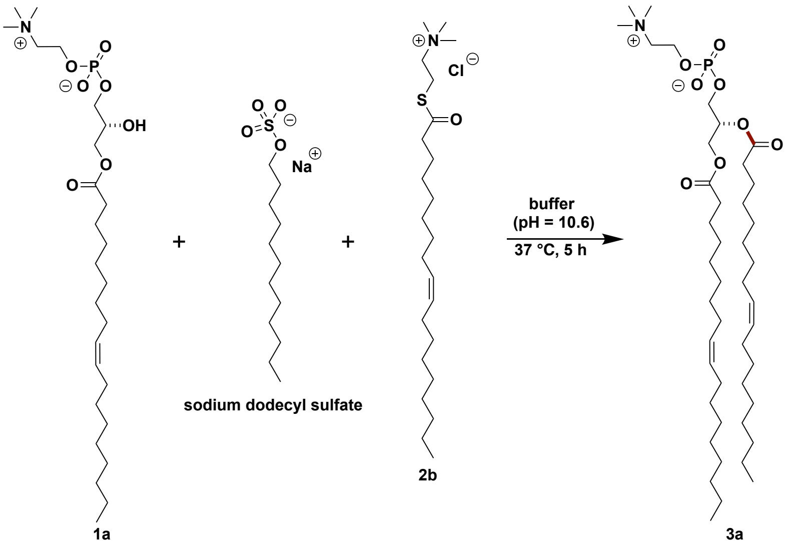


**Figure S8. Oleoylation reaction of 1-oleoyl-2-hydroxy-*sn*-glycero-3-phosphocholine 1a in the presence of sodium dodecyl sulfate.** To a mixture of lysophospholipid **1a** (4 µmol), sodium dodecyl sulfate (6 µmol) and oleoylation reagent **2b** (6 µmol), $Na_2CO_3/NaHCO_3$ buffer of pH = 10.6 (8 mL) was added. The reaction was tumbled at 37 °C for 5 hours. Less than 10% yield of **3a** was detected (HPLC yield).

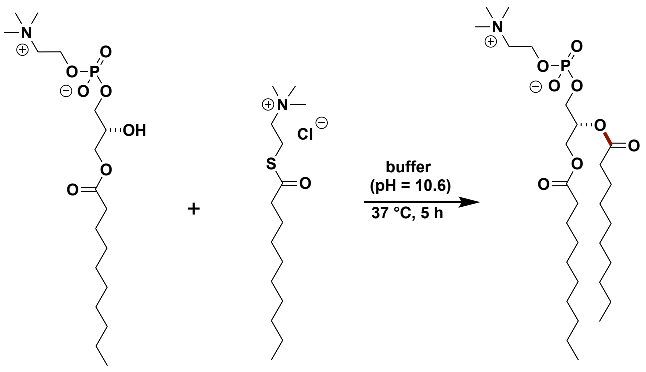


**1e** **2e** **3p**

**Figure S9. Decanoylation reaction of 1-decanoyl-2-hydroxy-*sn*-glycero-3-phosphocholine 1e. Condition A:** To a mixture of lysophospholipid **1e** (4 µmol) and decanoylation reagent **2e** (6 µmol), $Na_2CO_3/NaHCO_3$ buffer of pH = 10.6 (8 mL) was added. The reaction was tumbled at 37 °C for 5 hours. No obvious **3p** was obtained (LC-MC). **Condition B:** To a mixture of lysophospholipid **1e** (20 µmol) and decanoylation reagent **2e** (30 µmol), $Na_2CO_3/NaHCO_3$ buffer of pH = 10.6 (4 mL) was added. The reaction was tumbled at 37 °C for 5 hours. An isolated yield of 83% of **3p** was obtained using 40/60/2–80/20/2 MeOH/$CHCl_3$/$NH_3$•$H_2O$ as the eluents.

**1,2- Didecanoyl -*sn*-glycero-3-phosphocholine (3p)**

**$^1$H NMR** (500 MHz, $CDCl_3$): δ 5.20 (dq, *J* = 10.0 Hz, *J* = 5.0 Hz, 1 H), 4.39 (dd, *J* = 10.0 Hz, *J* = 5.0 Hz, 1 H), 4.37–4.34 (m, 2 H), 4.12 (q, *J* = 5.0 Hz, 1 H), 3.99–3.95 (m, 2 H), 3.88 (p, *J* = 5.0 Hz, 2 H), 3.39 (s, 9 H), 2.28 (dq, *J* = 5.0 Hz, *J* = 10.0 Hz, 4 H), 1.57 (dt, *J* = 5.0 Hz, *J* = 10.0 Hz, 4 H), 1.29–1.25 (m, 24 H), 0.87 (t, *J* = 5.0 Hz, 6 H).

**$^{13}$C NMR** (126 MHz, $CDCl_3$): δ 173.73, 173.37, 71.02, 70.96, 70.24, 70.17, 69.85, 69.79, 66.41, 63.81, 59.70, 59.66, 55.13, 54.69, 54.05, 34.44, 34.25, 32.02, 29.61, 29.46, 29.29, 25.08, 25.01, 22.82, 14.31, 14.25.

**HRMS** *m/z* (ESI): calcd. for $C_{28}H_{56}NO_8PNa$ $[M+Na]^+$: 588.3636; found: 588.3636.

### 6. In Situ Formation of Liposomes

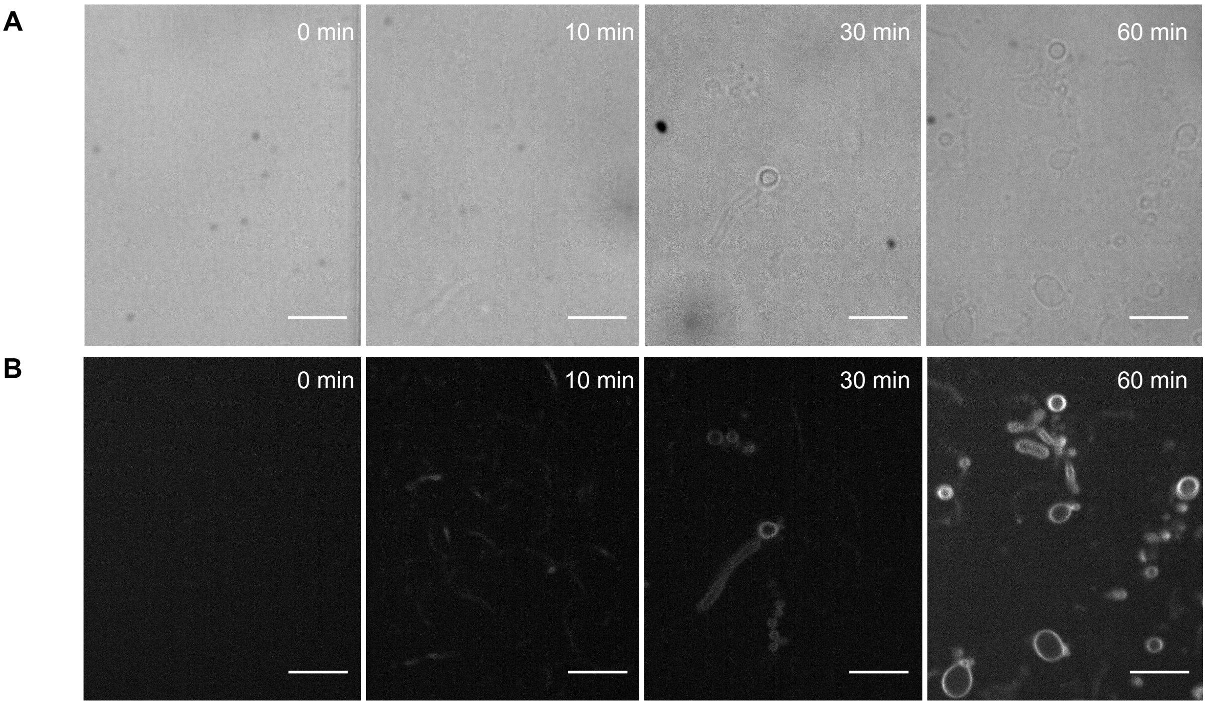


**Figure S10. Microscopy images of in vesicles formed during the reaction between 1-oleoyl-2-hydroxy-*sn*-glycero-3-phosphocholine 1a and 2-(oleoylthio)-*N*,*N*,*N*-trimethylethan-1-aminium chloride 2b in $Na_2CO_3$/$NaHCO_3$ buffer (pH = 8.8). A.** Brightfield micrographs are presented. **B.** Fluorescence micrographs are presented. The reaction was carried out by mixing lysophospholipid **1a** (0.5 mM) with 2-(oleoylthio)-*N*,*N*,*N*-trimethylethan-1-aminium chloride **2b** (0.75 mM) in the presence of $Na_2CO_3$/$NaHCO_3$ buffer (pH = 8.8) at 37 °C. The samples of the reaction mixture were taken at different time points and were stained by using 0.1 mol % Nile red dye. Scale bar: 10 µm.

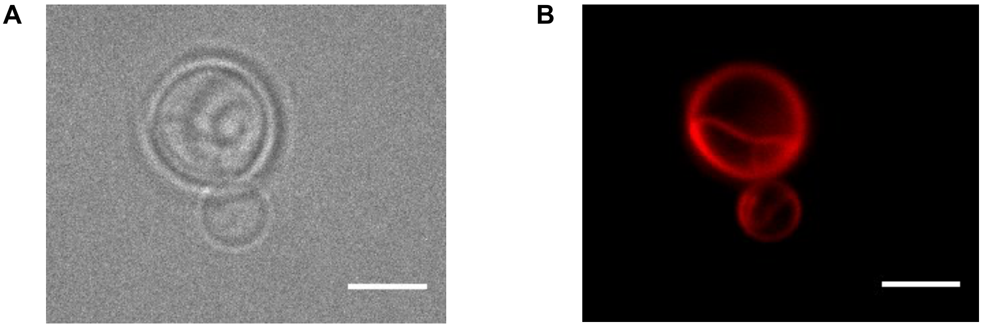


**Figure S11. Microscopy images of vesicles formed during the reaction between 1-oleoyl-2-hydroxy-*sn*-glycero-3-phosphocholine 1a and 2-(oleoylthio)-*N*,*N*,*N*-trimethylethan-1-aminium chloride 2b in Lost City vent fluid (pH = 9.1). A.** Brightfield micrographs are presented. **B.** Fluorescence micrographs are presented. The reaction was carried out by mixing lysophospholipid **1a** (0.5 mM) with 2-(oleoylthio)-*N*,*N*,*N*-trimethylethan-1-aminium chloride **2b** (0.75 mM) in the presence of Lost City vent fluid (pH = 9.1) at 37 °C. The sample of the reaction mixture was taken after 48 hours and stained by using 0.1 mol %

Nile red dye. Scale bar: 5 µm. Water samples from Lost City Hydrothermal Field (30°N, Mid-Atlantic Ridge) were taken by the Susan. Q. Lang group with the ROV Jason on September 22nd, 2018.

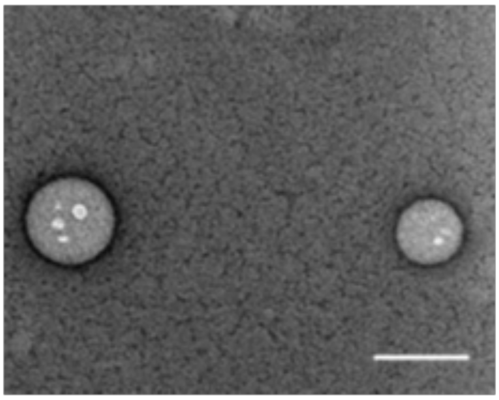

**Figure S12. TEM image vesicles formed during the reaction between 1-oleoyl-2-hydroxy-*sn*-glycero-3-phosphocholine 1a and 2-(oleoylthio)-*N*,*N*,*N*-trimethylethan-1-aminium chloride 2b in Mono Lake water (pH = 10).** The reaction was carried out by mixing lysophospholipid **1a** (0.5 mM) with 2-(oleoylthio)-*N*,*N*,*N*-trimethylethan-1-aminium chloride **2b** (0.75 mM) in the presence of Mono Lake water (pH = 10) at 37 °C. The samples of the reaction mixture were taken after 5 hours. After this, 4.8 µL of a reaction mixture was added to the formvar-coated Cu grid surface and allowed to sit for ~10 s and the excess solution was blotted into a filter paper. The grid was washed with 4.8 µL drops of $H_2O$ and subsequently stained with 4.8 µL drops of 2% uranyl acetate. The staining was carried out for ~30 s following which excess stain was blotted with filter paper. The grids were dried in air. After this, samples were imaged by transmission electron microscopy (TEM) using a FEI Tecnai Spirit G2 BioTWIN microscope operating at 80 kV and equipped with a bottom mount Eagle 4k (16 megapixel) camera. Negative staining transmission electron micrograph is presented. Scale bar: 200 nm. Water samples from Mono Lake, California were taken by Luping Liu and Ahanjit Bhattacharya on March 17th, 2019.

### 7. In Situ Encapsulation of HPTS And Proton Gradient Formation

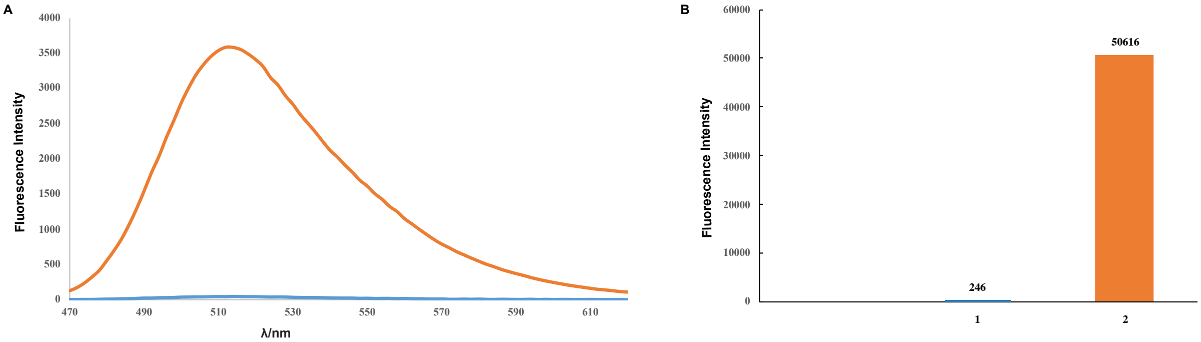


**Figure S13. A. Emission spectra of 8-hydroxypyrene-1,3,6-trisulfonic acid (HPTS).** Blue line: 8-hydroxypyrene-1,3,6-trisulfonic acid (HPTS) (0.25 mM) in the presence of citrate buffer (pH = 4.6). Orange line:8-hydroxypyrene-1,3,6-trisulfonic acid (HPTS) (0.25 mM) in the presence of Mono Lake water (pH = 10). **B. Fluorescence intensity of 8-hydroxypyrene-1,3,6-trisulfonic acid (HPTS).** 1. 8-hydroxypyrene-1,3,6-trisulfonic acid (HPTS) (0.25 mM) in the presence of citrate buffer (pH = 4.6). 2. 8-hydroxypyrene-1,3,6-trisulfonic acid (HPTS) (0.25 mM) in the presence of Mono Lake water (pH = 10). Excitation wavelength: 488 nm; emission wavelength: 514 nm.

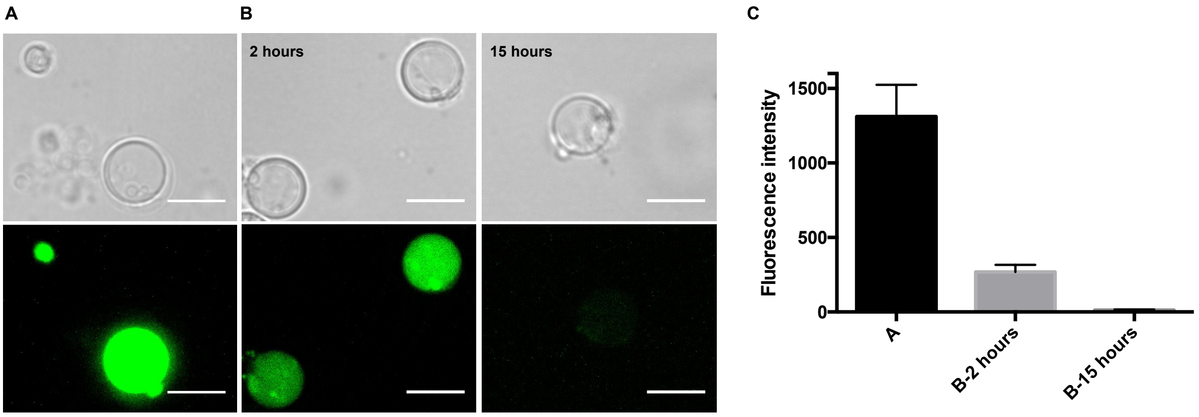


**Figure S14. pH gradient decay in the *de novo* formed vesicles. A.** The reaction was carried out by adding 8-hydroxypyrene-1,3,6-trisulfonic acid (HPTS) (0.25 mM) prior to the reaction of lysophospholipid **1a** (0.5 mM) with 2-(oleoylthio)-*N*,*N*,*N*-trimethylethan-1-aminium chloride **2b** (0.75 mM) in the presence of Mono Lake water (pH = 10). The reaction mixture was tumbled at 37 °C for 5 hours. **B.** The media of vesicles was exchanged to citrate buffer (pH = 4.6, in the same osmolarity as Mono Lake water) using spin-filtration, and a transmembrane pH gradient is spontaneously generated. Samples of the reaction mixture were taken at different time points to monitor the maintenance of pH gradient of the *de novo* formed vesicles. Scale bar: 10 µm. **Top Panels:** Brightfield micrographs are presented. **Bottom Panels:** Fluorescence micrographs are presented. **C.** Fluorescence intensity of encapsulated HPTS. The average background-subtracted fluorescence intensity was measured from 5 replicates. Error bars indicate SEM.

### 8. HPLC/ELSD Spectra

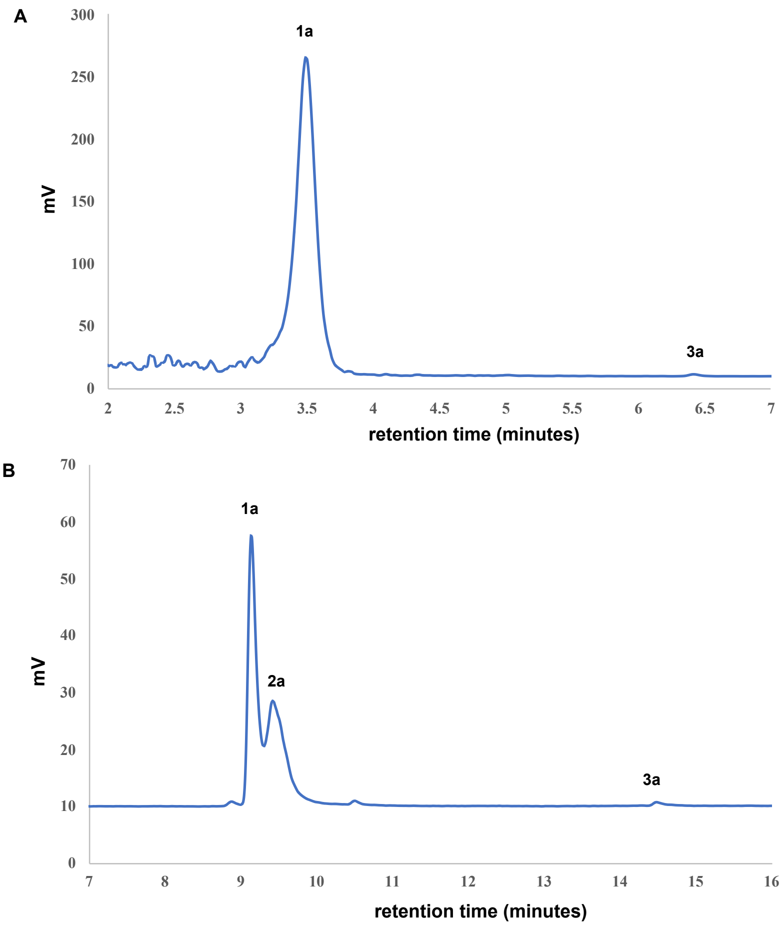


**Figure S15. HPLC/ELSD spectra of the oleoylation of 1a using acylation reagent 2a.** The reaction was carried out by mixing lysophospholipid **1a** (0.5 mM) with sodium 2-(oleoylthio)ethane-1-sulfonate **2a** (0.75 mM) in the presence of $Na_2CO_3$/$NaHCO_3$ buffer (pH = 10.6) at 37 °C. The sample of the reaction mixture was taken after 5 hours and was examined by HPLC/ELSD and LC-MS. Less than 1% of 1-oleoyl-2-hydroxy-*sn*-glycero-3-phosphocholine **1a** was converted to the desired product 1,2-dioleoyl-*sn*-glycero-3-phosphocholine **3a**.

**A.** HPLC analysis was carried out on an Eclipse Plus C8 analytical column with Phase A/Phase B gradients [Phase A: MeOH with 0.1% formic acid, Phase B: $H_2O$ with 0.1% formic acid]. 50%–95% Phase A in Phase B, 1 minute, and 95%–100% Phase A in Phase B, 5.5 minutes, then 100% Phase A, 0.5 minute. **B.** HPLC analysis was carried out on an Eclipse Plus C8 analytical column with Phase A/Phase B gradients [Phase A: MeOH with 0.1% trifluoroacetic acid, Phase B: $H_2O$ with 0.1% trifluoroacetic acid]. 50%–70% Phase A in Phase B, 1 minute, and 70%–90% Phase A in Phase B, 8 minutes, then 90%–95% Phase A in Phase B, 3 minutes, 100% Phase A, 4 minutes.

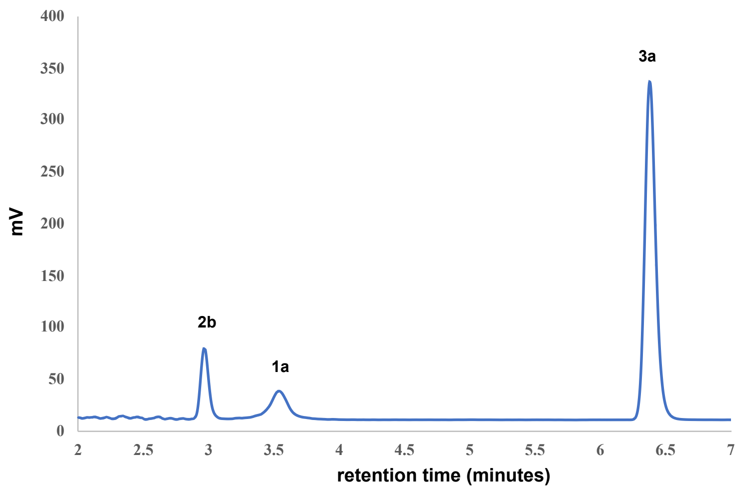


**Figure S16. HPLC/ELSD spectrum of the oleoylation of 1a using acylation reagent 2b.** The reaction was carried out by mixing lysophospholipid **1a** (0.5 mM) with 2-(oleoylthio)-*N*,*N*,*N*-trimethylethan-1-aminium chloride **2b** (0.75 mM) in the presence of $Na_2CO_3/NaHCO_3$ buffer (pH = 10.6) at 37 °C. The sample of the reaction mixture was taken after 5 hours and was examined by HPLC/ELSD and LC-MS. 88% yield of the desired product 1,2-dioleoyl-*sn*-glycero-3-phosphocholine **3a** was obtained.

HPLC analysis was carried out on an Eclipse Plus C8 analytical column with Phase A/Phase B gradients [Phase A: MeOH with 0.1% formic acid, Phase B: $H_2O$ with 0.1% formic acid]. 50%–95% Phase A in Phase B, 1 minute, and 95%–100% Phase A in Phase B, 5.5 minutes, then 100% Phase A, 0.5 minute.

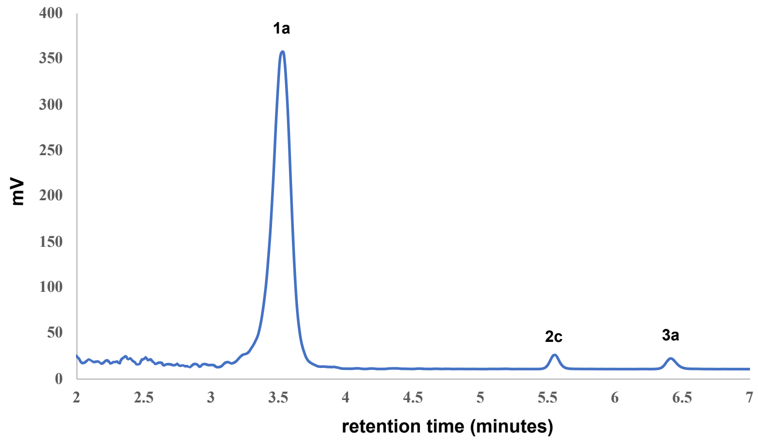


**Figure S17. HPLC/ELSD spectrum of the oleoylation of 1a using acylation reagent 2c.** The reaction was carried out by mixing lysophospholipid **1a** (0.5 mM) with S-propyl (*Z*)-octadec-9-enethioate **2c** (0.75 mM) in the presence of $Na_2CO_3/NaHCO_3$ buffer (pH = 10.6) at 37 °C. The sample of the reaction mixture was taken after 5 hours and was examined by HPLC/ELSD and LC-MS. Less than 1% of lysophospholipid **1a** was converted to the desired product 1,2-dioleoyl-*sn*-glycero-3-phosphocholine **3a**. More than 70% yield of **2c** was recovered, even though the ELSD signal of **2c** is poor.

HPLC analysis was carried out on an Eclipse Plus C8 analytical column with Phase A/Phase B gradients [Phase A: MeOH with 0.1% formic acid, Phase B: $H_2O$ with 0.1% formic acid]. 50%–95% Phase A in Phase B, 1 minute, and 95%–100% Phase A in Phase B, 5.5 minutes, then 100% Phase A, 0.5 minute.

### 9. $^{1}H$ and $^{13}C$ NMR Spectra

**2-(Oleoylthio)-*N*,*N*,*N*-trimethylethan-1-aminium chloride (2b)**

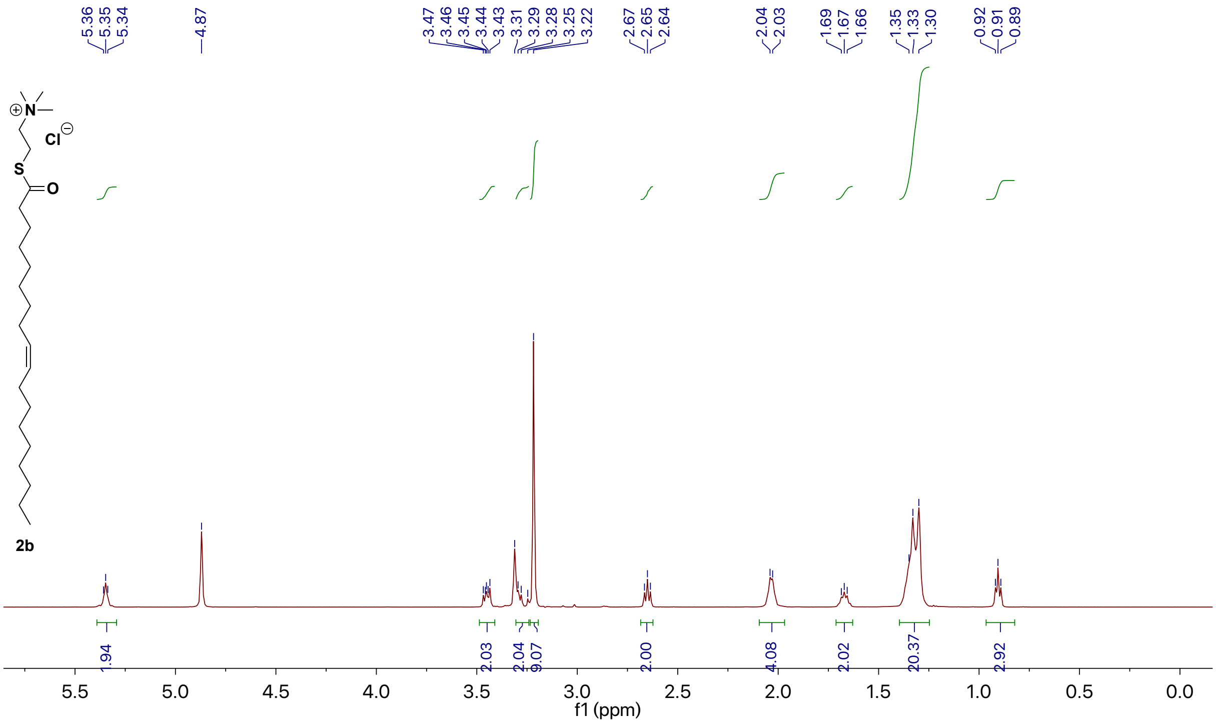


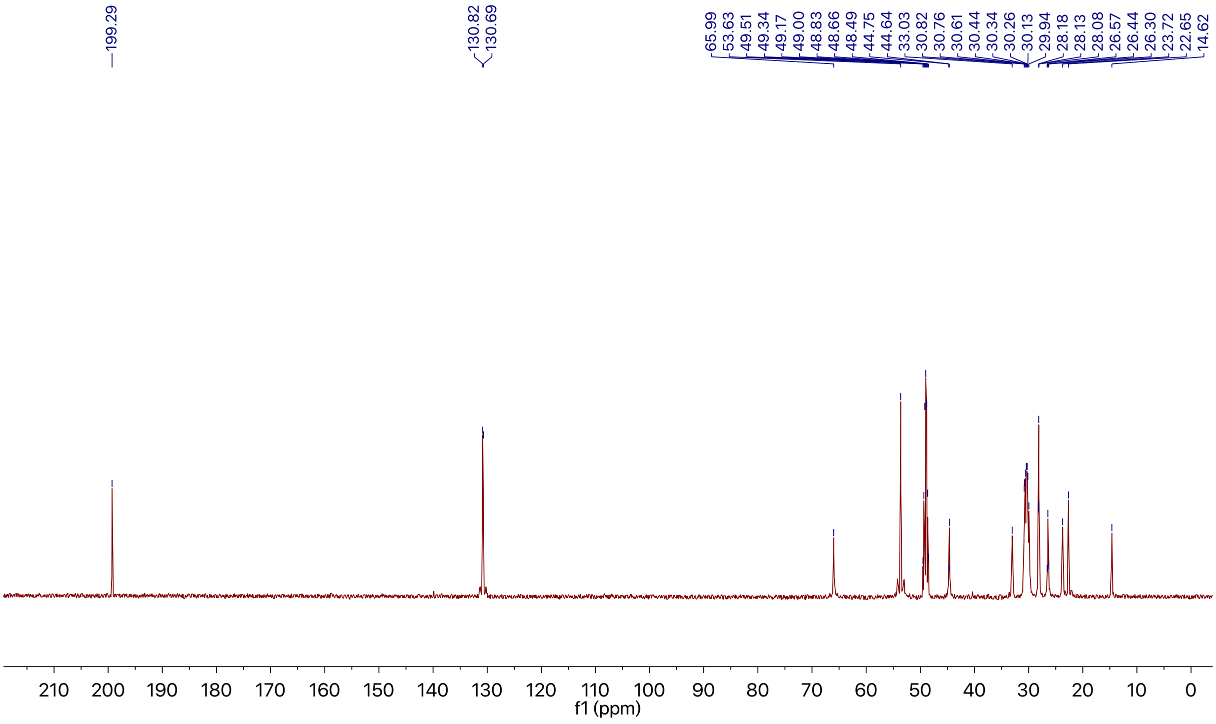

**S-propyl (*Z*)-octadec-9-enethioate (2c)**

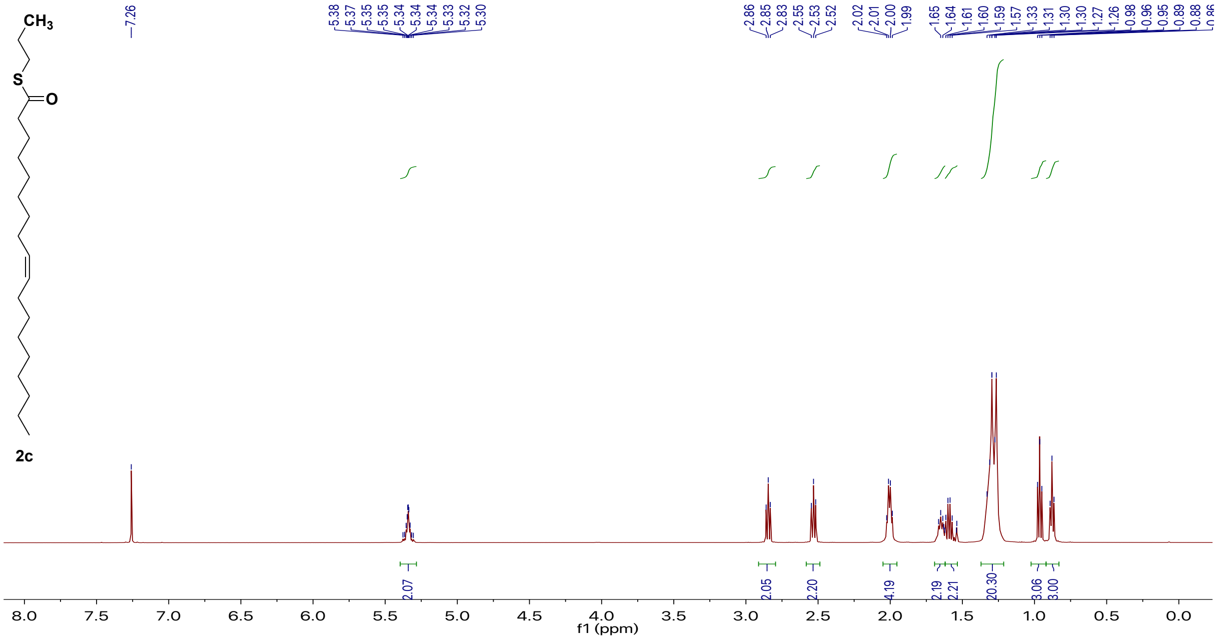


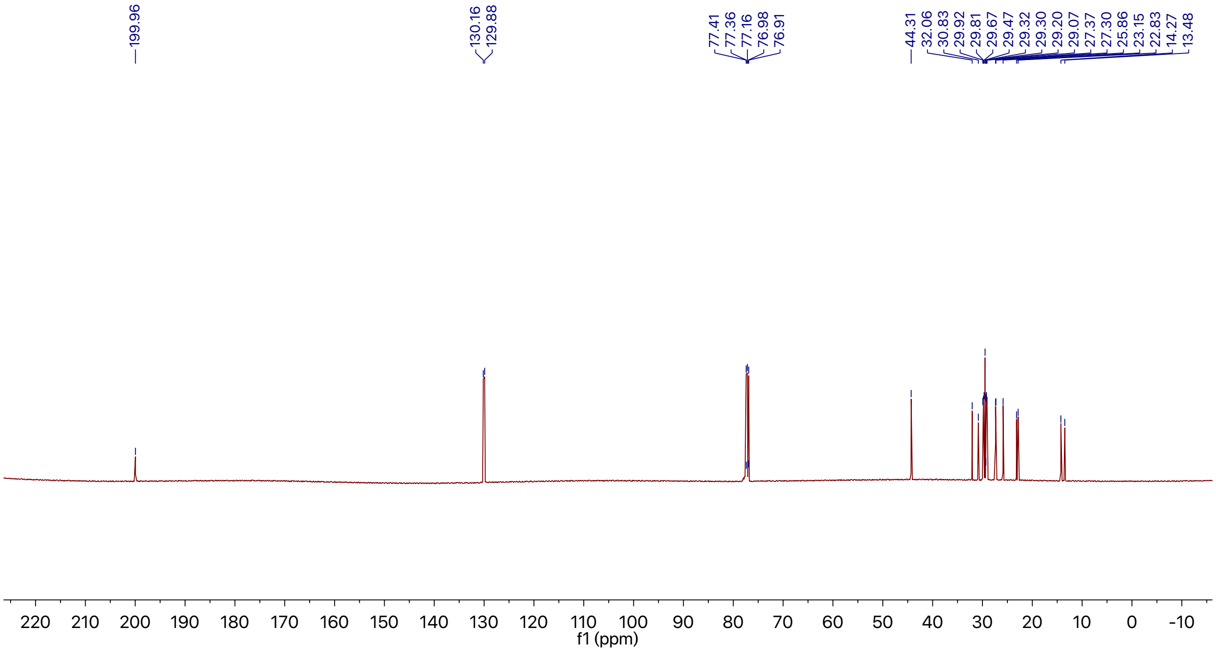

**2-(Palmitoylthio)-*N*,*N*,*N*-trimethylethan-1-aminium chloride (2d)**

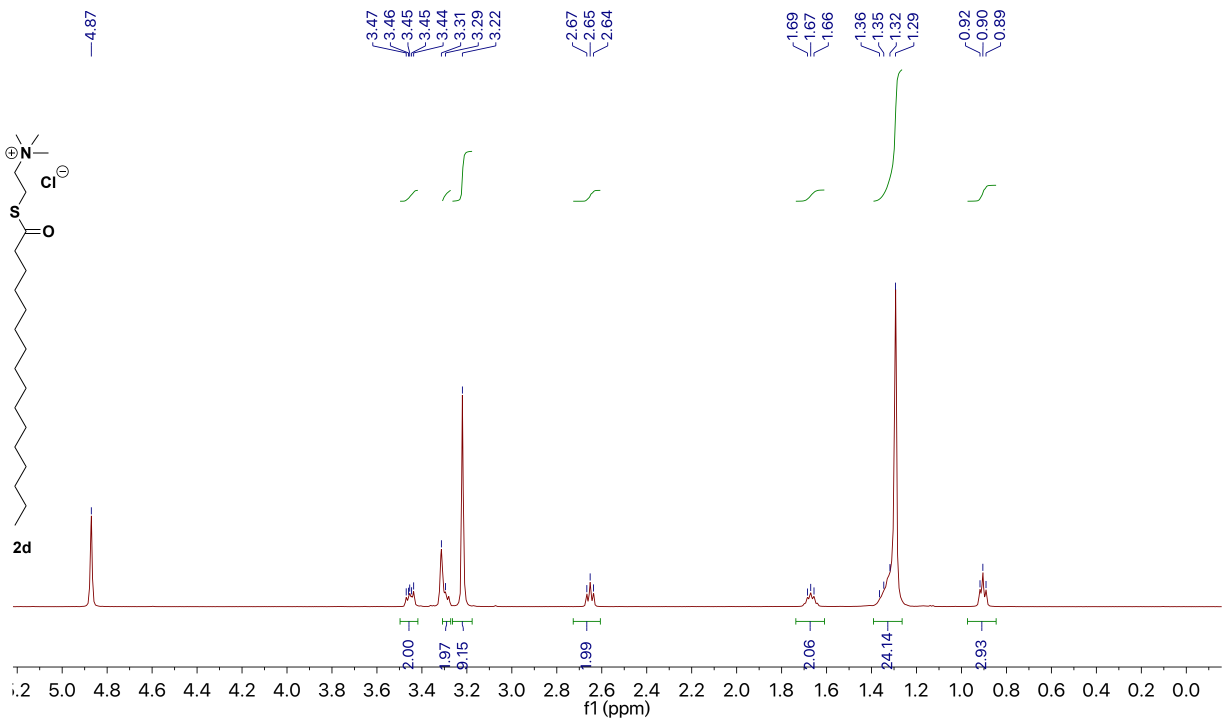

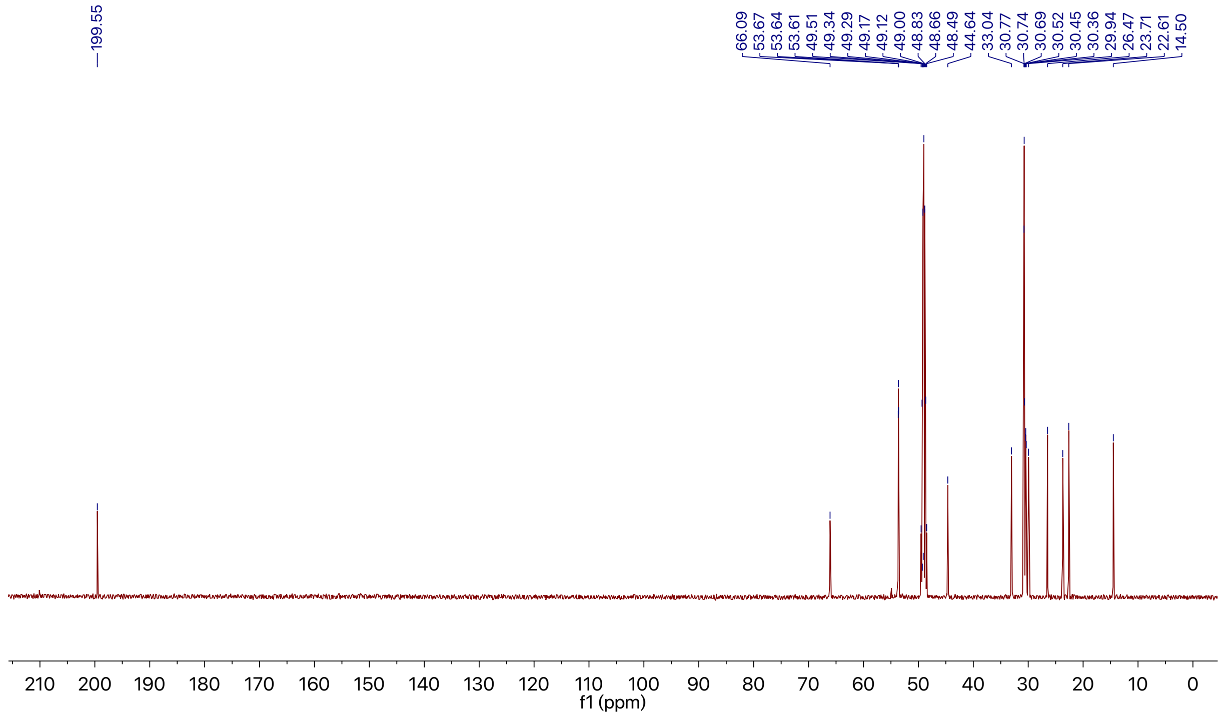

### 2-(Decanoylthio)-*N*,*N*,*N*-trimethylethan-1-aminium chloride (2e)

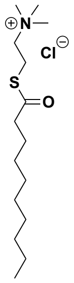


2e

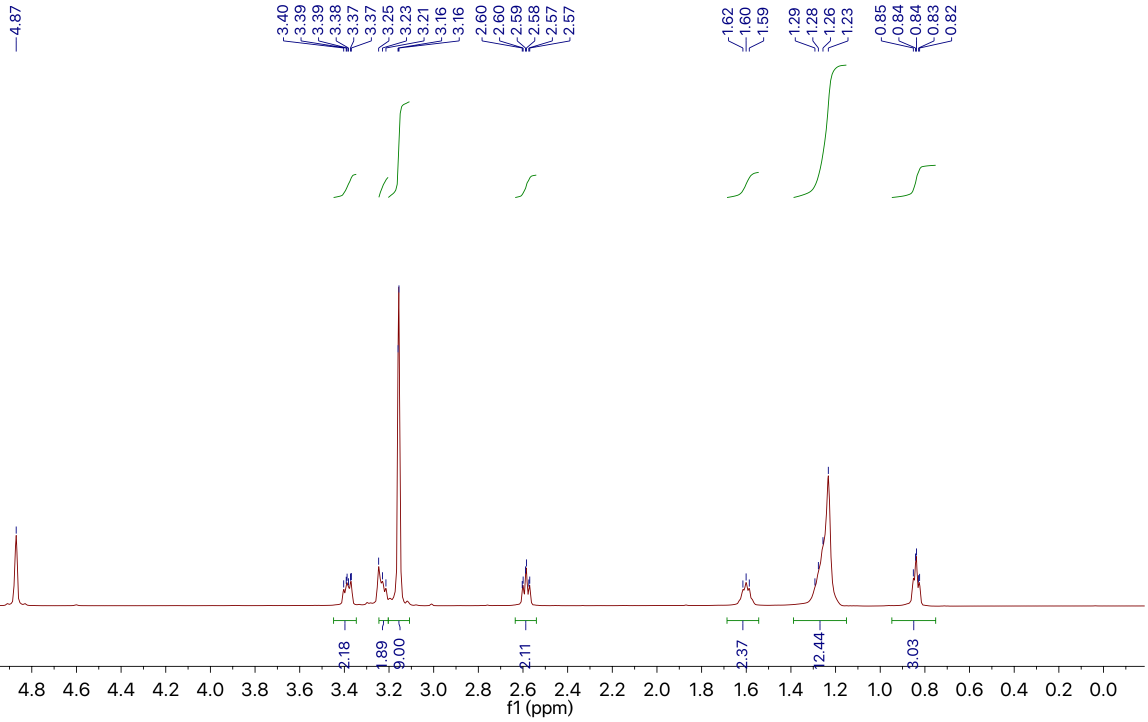


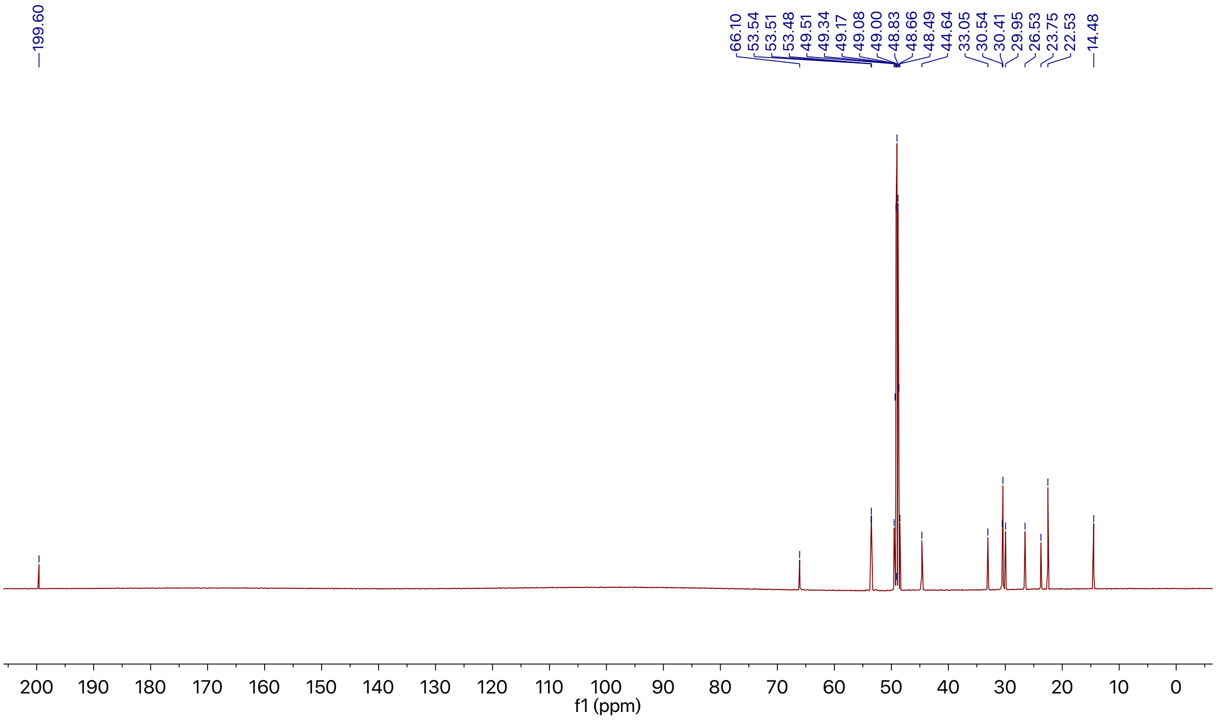

**1,2-Dioleoyl-*sn*-glycero-3-phosphocholine (3a)**

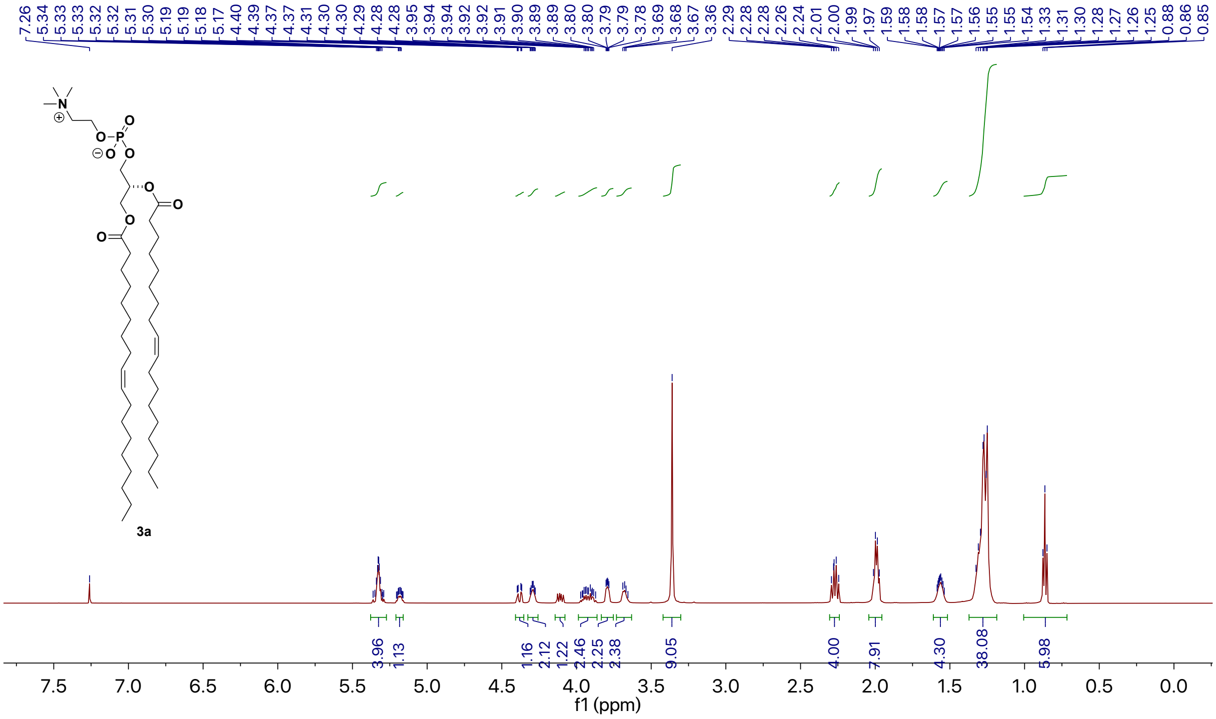


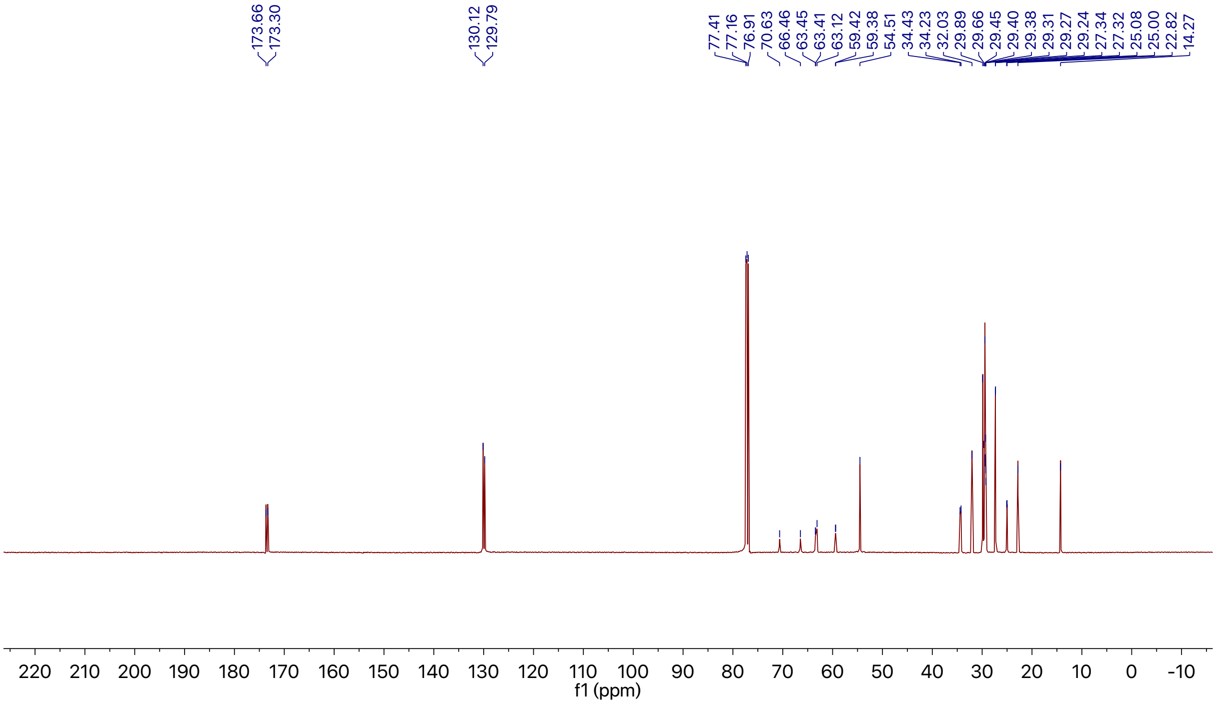

**1-Palmitoyl-2-oleoyl-*sn*-glycero-3-phosphocholine (3b)**

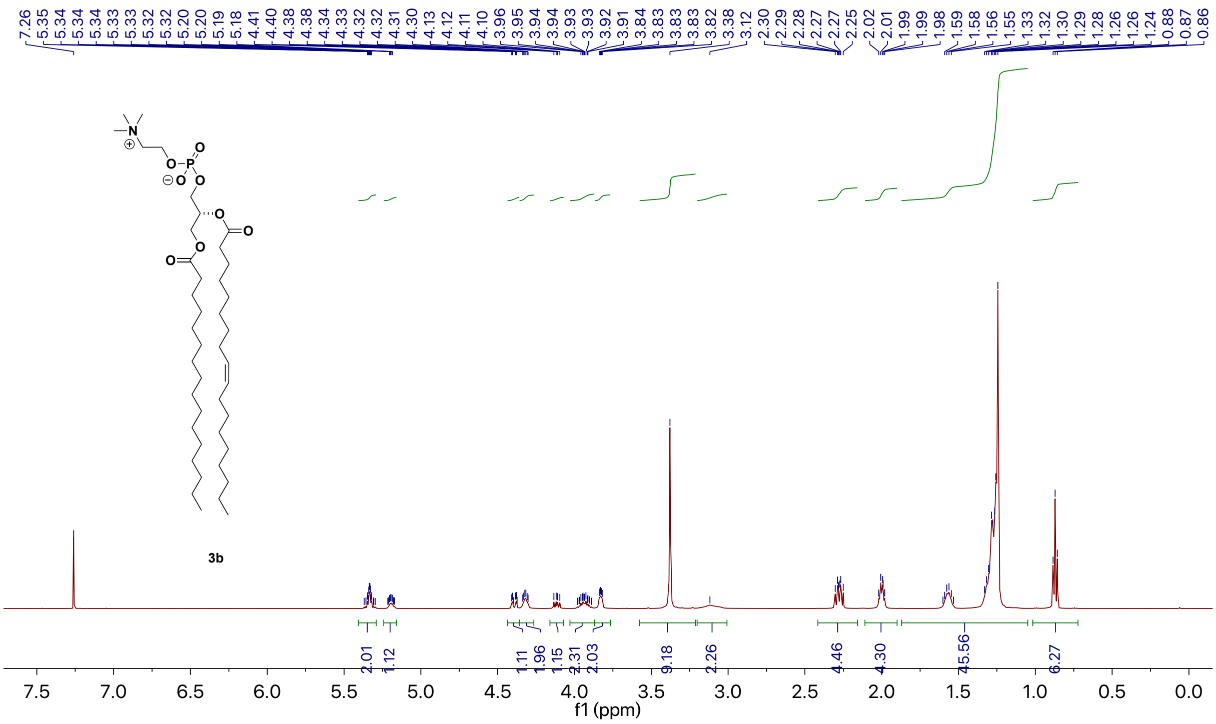


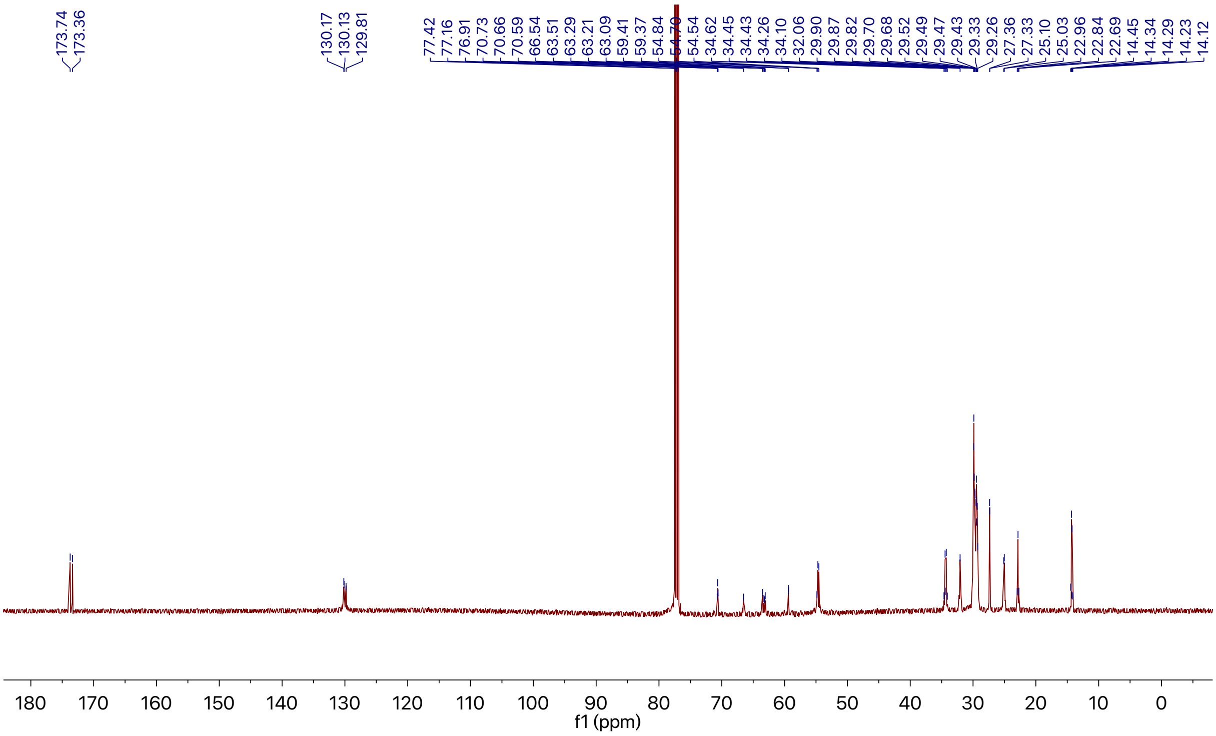

**1-Stearoyl-2-oleoyl-*sn*-glycero-3-phosphocholine (3c)**

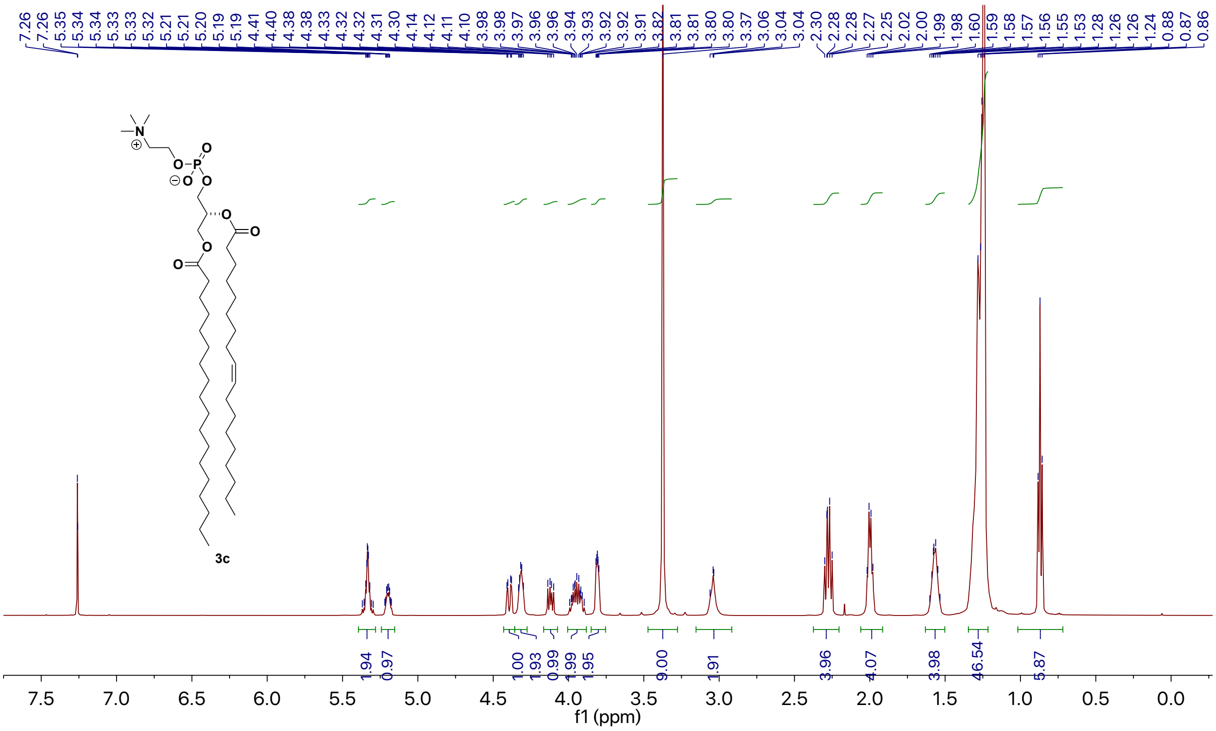


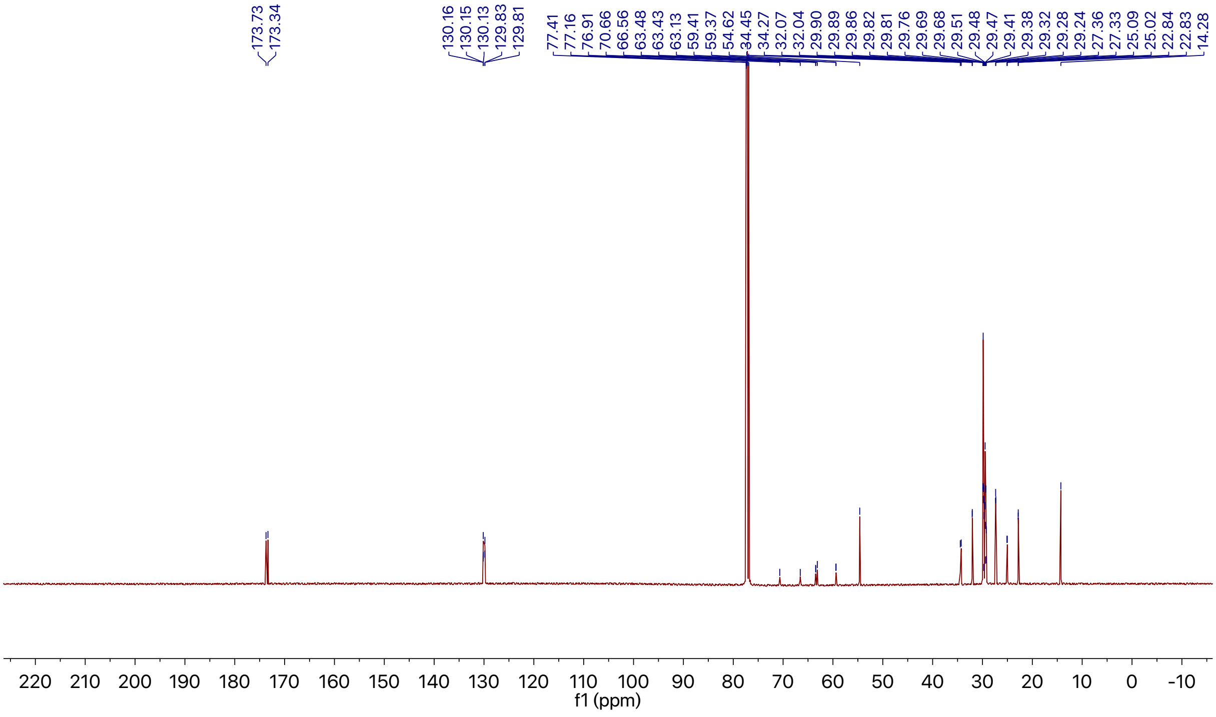

**1-Oleoyl-2-palmitoyl-*sn*-glycero-3-phosphocholine (3d)**

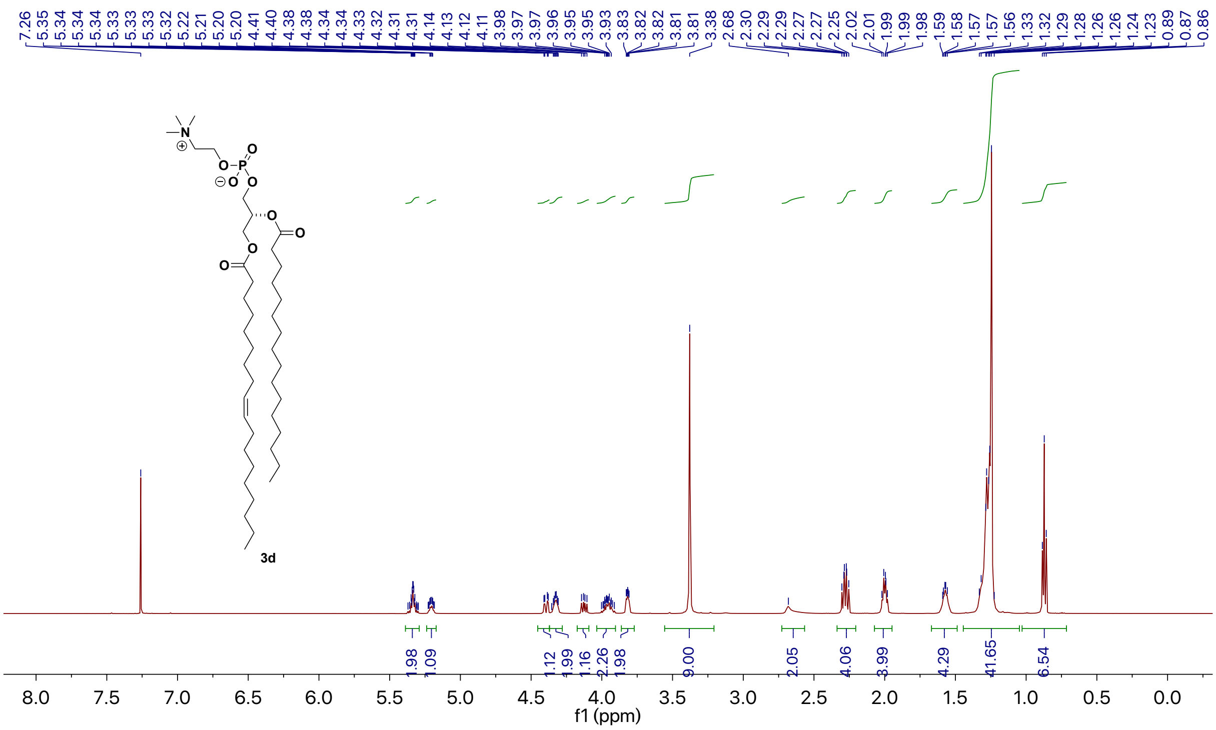


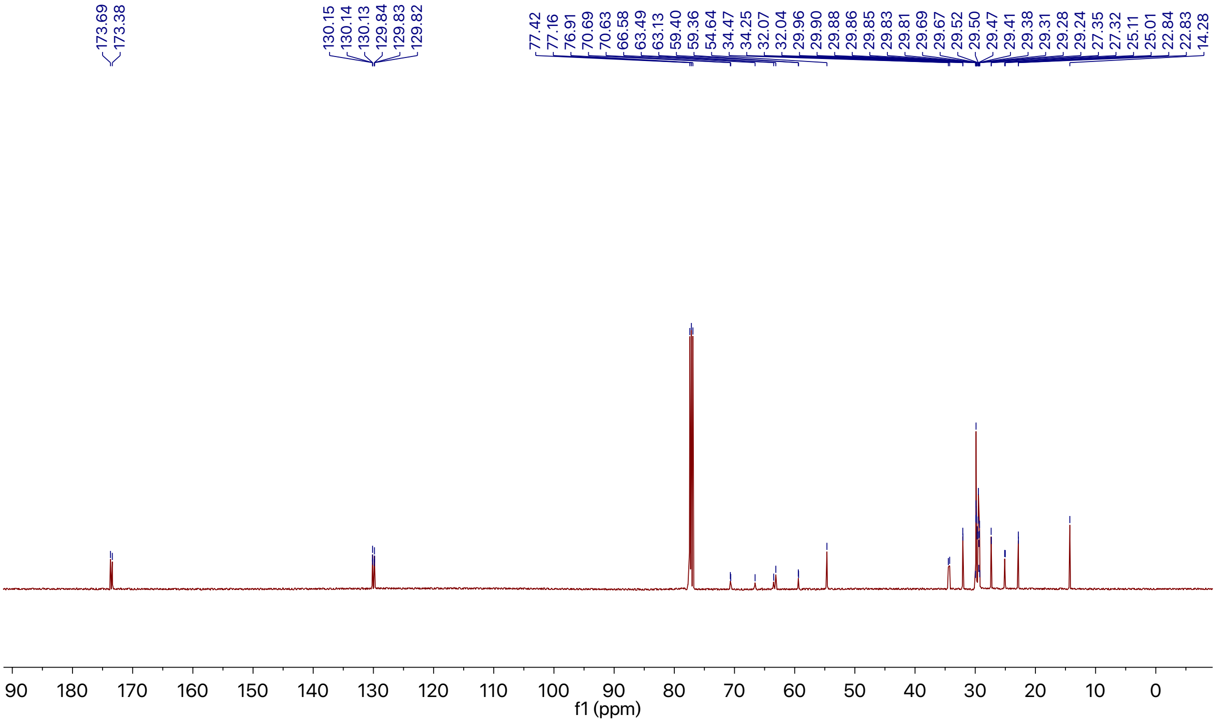

**1,2-Dipalmitoyl-*sn*-glycero-3-phosphocholine (3e)**

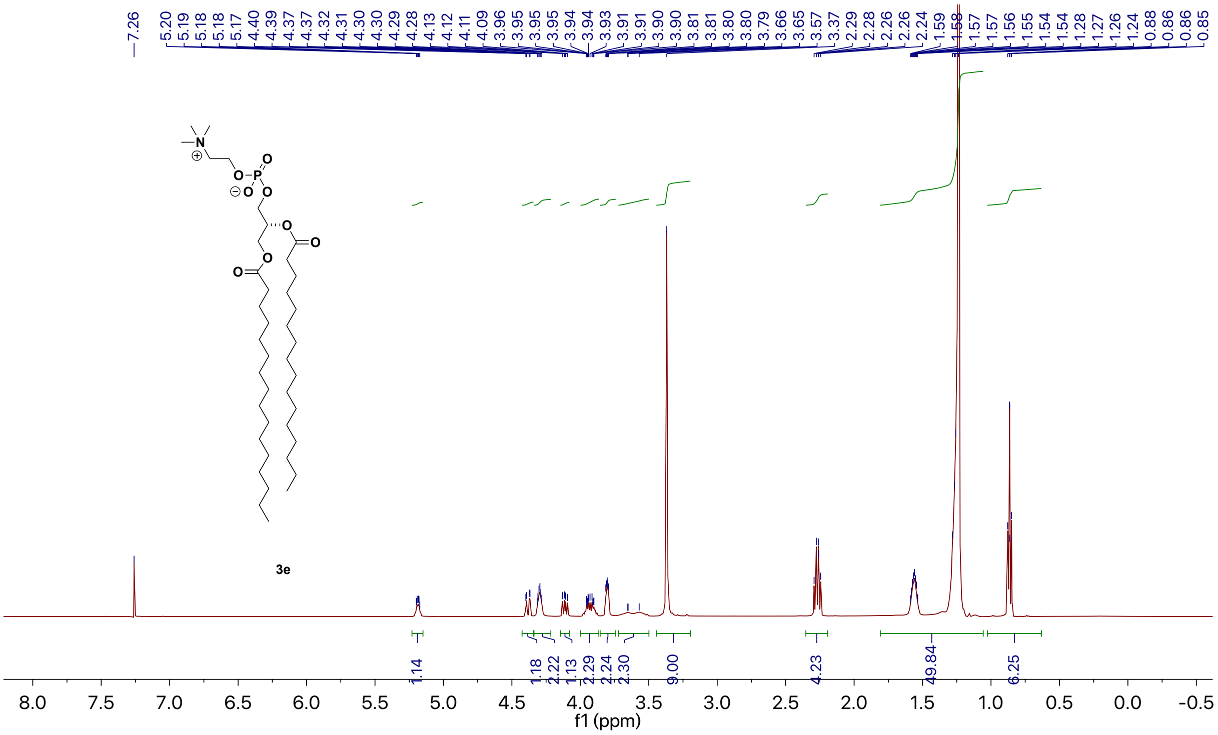

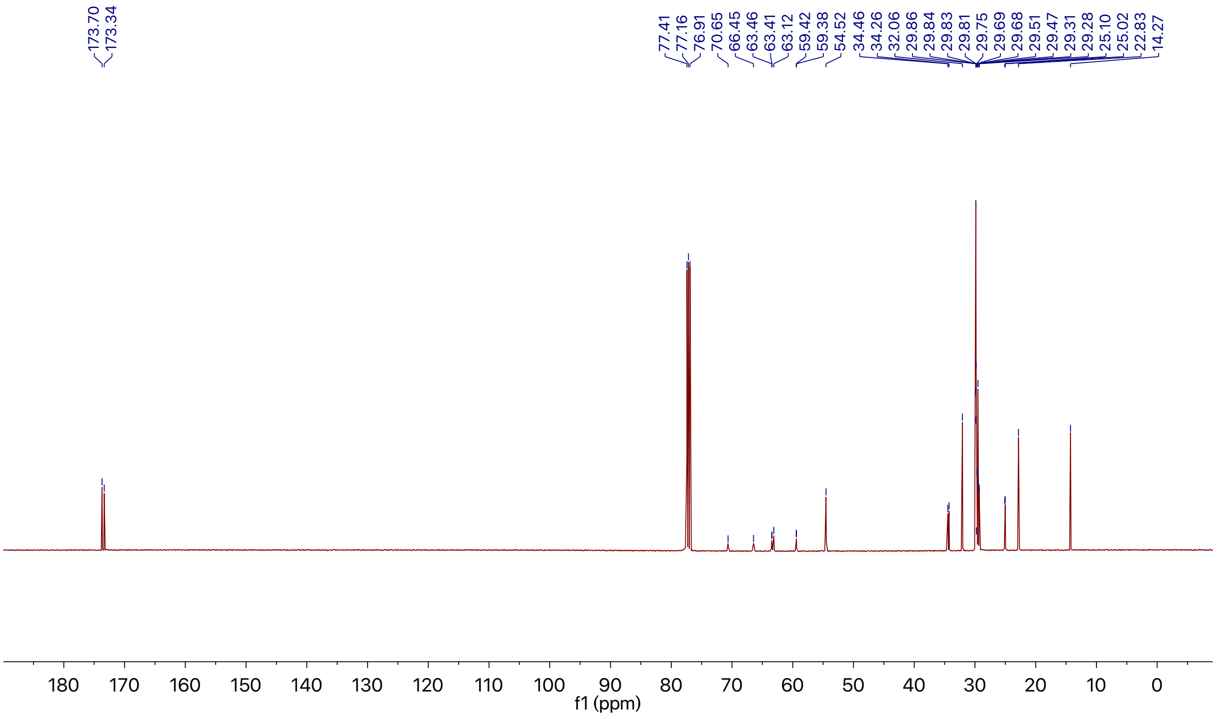

**1-Stearoyl-2-palmitoyl-*sn*-glycero-3-phosphocholine (3f)**

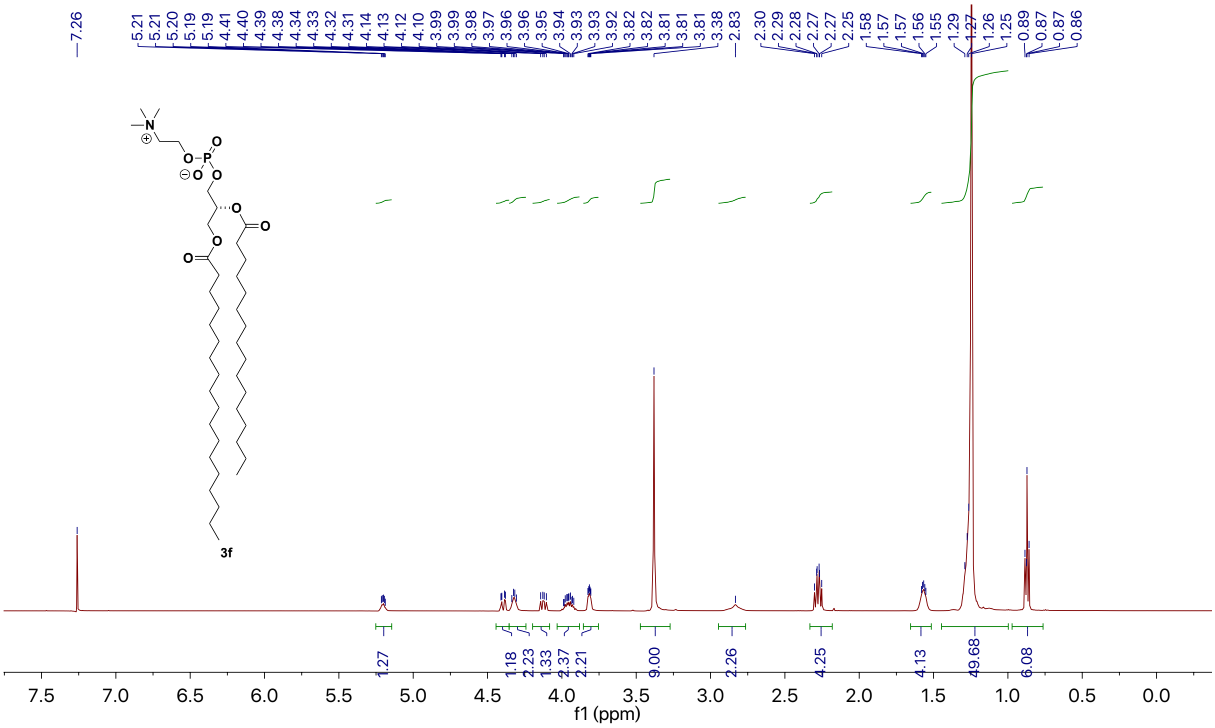

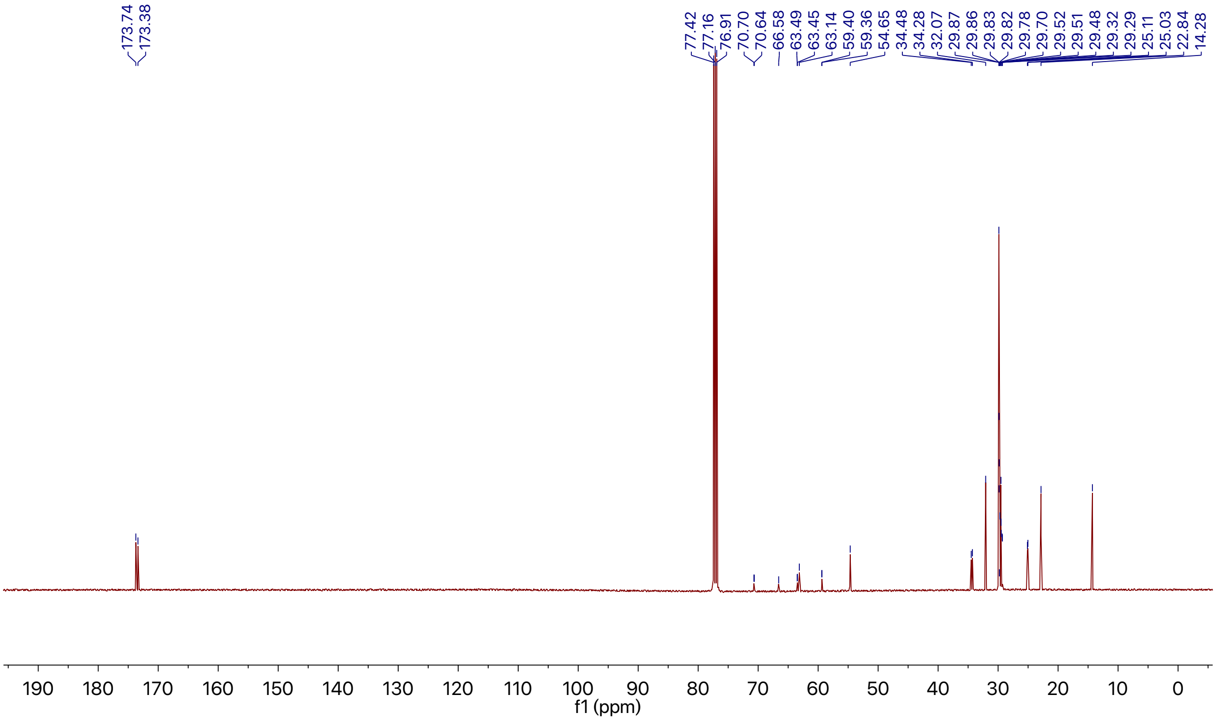

**1,2-Dioleoyl-*sn*-glycero-3-phosphate sodium salt (3g)**

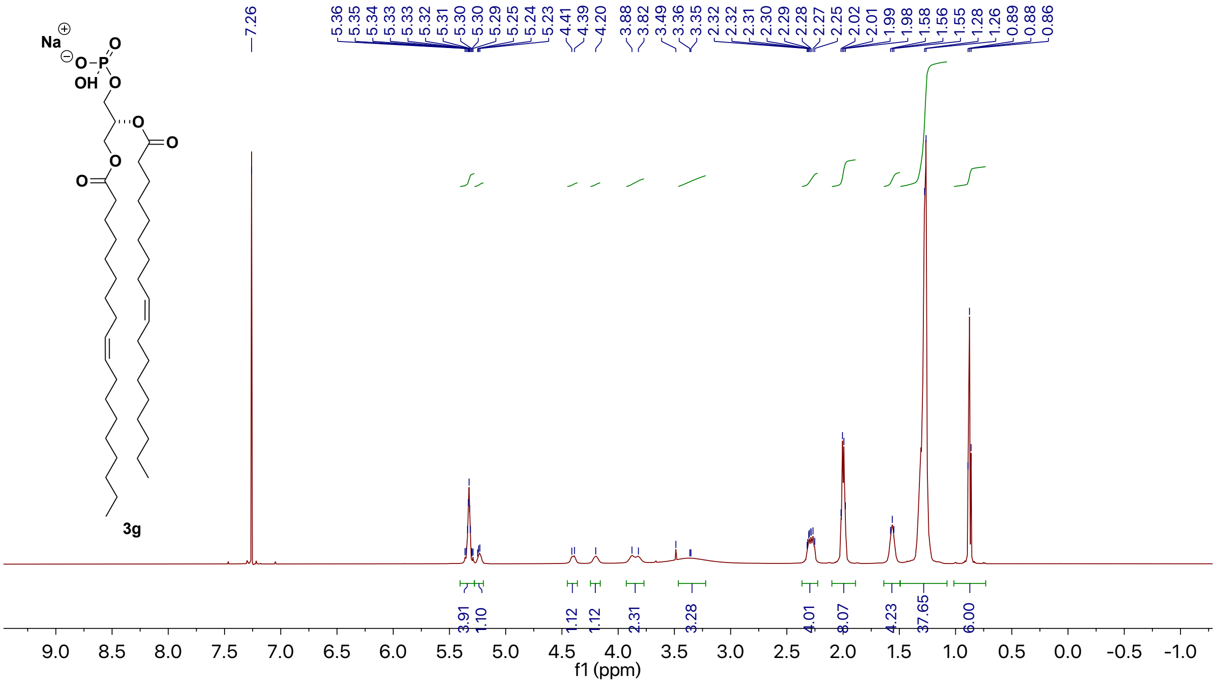


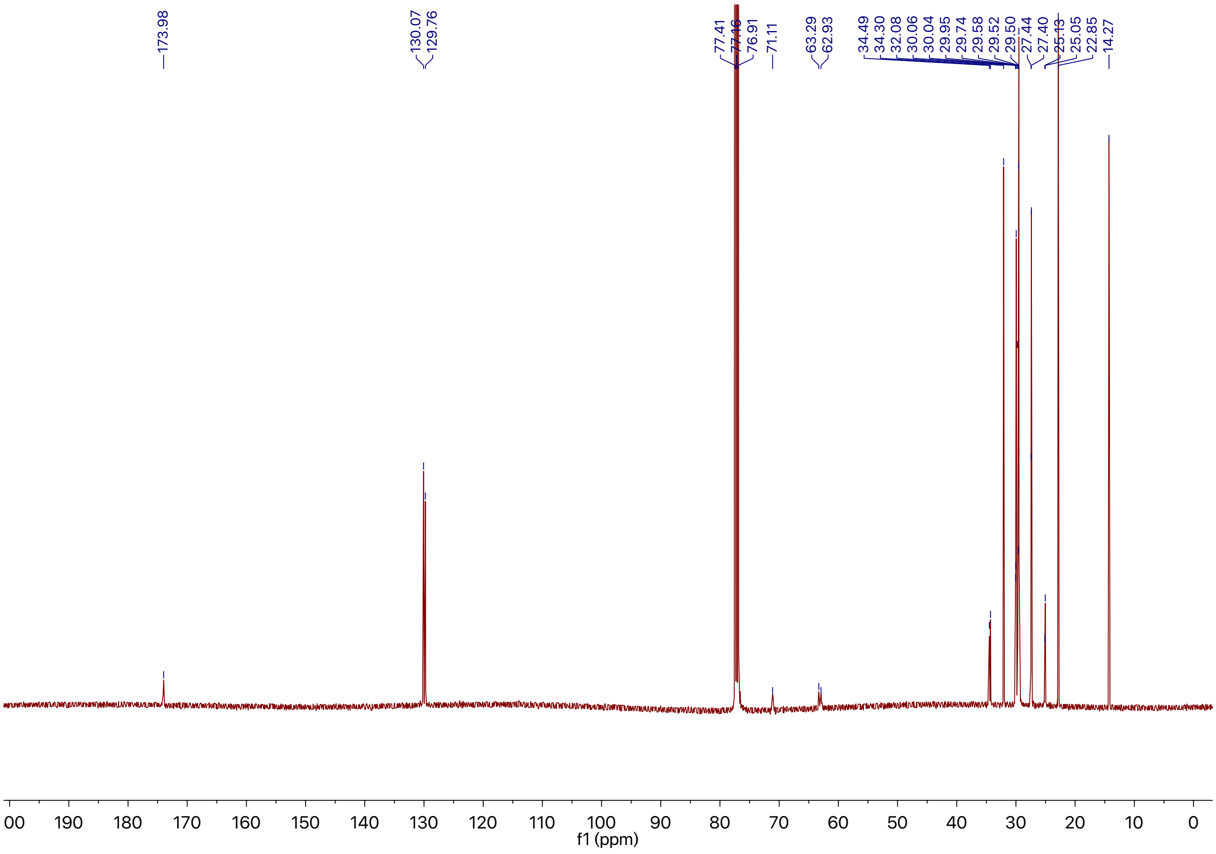

**1,2- Didecanoyl-*sn*-glycero-3-phosphocholine (3p)**

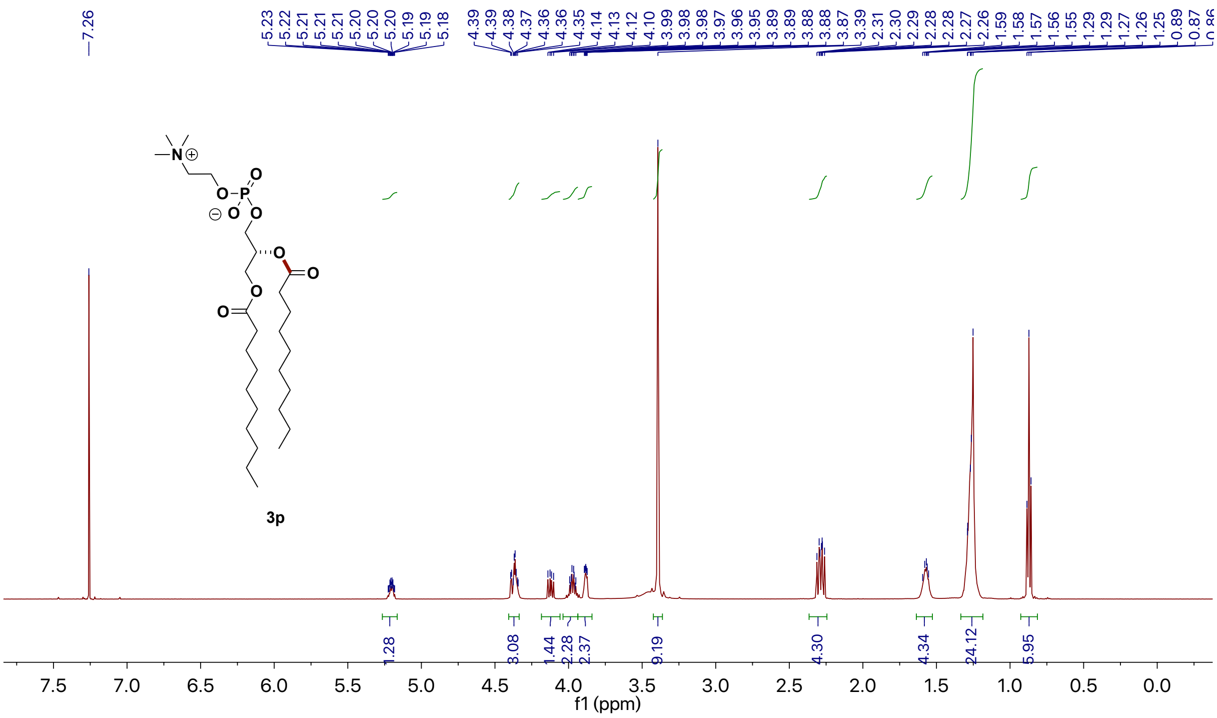

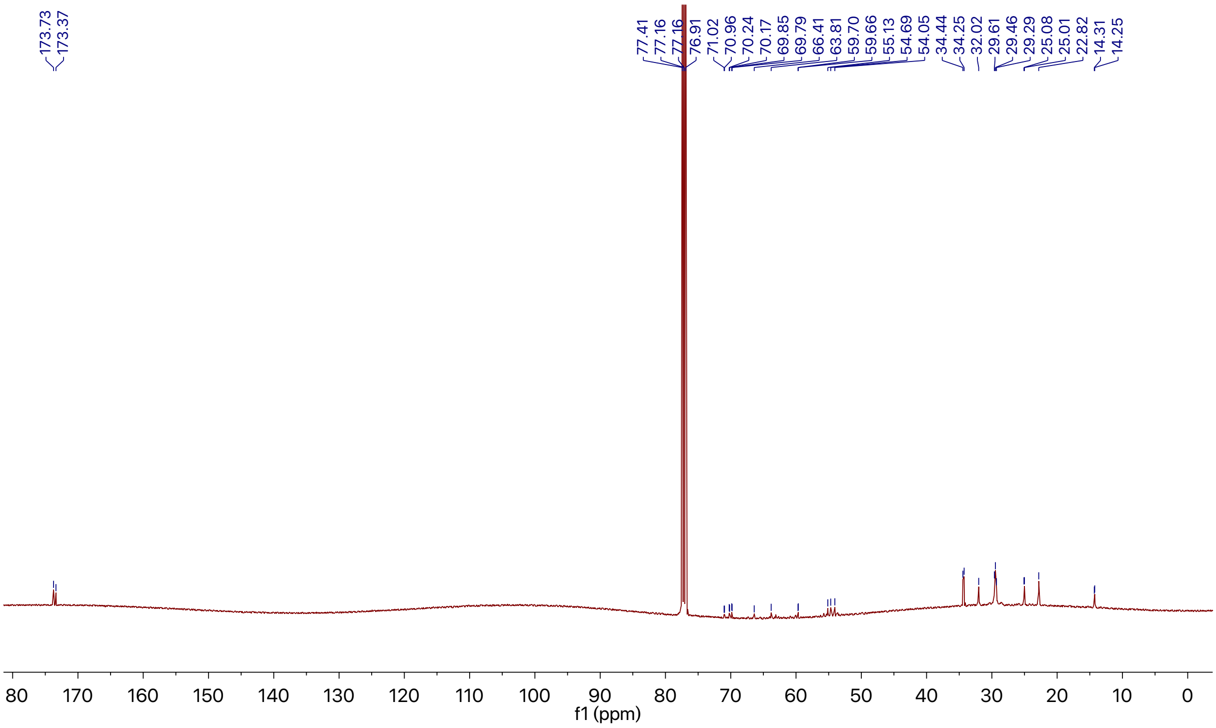

### 10. Theoretical Study

#### General computation procedure

All calculations were performed using Gaussian 09.[20] The initial structure of each species was built in such a way that the chain was in an anti-periplanar conformation to minimize steric interactions. For each species of **INT** (**1~4**), the initial structure was generated by rotating O2-C1' bond in such a way that an internal hydrogen bond was formed between C1'-OH and C1-O, while the chain was in an anti-periplanar conformation. We assumed these structures to be the representative low energy conformations. The representative low energy conformation of each species was then used for geometry optimization at the B3LYP/6-31G(d,p)/SMD(water) level. A frequency calculation was performed on each optimized structure at the same theoretical level to verify that an energy minimum was obtained and to determine frequencies for the free energy calculations. Single point energy calculations were then conducted at D3-B3LYP/6-311++G(2d,p)/SMD(water) level. All reported Gibbs free energies are for 298 K.

**The optimized representative low energy conformations and relative Gibbs free energies**

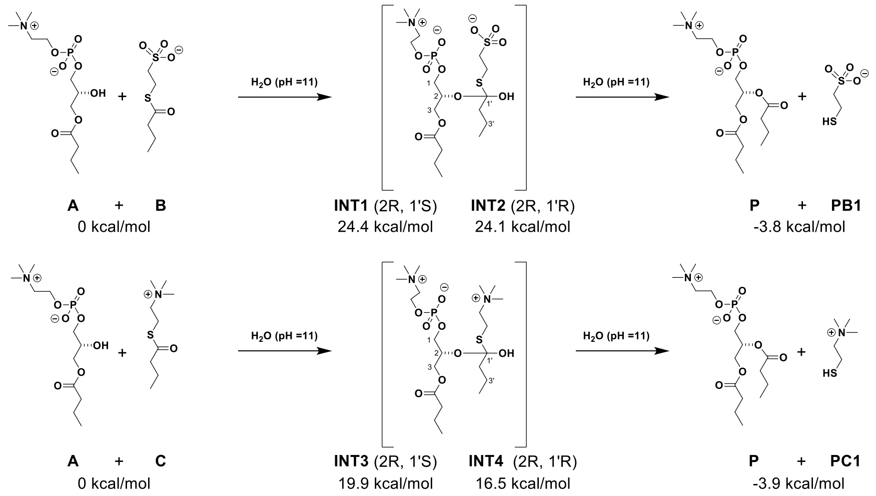


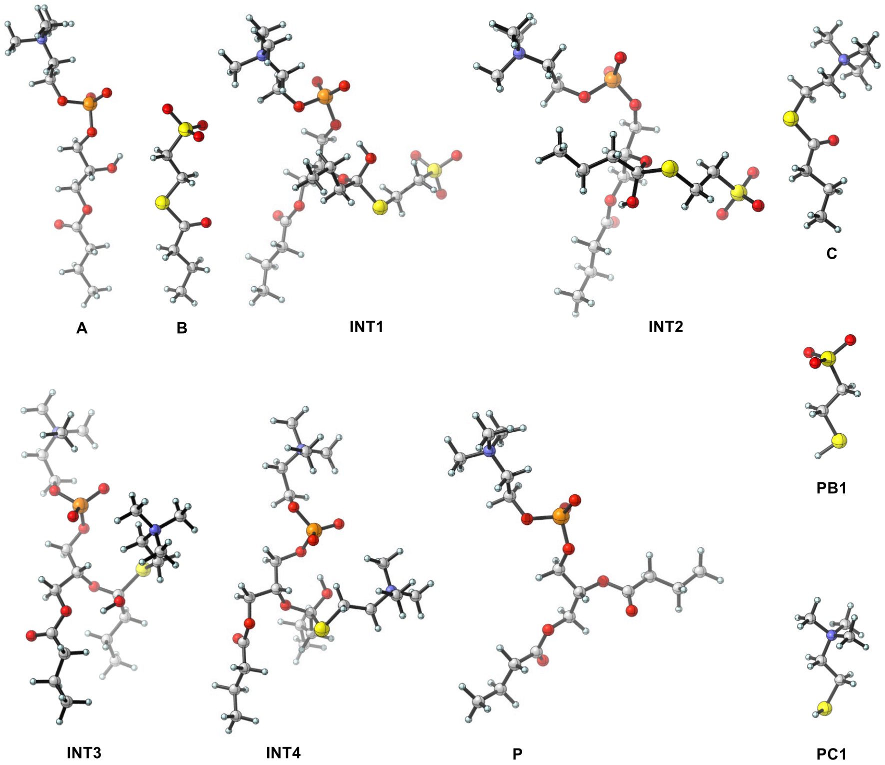

**Cartesian Coordinates**

| A | | | |
|---|---|---|---|
| N | -6.107100 | 0.867500 | -0.180200 |
| C | -4.720400 | 0.434700 | -0.619100 |
| C | -3.583000 | 0.773400 | 0.348200 |
| O | -2.366200 | 0.296300 | -0.243900 |
| O | -1.799300 | -1.379100 | 1.638700 |
| P | -1.831900 | -1.230000 | 0.136400 |
| O | -0.323500 | -1.071500 | -0.491200 |
| O | 4.331900 | -0.083300 | -0.179300 |
| C | 0.594500 | -0.117100 | 0.076900 |
| C | 3.035900 | 0.351100 | 0.298800 |
| C | 1.999500 | -0.598100 | -0.267400 |
| C | -7.062800 | 0.524900 | -1.292400 |
| C | -6.534800 | 0.135000 | 1.063100 |
| C | -6.161200 | 2.350900 | 0.064600 |
| O | -2.526000 | -2.260400 | -0.718800 |
| O | 2.234500 | -1.895300 | 0.280000 |
| O | 5.277900 | 1.680900 | 0.864900 |
| C | 5.389000 | 0.672800 | 0.184800 |
| C | 6.683600 | 0.102800 | -0.339800 |
| C | 7.895600 | 0.997800 | -0.077900 |
| C | 9.193100 | 0.378200 | -0.598000 |
| H | -4.768200 | -0.644300 | -0.772000 |
| H | -4.536100 | 0.920700 | -1.578800 |
| H | -3.468200 | 1.852100 | 0.470200 |
| H | -3.726500 | 0.323900 | 1.334100 |
| H | 0.404600 | 0.872800 | -0.350300 |
| H | 0.466000 | -0.074000 | 1.163400 |
| H | 3.030500 | 0.329400 | 1.392300 |
| H | 2.847800 | 1.373300 | -0.039100 |
| H | 2.100700 | -0.624100 | -1.362600 |
| H | -8.067700 | 0.812000 | -0.984100 |
| H | -6.769000 | 1.074800 | -2.186100 |
| H | -7.014200 | -0.548700 | -1.473800 |
| H | -6.460800 | -0.936000 | 0.875600 |
| H | -5.891300 | 0.421900 | 1.892800 |
| H | -7.566400 | 0.410100 | 1.282200 |
| H | -5.545000 | 2.598100 | 0.927300 |
| H | -5.796300 | 2.863300 | -0.825400 |

| | | | |
|---|---|---|---|
| H | -7.197200 | 2.625200 | 0.261900 |
| H | 1.504600 | -2.453900 | -0.029900 |
| H | 6.815200 | -0.881700 | 0.128100 |
| H | 6.556200 | -0.091900 | -1.411400 |
| H | 7.732900 | 1.973200 | -0.551200 |
| H | 7.978000 | 1.186300 | 0.998400 |
| H | 10.047100 | 1.034600 | -0.402300 |
| H | 9.395400 | -0.585500 | -0.117000 |
| H | 9.146800 | 0.205400 | -1.679300 |

| B | | | |
|---|---|---|---|
| C | 6.056600 | -0.008600 | -0.032000 |
| C | 4.659500 | 0.598600 | -0.163900 |
| C | 3.560600 | -0.417800 | 0.161200 |
| C | 2.163800 | 0.158700 | 0.069000 |
| S | 0.864600 | -1.084900 | -0.036800 |
| O | 1.909900 | 1.351300 | 0.087200 |
| C | -0.619300 | 0.003700 | -0.003200 |
| C | -1.874800 | -0.856400 | -0.023400 |
| S | -3.384200 | 0.155700 | -0.005700 |
| O | -4.505700 | -0.832200 | -0.013100 |
| O | -3.321400 | 0.967300 | 1.250200 |
| O | -3.330500 | 0.995700 | -1.243400 |
| H | 6.828800 | 0.731100 | -0.266700 |
| H | 6.238500 | -0.369600 | 0.986600 |
| H | 6.188900 | -0.856500 | -0.713500 |
| H | 4.561600 | 1.459300 | 0.506100 |
| H | 4.511900 | 0.976700 | -1.182200 |
| H | 3.630400 | -1.298000 | -0.488700 |
| H | 3.678400 | -0.790600 | 1.188500 |
| H | -0.576400 | 0.610800 | 0.902400 |
| H | -0.582700 | 0.661100 | -0.873100 |
| H | -1.931000 | -1.473200 | -0.924200 |
| H | -1.932000 | -1.510700 | 0.850400 |

| INT1 | | | |
|---|---|---|---|
| N | 6.927300 | -0.266400 | 0.178900 |
| C | 5.432600 | -0.462700 | 0.360900 |
| C | 4.552300 | 0.133400 | -0.740700 |
| O | 3.186300 | -0.155100 | -0.405600 |

| | | | |
|---|---|---|---|
| O | 2.817900 | -1.727200 | -2.420600 |
| P | 2.499500 | -1.563400 | -0.955800 |
| O | 0.927200 | -1.110300 | -0.742900 |
| O | -1.467200 | 3.052200 | -0.617700 |
| C | 0.342000 | -0.082400 | -1.568000 |
| C | -0.923400 | 2.067500 | -1.526200 |
| C | -0.386500 | 0.923100 | -0.682200 |
| C | 7.612500 | -0.819600 | 1.400500 |
| C | 7.429900 | -1.011400 | -1.028500 |
| C | 7.273300 | 1.192200 | 0.050800 |
| O | 2.731900 | -2.682400 | 0.025200 |
| O | -2.003300 | 4.349900 | -2.385100 |
| C | -1.992500 | 4.162000 | -1.178800 |
| C | -2.549100 | 5.098300 | -0.135400 |
| C | -3.114800 | 6.394400 | -0.716800 |
| C | -3.678000 | 7.312700 | 0.368300 |
| H | 5.269500 | -1.539600 | 0.429100 |
| H | 5.176200 | -0.004800 | 1.317900 |
| H | 4.637400 | 1.221000 | -0.776300 |
| H | 4.789300 | -0.272100 | -1.727300 |
| H | 1.115600 | 0.430200 | -2.145300 |
| H | -0.358800 | -0.556600 | -2.261600 |
| H | -1.711500 | 1.723000 | -2.201300 |
| H | -0.121600 | 2.520000 | -2.114700 |
| H | 0.313000 | 1.326600 | 0.053100 |
| H | 8.688300 | -0.704500 | 1.271900 |
| H | 7.272500 | -0.262900 | 2.273500 |
| H | 7.353200 | -1.873500 | 1.497700 |
| H | 7.155700 | -2.061300 | -0.927200 |
| H | 6.988900 | -0.587300 | -1.928700 |
| H | 8.513800 | -0.904800 | -1.064200 |
| H | 6.861700 | 1.582400 | -0.878300 |
| H | 6.859400 | 1.724000 | 0.907300 |
| H | 8.359200 | 1.283700 | 0.038100 |
| H | -3.317700 | 4.547800 | 0.421800 |
| H | -1.748900 | 5.304200 | 0.586200 |
| H | -2.326300 | 6.914700 | -1.272800 |
| H | -3.898300 | 6.151500 | -1.443500 |
| H | -4.071900 | 8.237600 | -0.065000 |
| H | -4.493300 | 6.827500 | 0.916800 |
| H | -2.906100 | 7.587700 | 1.095900 |

| | | | |
|---|---|---|---|
| C | 1.328600 | 1.460100 | 3.593300 |
| C | 0.744000 | 0.456900 | 2.595800 |
| C | -0.734100 | 0.757800 | 2.299600 |
| C | -1.440500 | -0.190300 | 1.296300 |
| S | -3.208800 | -0.323200 | 1.876700 |
| C | -3.967800 | -1.356500 | 0.558200 |
| C | -3.874800 | -2.854100 | 0.827700 |
| S | -4.650000 | -3.846100 | -0.479500 |
| O | -4.511200 | -5.265600 | -0.024500 |
| O | -6.077100 | -3.396000 | -0.557800 |
| O | -3.881800 | -3.557100 | -1.732300 |
| H | 2.373700 | 1.225500 | 3.820900 |
| H | 1.295100 | 2.481900 | 3.198200 |
| H | 0.771500 | 1.452300 | 4.537200 |
| H | 1.342800 | 0.464300 | 1.679400 |
| H | 0.823800 | -0.556500 | 3.003200 |
| H | -1.272400 | 0.697400 | 3.253000 |
| H | -0.860900 | 1.785300 | 1.943200 |
| H | -5.012900 | -1.042600 | 0.505100 |
| H | -3.488800 | -1.091900 | -0.385700 |
| H | -2.835200 | -3.179900 | 0.887700 |
| H | -4.379300 | -3.123600 | 1.759100 |
| O | -0.898800 | -1.491500 | 1.292000 |
| H | -0.136400 | -1.487200 | 0.673700 |
| O | -1.522300 | 0.320300 | -0.030300 |

| INT2 | | | |
|---|---|---|---|
| N | 6.776960 | 0.138483 | 0.234804 |
| C | 5.386211 | -0.469126 | 0.285463 |
| C | 4.370541 | 0.128922 | -0.691471 |
| O | 3.097706 | -0.481943 | -0.434163 |
| O | 3.061176 | -1.796446 | -2.658138 |
| P | 2.717749 | -1.915826 | -1.192391 |
| O | 1.095527 | -1.867743 | -0.951188 |
| O | -1.362061 | 2.281989 | -0.750903 |
| C | 0.278693 | -0.953952 | -1.703805 |
| C | -1.029497 | 1.171084 | -1.622620 |
| C | -0.272317 | 0.141187 | -0.795428 |
| C | 7.615890 | -0.567649 | 1.267077 |
| C | 7.407152 | -0.057398 | -1.116716 |
| C | 6.745182 | 1.606188 | 0.565552 |

| | | | |
|---|---|---|---|
| O | 3.220912 | -3.078973 | -0.375509 |
| O | -2.353736 | 3.333688 | -2.482275 |
| C | -2.059905 | 3.307765 | -1.300718 |
| C | -2.397989 | 4.358644 | -0.275129 |
| C | -3.196508 | 5.530232 | -0.847302 |
| C | -3.540308 | 6.568943 | 0.220753 |
| H | 5.510815 | -1.535786 | 0.093300 |
| H | 5.031877 | -0.333606 | 1.308801 |
| H | 4.235503 | 1.198899 | -0.523879 |
| H | 4.658205 | -0.033165 | -1.732952 |
| H | 0.849800 | -0.511378 | -2.524076 |
| H | -0.546602 | -1.531920 | -2.130820 |
| H | -1.950523 | 0.749151 | -2.032380 |
| H | -0.406621 | 1.540450 | -2.440581 |
| H | 0.564756 | 0.624757 | -0.288664 |
| H | 8.624576 | -0.156960 | 1.230331 |
| H | 7.172339 | -0.399564 | 2.248358 |
| H | 7.630396 | -1.632115 | 1.034405 |
| H | 7.398018 | -1.121286 | -1.353308 |
| H | 6.848237 | 0.504816 | -1.862819 |
| H | 8.431643 | 0.310798 | -1.069757 |
| H | 6.243739 | 2.149143 | -0.233723 |
| H | 6.217031 | 1.738701 | 1.509443 |
| H | 7.773664 | 1.954577 | 0.656434 |
| H | -2.948995 | 3.861685 | 0.533705 |
| H | -1.456011 | 4.699894 | 0.172244 |
| H | -2.620590 | 5.999776 | -1.652924 |
| H | -4.115762 | 5.148098 | -1.305388 |
| H | -4.111729 | 7.398161 | -0.208159 |
| H | -4.141763 | 6.129084 | 1.024329 |
| H | -2.635132 | 6.987165 | 0.675473 |
| C | 2.181405 | 1.489811 | 2.851191 |
| C | 0.754116 | 1.467873 | 2.299578 |
| C | 0.224317 | 0.030655 | 2.172093 |
| C | -1.177134 | -0.119251 | 1.527138 |
| S | -2.023129 | -1.516136 | 2.424202 |
| C | -3.660529 | -1.555087 | 1.589393 |
| C | -3.728111 | -2.510858 | 0.403778 |
| S | -5.366579 | -2.531071 | -0.376131 |
| O | -5.259116 | -3.528647 | -1.486171 |
| O | -6.331973 | -2.946991 | 0.691310 |

| | | | |
|---|---|---|---|
| O | -5.610154 | -1.136419 | -0.866680 |
| H | 2.544489 | 2.516044 | 2.969730 |
| H | 2.237312 | 1.000838 | 3.830549 |
| H | 2.870104 | 0.965697 | 2.178884 |
| H | 0.091239 | 2.034410 | 2.961921 |
| H | 0.728367 | 1.981925 | 1.332444 |
| H | 0.924458 | -0.593063 | 1.608161 |
| H | 0.182517 | -0.393223 | 3.182119 |
| H | -4.378814 | -1.850361 | 2.357633 |
| H | -3.890486 | -0.530987 | 1.289192 |
| H | -3.018394 | -2.225201 | -0.374061 |
| H | -3.518069 | -3.539989 | 0.706155 |
| O | -1.147723 | -0.503722 | 0.155067 |
| O | -1.988665 | 1.021324 | 1.702813 |
| H | -1.781625 | 1.641008 | 0.976719 |

| P | | | |
|---|---|---|---|
| C | -3.643400 | 6.401000 | 0.229100 |
| C | -3.813900 | 4.981100 | -0.312300 |
| C | -2.887000 | 2.614100 | -0.268400 |
| O | -3.597200 | 2.260900 | -1.194700 |
| C | -2.801800 | 4.009600 | 0.295800 |
| N | 6.248200 | -0.868200 | -0.004900 |
| C | 4.829700 | -0.630800 | -0.491900 |
| C | 3.730800 | -0.767800 | 0.565800 |
| O | 2.477100 | -0.515100 | -0.086300 |
| O | 2.554500 | 1.816700 | -1.246800 |
| P | 1.903600 | 1.045000 | -0.126300 |
| O | 0.391300 | 0.691900 | -0.650500 |
| O | -2.965200 | -1.726600 | -0.072100 |
| C | -0.511900 | -0.038200 | 0.194700 |
| C | -2.984800 | -0.438500 | 0.582000 |
| C | -1.926600 | 0.415000 | -0.111800 |
| C | 7.163700 | -0.733300 | -1.193300 |
| C | 6.646200 | 0.154600 | 1.025500 |
| C | 6.402500 | -2.249500 | 0.570600 |
| O | 1.892300 | 1.596800 | 1.279200 |
| O | -2.042800 | 1.773800 | 0.380400 |
| O | -4.612100 | -2.425100 | 1.304400 |
| C | -3.843100 | -2.648300 | 0.384100 |
| C | -3.738000 | -3.936600 | -0.392300 |

| | | | |
|---|---|---|---|
| C | -4.688600 | -5.028500 | 0.098900 |
| C | -4.565700 | -6.309200 | -0.727500 |
| H | -4.375900 | 7.081700 | -0.216500 |
| H | -3.777400 | 6.432800 | 1.316300 |
| H | -2.644600 | 6.792600 | 0.005700 |
| H | -4.829200 | 4.622300 | -0.107000 |
| H | -3.699800 | 4.983900 | -1.402100 |
| H | -1.774400 | 4.358200 | 0.127400 |
| H | -2.917400 | 3.940100 | 1.384000 |
| H | 4.813100 | 0.376400 | -0.910200 |
| H | 4.655000 | -1.350500 | -1.293400 |
| H | 3.676600 | -1.784300 | 0.960200 |
| H | 3.868300 | -0.074800 | 1.400000 |
| H | -0.421200 | -1.109100 | -0.011100 |
| H | -0.285300 | 0.142200 | 1.250300 |
| H | -3.971700 | 0.014800 | 0.467500 |
| H | -2.767900 | -0.559000 | 1.647100 |
| H | -2.101900 | 0.411700 | -1.189900 |
| H | 8.188000 | -0.893200 | -0.857900 |
| H | 6.885500 | -1.483100 | -1.933400 |
| H | 7.050900 | 0.268900 | -1.606300 |
| H | 7.701100 | 0.010200 | 1.257100 |
| H | 6.482300 | 1.148500 | 0.609700 |
| H | 6.050200 | 0.013100 | 1.925300 |
| H | 5.804900 | -2.331700 | 1.476600 |
| H | 6.072900 | -2.974900 | -0.173000 |
| H | 7.455200 | -2.402500 | 0.807500 |
| H | -3.925800 | -3.697400 | -1.446800 |
| H | -2.693700 | -4.268900 | -0.344600 |
| H | -4.479100 | -5.243400 | 1.153200 |
| H | -5.718300 | -4.656000 | 0.056600 |
| H | -5.256200 | -7.077600 | -0.365500 |
| H | -4.795100 | -6.124500 | -1.783200 |
| H | -3.551300 | -6.720700 | -0.675900 |

| PB1 | | | |
|---|---|---|---|
| S | -3.029700 | 0.210400 | 0.000000 |
| C | -1.326500 | -0.516500 | 0.000000 |
| C | -0.307700 | 0.614600 | 0.000000 |
| S | 1.401200 | -0.000400 | 0.000000 |
| O | 2.250700 | 1.230500 | 0.000000 |

| | | | |
|---|---|---|---|
| O | 1.548700 | -0.815600 | 1.246600 |
| O | 1.548700 | -0.815600 | -1.246600 |
| H | -3.699400 | -0.960700 | 0.000000 |
| H | -1.206200 | -1.136700 | 0.889400 |
| H | -1.206200 | -1.136700 | -0.889400 |
| H | -0.406300 | 1.245500 | -0.887400 |
| H | -0.406300 | 1.245600 | 0.887400 |

| C | | | |
|---|---|---|---|
| C | -5.335000 | -0.809000 | 0.364000 |
| C | -3.842000 | -0.980900 | 0.082800 |
| C | -3.118700 | 0.367600 | -0.012800 |
| C | -1.643000 | 0.232000 | -0.311400 |
| S | -0.660000 | 1.658900 | 0.208100 |
| O | -1.136700 | -0.724500 | -0.870300 |
| C | 0.990400 | 1.149000 | -0.407600 |
| C | 1.626000 | 0.167600 | 0.573500 |
| N | 2.952900 | -0.426500 | 0.114900 |
| C | 3.452100 | -1.316700 | 1.222600 |
| C | 2.775500 | -1.259300 | -1.125300 |
| C | 3.970500 | 0.650200 | -0.142200 |
| H | -5.833500 | -1.781300 | 0.432200 |
| H | -5.504300 | -0.281000 | 1.309000 |
| H | -5.826200 | -0.236300 | -0.430600 |
| H | -3.379400 | -1.580700 | 0.875300 |
| H | -3.697600 | -1.532300 | -0.852000 |
| H | -3.545400 | 0.971300 | -0.826700 |
| H | -3.252300 | 0.954000 | 0.903300 |
| H | 0.845400 | 0.719100 | -1.399100 |
| H | 1.563200 | 2.073000 | -0.497700 |
| H | 1.829600 | 0.659000 | 1.526400 |
| H | 0.967600 | -0.684500 | 0.744200 |
| H | 4.401900 | -1.751400 | 0.913100 |
| H | 3.585600 | -0.713600 | 2.120100 |
| H | 2.715900 | -2.100900 | 1.397700 |
| H | 3.722300 | -1.756200 | -1.336200 |
| H | 1.992100 | -1.994400 | -0.943100 |
| H | 2.507200 | -0.615800 | -1.960900 |
| H | 3.652000 | 1.254100 | -0.990400 |
| H | 4.061500 | 1.264700 | 0.753400 |
| H | 4.922400 | 0.170700 | -0.370300 |

| INT3 | | | |
|---|---|---|---|
| C | -5.508662 | -0.605902 | 3.314422 |
| C | -4.433354 | -0.550474 | 2.226523 |
| C | -3.327266 | 0.456335 | 2.572861 |
| C | -2.233426 | 0.601042 | 1.503281 |
| S | -0.972136 | 1.861954 | 2.179036 |
| C | -0.846410 | 3.186170 | 0.911606 |
| C | 0.175831 | 2.822623 | -0.159119 |
| N | 0.517718 | 3.941388 | -1.143076 |
| C | 1.500314 | 3.377886 | -2.136431 |
| C | -0.706716 | 4.408613 | -1.877894 |
| C | 1.162298 | 5.106735 | -0.443106 |
| H | -6.296412 | -1.320984 | 3.055506 |
| H | -5.083926 | -0.912274 | 4.277198 |
| H | -5.981039 | 0.372674 | 3.457416 |
| H | -3.997487 | -1.545130 | 2.084792 |
| H | -4.895888 | -0.268865 | 1.272760 |
| H | -3.762860 | 1.448743 | 2.724380 |
| H | -2.848528 | 0.170225 | 3.516979 |
| H | -1.836784 | 3.369970 | 0.497305 |
| H | -0.542354 | 4.066462 | 1.482373 |
| H | 1.124369 | 2.531758 | 0.292538 |
| H | -0.191694 | 2.000222 | -0.772795 |
| H | 1.762902 | 4.164479 | -2.843433 |
| H | 2.384014 | 3.040857 | -1.595296 |
| H | 1.034506 | 2.540572 | -2.654421 |
| H | -0.396943 | 5.133112 | -2.631164 |
| H | -1.177296 | 3.547544 | -2.352493 |
| H | -1.393024 | 4.877462 | -1.174530 |
| H | 0.446227 | 5.564552 | 0.236844 |
| H | 2.031543 | 4.743087 | 0.104806 |
| H | 1.466974 | 5.829392 | -1.199727 |
| N | 6.566301 | -1.447018 | 0.027311 |
| C | 5.543122 | -2.540069 | 0.318094 |
| C | 4.130849 | -2.105932 | 0.661786 |
| O | 3.414175 | -1.665582 | -0.504263 |
| O | 3.245158 | 0.836392 | 0.148305 |
| P | 2.449967 | -0.330876 | -0.390810 |
| O | 1.421864 | -0.749244 | 0.828414 |
| O | -3.006252 | -1.368001 | -1.014619 |

| | | | |
|---|---|---|---|
| C | 0.391243 | -1.715975 | 0.562599 |
| C | -1.792473 | -2.042548 | -0.596744 |
| C | -0.932892 | -1.050475 | 0.180508 |
| C | 7.895973 | -2.127083 | -0.162668 |
| C | 6.236003 | -0.685252 | -1.228776 |
| C | 6.678227 | -0.481025 | 1.176395 |
| O | 1.726456 | -0.249687 | -1.711698 |
| O | -2.710950 | 1.107710 | 0.289451 |
| O | -4.075987 | -3.350312 | -1.194845 |
| C | -4.085611 | -2.136915 | -1.303689 |
| C | -5.240011 | -1.305331 | -1.801598 |
| C | -6.587580 | -2.025498 | -1.716078 |
| C | -7.726722 | -1.189147 | -2.300524 |
| H | 5.537145 | -3.191317 | -0.558014 |
| H | 5.944308 | -3.097418 | 1.167242 |
| H | 3.630348 | -2.995514 | 1.058889 |
| H | 4.120204 | -1.344700 | 1.444901 |
| H | 0.707484 | -2.401284 | -0.229330 |
| H | 0.252999 | -2.296023 | 1.478686 |
| H | -2.050740 | -2.904136 | 0.021627 |
| H | -1.266953 | -2.383200 | -1.492573 |
| H | -0.732124 | -0.173206 | -0.436051 |
| H | 8.638941 | -1.367799 | -0.404778 |
| H | 8.165108 | -2.637842 | 0.761916 |
| H | 7.807826 | -2.843096 | -0.979576 |
| H | 6.111432 | -1.395504 | -2.045813 |
| H | 5.323143 | -0.117562 | -1.068757 |
| H | 7.065640 | -0.010169 | -1.439471 |
| H | 5.766111 | 0.109177 | 1.242935 |
| H | 6.840614 | -1.046659 | 2.093831 |
| H | 7.526023 | 0.175717 | 0.982971 |
| H | -5.253553 | -0.358004 | -1.251968 |
| H | -5.007771 | -1.048273 | -2.844735 |
| H | -6.517301 | -2.981696 | -2.245646 |
| H | -6.800569 | -2.265370 | -0.667618 |
| H | -8.681312 | -1.718837 | -2.220730 |
| H | -7.830531 | -0.233122 | -1.775179 |
| H | -7.554007 | -0.969308 | -3.360147 |
| O | -1.637937 | -0.674582 | 1.381458 |
| H | -3.140861 | 0.372362 | -0.190037 |

| INT4 | | | |
|---|---|---|---|
| C | -3.900500 | -0.244100 | 4.337000 |
| C | -2.842600 | -0.457400 | 3.252600 |
| C | -3.045400 | 0.489400 | 2.061200 |
| C | -1.925000 | 0.433800 | 1.018300 |
| S | -2.524400 | 1.499700 | -0.466600 |
| C | -1.038500 | 2.468100 | -0.939700 |
| C | -0.955800 | 3.748600 | -0.114400 |
| N | 0.231400 | 4.654600 | -0.441400 |
| C | 0.208100 | 5.800700 | 0.534100 |
| C | 1.530900 | 3.913200 | -0.287700 |
| C | 0.129300 | 5.204800 | -1.836500 |
| H | -3.752400 | -0.935700 | 5.173100 |
| H | -3.860700 | 0.775600 | 4.736800 |
| H | -4.911100 | -0.407600 | 3.945600 |
| H | -1.843800 | -0.298500 | 3.675500 |
| H | -2.877900 | -1.494600 | 2.902800 |
| H | -3.993600 | 0.259000 | 1.561600 |
| H | -3.108600 | 1.524700 | 2.410500 |
| H | -0.156400 | 1.840900 | -0.820100 |
| H | -1.165300 | 2.677100 | -2.004300 |
| H | -1.845800 | 4.362800 | -0.260900 |
| H | -0.858000 | 3.505400 | 0.943500 |
| H | 1.029200 | 6.475600 | 0.293400 |
| H | -0.746500 | 6.318600 | 0.441900 |
| H | 0.328500 | 5.402800 | 1.541500 |
| H | 2.344200 | 4.634500 | -0.369200 |
| H | 1.546500 | 3.434200 | 0.690100 |
| H | 1.618700 | 3.170700 | -1.078200 |
| H | 0.168700 | 4.385600 | -2.552300 |
| H | -0.810900 | 5.748100 | -1.929100 |
| H | 0.971400 | 5.877400 | -1.999100 |
| N | 6.464300 | -1.706500 | 0.023200 |
| C | 5.284500 | -2.626000 | 0.326200 |
| C | 3.950600 | -1.968600 | 0.616900 |
| O | 3.400800 | -1.347100 | -0.557500 |
| O | 3.032300 | 0.979100 | 0.546600 |
| P | 2.416000 | -0.040800 | -0.376600 |
| O | 1.163700 | -0.691800 | 0.506400 |
| O | -3.014200 | -2.087200 | -1.628200 |
| C | 0.392400 | -1.786900 | -0.032900 |

| | | | |
|---|---|---|---|
| C | -1.723900 | -2.501200 | -1.131900 |
| C | -0.995200 | -1.330700 | -0.476500 |
| C | 7.713800 | -2.542600 | 0.105900 |
| C | 6.370300 | -1.134200 | -1.367300 |
| C | 6.567200 | -0.585100 | 1.019400 |
| O | 1.956100 | 0.312300 | -1.766300 |
| O | -1.795400 | -0.918700 | 0.645100 |
| O | -4.074100 | -2.960500 | 0.181400 |
| C | -4.110900 | -2.365600 | -0.882700 |
| C | -5.361400 | -1.852500 | -1.553500 |
| C | -6.650300 | -2.296900 | -0.861000 |
| C | -7.898800 | -1.764300 | -1.566000 |
| H | 5.209400 | -3.309600 | -0.521500 |
| H | 5.567700 | -3.195500 | 1.213900 |
| H | 3.281600 | -2.772600 | 0.943500 |
| H | 4.028000 | -1.247300 | 1.433500 |
| H | 0.916000 | -2.240500 | -0.878600 |
| H | 0.303700 | -2.531900 | 0.762500 |
| H | -1.847900 | -3.326200 | -0.428400 |
| H | -1.165800 | -2.842800 | -2.005300 |
| H | -0.905800 | -0.508600 | -1.192900 |
| H | 8.562900 | -1.923100 | -0.181300 |
| H | 7.831700 | -2.896400 | 1.129900 |
| H | 7.613600 | -3.384600 | -0.579000 |
| H | 6.342700 | -1.959900 | -2.078000 |
| H | 5.466900 | -0.535600 | -1.444800 |
| H | 7.254400 | -0.519800 | -1.537500 |
| H | 5.729100 | 0.097200 | 0.886700 |
| H | 6.563300 | -1.006900 | 2.024600 |
| H | 7.504100 | -0.058400 | 0.837700 |
| H | -5.288800 | -0.757300 | -1.580500 |
| H | -5.339000 | -2.178400 | -2.599900 |
| H | -6.680500 | -3.392200 | -0.827000 |
| H | -6.636000 | -1.955800 | 0.180200 |
| H | -8.808600 | -2.094300 | -1.054500 |
| H | -7.906900 | -0.668600 | -1.587300 |
| H | -7.952200 | -2.116900 | -2.602200 |
| O | -0.751000 | 0.958400 | 1.560600 |
| H | 0.022700 | 0.479200 | 1.188000 |

| PC1 |
|---|

| | | | |
|---|---|---|---|
| S | -2.871000 | 0.129000 | -0.097800 |
| C | -1.180200 | -0.570500 | 0.110700 |
| C | -0.128800 | 0.512900 | -0.104600 |
| N | 1.319600 | 0.038100 | -0.007300 |
| C | 2.200000 | 1.241600 | -0.215300 |
| C | 1.612700 | -0.544500 | 1.347800 |
| C | 1.633400 | -0.975300 | -1.073200 |
| H | -2.852900 | 0.930400 | 0.988300 |
| H | -1.110800 | -1.029200 | 1.096900 |
| H | -1.113700 | -1.356400 | -0.643800 |
| H | -0.230700 | 0.953800 | -1.097300 |
| H | -0.229000 | 1.300300 | 0.645100 |
| H | 3.239900 | 0.920400 | -0.162300 |
| H | 1.984200 | 1.665100 | -1.196000 |
| H | 1.987600 | 1.968700 | 0.568100 |
| H | 1.059400 | -1.473900 | 1.470200 |
| H | 2.682500 | -0.743200 | 1.407200 |
| H | 1.320300 | 0.179600 | 2.108000 |
| H | 1.384100 | -0.547900 | -2.044300 |
| H | 2.698000 | -1.202500 | -1.022200 |
| H | 1.057700 | -1.881600 | -0.894400 |

**11. Full reference list**

(20) Gaussian 09, Revision A.02, M. J. Frisch, G. W. Trucks, H. B. Schlegel, G. E. Scuseria, M. A. Robb, J. R. Cheeseman, G. Scalmani, V. Barone, G. A. Petersson, H. Nakatsuji, X. Li, M. Caricato, A. Marenich, J. Bloino, B. G. Janesko, R. Gomperts, B. Mennucci, H. P. Hratchian, J. V. Ortiz, A. F. Izmaylov, J. L. Sonnenberg, D. Williams-Young, F. Ding, F. Lipparini, F. Egidi, J. Goings, B. Peng, A. Petrone, T. Henderson, D. Ranasinghe, V. G. Zakrzewski, J. Gao, N. Rega, G. Zheng, W. Liang, M. Hada, M. Ehara, K. Toyota, R. Fukuda, J. Hasegawa, M. Ishida, T. Nakajima, Y. Honda, O. Kitao, H. Nakai, T. Vreven, K. Throssell, J. A. Montgomery, Jr., J. E. Peralta, F. Ogliaro, M. Bearpark, J. J. Heyd, E. Brothers, K. N. Kudin, V. N. Staroverov, T. Keith, R. Kobayashi, J. Normand, K. Raghavachari, A. Rendell, J. C. Burant, S. S. Iyengar, J. Tomasi, M. Cossi, J. M. Millam, M. Klene, C. Adamo, R. Cammi, J. W. Ochterski, R. L. Martin, K. Morokuma, O. Farkas, J. B. Foresman, and D. J. Fox, Gaussian, Inc., Wallingford CT, 2016.
